# Expansion of human hematopoietic stem cells by inhibiting translation

**DOI:** 10.1101/2023.11.28.568925

**Authors:** Chenchen Li, Hanna Shin, Dheeraj Bhavanasi, Mai Liu, Maria I. Briones, Xiang Yu, Xiaolei Liu, Friederike Herbst, Hooda Said, Mohamad-Gabriel Alameh, Laura Breda, Narasaiah Kovuru, Yeipyeng Kwa, Nithin Sam Ravi, David C. Schultz, Sara Cherry, Stefano Rivella, Juan R. Alvarez-Dominguez, Gerd A. Blobel, Brian D. Gregory, Scott A. Peslak, Jian Huang, Peter S. Klein

## Abstract

Hematopoietic stem cell (HSC) transplantation using umbilical cord blood (UCB) is a potentially life-saving treatment for leukemia and bone marrow failure but is limited by the low number of HSCs in UCB. The loss of HSCs after ex vivo manipulation is also a major obstacle to gene editing for inherited blood disorders. HSCs require a low rate of translation to maintain their capacity for self-renewal, but hematopoietic cytokines used to expand HSCs stimulate protein synthesis and impair long-term self-renewal. We previously described cytokine-free conditions that maintain but do not expand human and mouse HSCs ex vivo. Here we performed a high-throughput screen and identified translation inhibitors that allow ex vivo expansion of human HSCs while minimizing cytokine exposure. Transplantation assays show a ∼4.3-fold expansion of long-term HSCs from UCB after one week of culture in low cytokine conditions that include the translation initiation inhibitor 4E1RCat. Single cell transcriptomic analysis demonstrates maintenance of the HSC/MPP transcriptional signature and reduced expression of stress markers under culture conditions that maintain or expand functional long-term HSCs. Addition of 4E1RCat induces a dose-dependent increase in autophagic vacuoles and reduces expression of oxidative phosphorylation and cell cycle gene expression programs. These culture conditions maintain adult HSCs after CRISPR/Cas9 editing of the *BCL11A*+58 enhancer, overcoming a major obstacle to ex vivo gene correction for human hemoglobinopathies.

**Significance:**

- We report ex vivo expansion of human hematopoietic stem cells from umbilical cord blood using low cytokines and small molecule inhibitors of GSK-3, mTORC1, and the translation initiation factor eIF4E.
- Our method maintains human HSCs ex vivo after gene editing with conventional CRISPR/Cas9, lipid nanoparticle delivery, and base editing approaches, overcoming a major obstacle to gene editing for inherited blood disorders.

## Introduction

Hematopoietic stem cells (HSCs) are primarily quiescent but on demand are capable of expansion and differentiation into multiple lineages. Quiescent HSCs maintain a low rate of protein synthesis and mitochondrial activity compared to more differentiated hematopoietic cells, whereas activation of HSCs is associated with increased translation, a shift from glycosis to oxidative phosphorylation, reduced autophagy, and ultimately restriction of self-renewal capacity^1–7^. HSCs survive poorly outside of the complex hematopoietic niche and traditionally have been supported in culture by the addition of multiple hematopoietic cytokines, which promote survival but also drive proliferation, lineage commitment, and the loss of self-renewing, long-term HSCs (LT-HSCs)^3,8,9^. The loss of long-term repopulating capacity in cultured human HSCs appears to occur before cell division^10^, suggesting that factors aside from cell cycle entry contribute to impaired HSC function, including reduced autophagy, increased oxidative phosphorylation, and increased protein synthesis; thus interventions that mitigate these changes may improve maintenance and expansion of long-term HSCs in culture^2–6,11,12^.

HSC transplantation (HSCT) is a life-saving therapy for hematological neoplasms and bone marrow failure but is constrained by the limited availability of suitably matched donors, especially for ethnic groups that are underrepresented in bone marrow registries^13,14^. While haploidentical transplants help to address this issue^15,16^, umbilical cord blood (UCB) remains a valuable and underutilized^17^ resource for human HSCs that requires reduced stringency in HLA matching and has a reduced risk of graft-versus-host disease (GVHD)^17–24^. UCB units, however, typically contain a low number of HSCs, resulting in delayed neutrophil and platelet engraftment. The use of two UCB donors can improve the rate of neutrophil engraftment, but is associated with immunological extinction of one of the two donors, increased GVHD, impaired platelet recovery^25^, and substantially increased cost. Thus, even modest expansion of HSCs in existing UCB units would dramatically increase the number of UCB units available for genetically diverse patients needing HSCT.

Approaches to expand HSCs in UCB have relied on cocktails of hematopoietic cytokines combined with an array of factors, including the pyrimidoindole derivative UM171, nicotinamide, the aryl hydrocarbon receptor antagonist SR1, the PPARγ antagonist GW9662, zwitterionic hydrogels, polyvinyl alcohol (PVA), histone deacetylase inhibitors, BET inhibitors, Notch ligands, angiopoietin-like proteins, pleiotrophin, small molecule cytokine-mimetics, a ferroptosis inhibitor, mitofuscin agonists, and others^26–47^. A subset of these approaches was validated with limiting dilution transplant assays and serial transplantation in immunocompromised mice. A formulation using CD133^+^ cells isolated from UCB, cultured with nicotinamide and multiple cytokines, and then combined with the CD133^-^ fraction (termed Omidubicel) showed improved neutrophil and platelet recovery compared to single-or double-unit UCB transplants in a Phase 3 trial^40^ and has been FDA-approved for use in hematopoietic malignancies. SR1 and UM171 have also shown promising data in early phase clinical trials^48–50^. However, all of these HSC expansion conditions depend on high concentrations of multiple cytokines or small molecule activators of cytokine signaling, incurring the potential for increased protein synthesis and mitochondrial activity and driving cells into lineage commitment^27,28,39,42–44^.

To circumvent the loss of self-renewal associated with cytokine activation, we have explored cytokine-free conditions for the ex vivo maintenance of HSCs. We showed that human and mouse LT-HSCs can be maintained ex vivo without cytokines, support cells, or serum when the signaling kinases glycogen synthase kinase 3 (GSK-3) and mechanistic target of rapamycin complex 1 (mTORC1) are inhibited^12,51,52^. Mechanistically, GSK-3 inhibition activates Wnt/β-catenin signaling, which is required for ex vivo HSC maintenance^51^. However, GSK-3 inhibition also activates mTORC1, leading to HSC exhaustion^53^. Adding an mTORC1 inhibitor to the GSK-3 inhibitor therefore allows Wnt-mediated HSC maintenance while preventing HSC exhaustion ^51,52^. In vivo limiting dilution assays showed no loss of long-term reconstituting activity in these cytokine-free conditions. However, the absolute number of LT-HSCs was not substantially increased. To identify an approach to expand LT-HSCs ex vivo, we performed a high-throughput chemical screen based on our previously described HSC culture conditions^51^ modified to include a mitogenic stimulus while minimizing cytokine exposure. The screen identified multiple inhibitors of translation initiation, consistent with published evidence that HSCs restrict translation to maintain long-term self-renewal^1,2,11,54–59^. However, pharmacological inhibition of translation has not previously been tested for therapeutic expansion of human HSCs. Here we show that pharmacological inhibition of translation paired with limited exposure to hematopoietic cytokines yields ex vivo expansion of human LT-HSCs.

## Methods

Human CD34^+^ cells from umbilical cord blood was obtained from the StemCell Technologies and cultured in StemSpan™ SFEM II medium with vehicle (DMSO), 3 µM CHIR99021 and 5 nM rapamycin (designated CR), CR with 12.5 ng/ml human stem cell factor (SCF), 12.5 ng/ml human Thrombopoietin (TPO), and 1.25 ng/ml human IL3 (designated CRCY), or CRCY with 2µM 4E1RCat (designted CRCE) and cultured for 2 or 7 days as described in detail in Supplemental Methods. Other additives and culture conditions are also described in greater detail in indicated. Single cell RNA-sequencing was performed on CD34+CD38-CD45RA-CD90+ cells at Day 0 (uncultured) and after 2 days of culture in DMSO, CR, CRCY, or CRCE as described in the text and supplemental methods. Limiting dilution assays, serial transplants, flow cytometry, autophagy assay, and gene editing were performed as described in the Supplemental Methods.

## Results

### Maintenance of an HSC signature ex vivo

LT-HSCs can be maintained in cytokine-free medium for at least 7 days by inhibiting GSK-3 and mTORC1^12,51^ but undergo limited cell division in the absence of mitogenic stimuli. To achieve a modest expansion of HSCs, we tested multiple combinations of common hematopoietic cytokines to find conditions that would minimize cytokine exposure and still allow expansion of CD34^+^ cells in the presence of GSK-3 and mTORC1 inhibitors (CHIR99021 and rapamycin; CR). We found that limiting the cocktail to 3 cytokines (stem cell factor (SCF), thrombopoietin (TPO), and interleukin-3 (IL-3)) was sufficient to achieve expansion in the presence of CR (Supplemental Figure 1A). Most HSC expansion methods use cytokines such as SCF and TPO at substantially higher concentrations^26–41^. We therefore tested different concentrations of SCF, TPO, and IL-3 (STI) to identify low cytokine concentrations that would still induce detectable proliferation in CR-containing medium (Supplemental Figure 1B). Robust expansion of CD34^+^ cells was detected with SCF and TPO at 12.5 ng/mL and IL-3 at 1.25 ng/mL (Supplemental Figure 1C) in CR medium, and these low cytokine conditions (CRCY) were selected for further study.

To examine in more detail how subpopulations of hematopoietic cells are maintained upon culture in CR, we performed single-cell RNA sequencing (scRNA-seq). As CD34^+^ cells from human UCB are heterogeneous, and only a small fraction are functional LT-HSCs, we enriched for HSCs using fluorescence-activated cell sorting (FACS) of CD34^+^CD38^-^CD45RA^-^CD90^+^ cells^64^. Sorted cells were then analyzed immediately (uncultured control/Day 0) or cultured for two days in STEM-Span medium with vehicle (DMSO), CR, or CRCY, followed by scRNA-seq.

We chose 2 days of culture to identify early transcriptional responses to our culture conditions and because CD34^+^ cells cultured in STEM-Span alone (DMSO) do not survive well in longer term culture, in stark contrast to cells cultured in CR, which remain viable at day 7 and maintain 100% of their LT-HSCs^51^. Analysis of single cell data using Seurat identified 10 groups of cells, with several groups showing substantial overlap in gene expression. These groups were then annotated manually and each group was assigned a label based on their distinct markers yielding five main groups: hematopoietic stem cells and multipotent progenitor cells (HSCs/MPPs), common myeloid progenitors (CMP), lymphoid-primed multipotent progenitors (LMPP), megakaryocyte-erythrocyte progenitors (MEP), and early erythroid commitment (EEC). Uniform manifold approximation and projection (UMAP) visualization showed that culture in CR or CRCY maintains the HSC/MPP population present in freshly isolated cells (Day 0, Figure 1A and 1B), whereas this population declines in cells cultured in STEM-span alone (DMSO, Figure 1A and 1B). These single cell findings are consistent with our prior functional assays showing maintenance of HSCs cultured with CR^12,51^.

**Figure 1.**
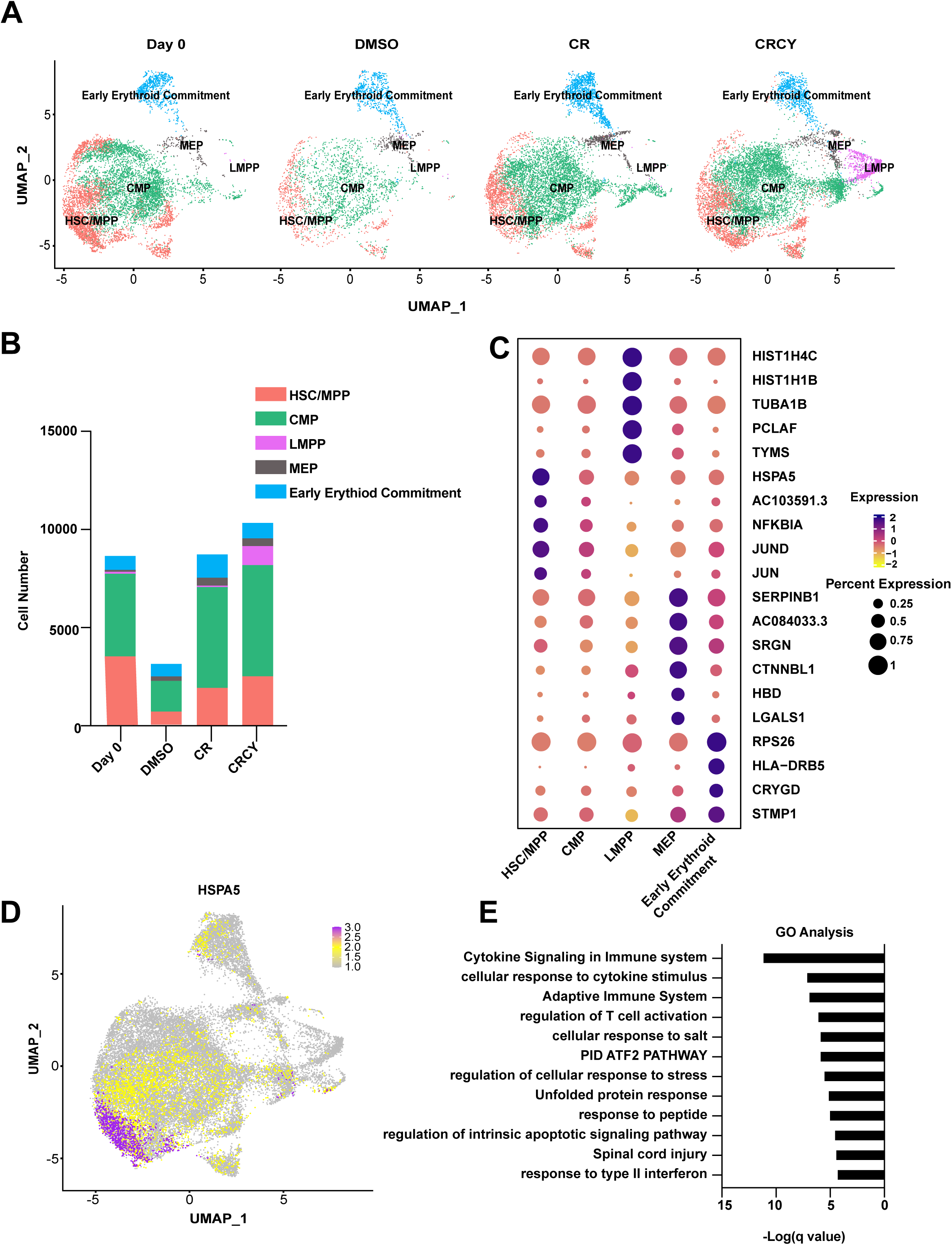
Maintenance of HSC/MPP gene signature in low cytokine culture conditions. **A.** HSCs were enriched from UCB CD34^+^ cells by purifying CD34^+^CD38^-^CD45RA^-^CD90^+^ cells, which contain ∼5-10% functional HSCs^64,104,105^, and were then cultured in cytokine-free medium with CHIR99021 and rapamycin (CR), CR with low-dose cytokines (CRCY), or control medium (DMSO) for 2 days and subjected to single cell RNA-seq. Freshly sorted cells (Day 0) were isolated at the same time. HSC/MPP, CMP, MEP, LMPP and early erythroid commitment populations were identified based on previously published data^60^ and visualized by uniform manifold approximation and projection (UMAP). Each dot represents one cell and colors represent cell clusters as indicated. **B.** Number of cells in HSC/MPP, CMP, MEP, LMPP, and early erythroid commitment populations at Day 0 or after culture in control (DMSO), CR, or CRCY media. **C.** Scorecard dot plot showing top 5 enriched genes within HSC, CMP, MEP, LMPP and early erythroid commitment populations. Diameter of circle represents percent of cells expressing each marker and color indicates relative expression in the respective populations. **D.** UMAP feature plot showing enrichment of HSPA5/GRP78 in the HSPC population. **E.** Gene Ontology (GO) enrichment analysis of HSCs signature genes compared to other populations. Bar graph shows significantly enriched pathways, with Fisher’s exact test-log [q value] on X-axis. GEO SubSeries Accession number GSE248311.

Although there was overlap between the HSC/MPP and CMP groups in CD34^+^CD38^-^CD45RA^-^ CD90^+^ cells, the HSC/MPP group was distinguished by the significantly higher expression of the unfolded protein response (UPR) chaperone *HSPA5* and several immediate early response genes (IEGs) including *IER2, NFKBIA,* and JUN family transcription factors, which are associated with stress response through Jun kinases (Figure 1C, Supplemental Table 1). Indeed, *HSPA5* (Figure 1D) and JUN family members were among the most highly enriched genes in the HSC/MPP population compared to other hematopoietic subpopulations. Gene Ontology (GO) analysis of the entire list of genes associated with the HSC/MPP population in CR-containing medium also showed significant enrichment of genes related to the regulation of cellular response to stress and the unfolded protein response pathways (Figure 1E, Supplemental Table 2). A primary output of the unfolded protein response is to suppress translation^65^. The high expression of *HSPA5* and stress response genes in the HSC/MPP population compared to other cell types is therefore consistent with the critical role of limited translation and proteostasis in HSC function^1,2,54,55^. While expression of *HSPA5*, other stress response genes, and the IEGs *IER2, NFKBIA*, and *JUN*-related genes is enriched in HSC/MPPs compared to other cell types, the expression of these genes was also very high in day 0 cells compared to cells in culture (Supplemental Figures 2 and 3), as expected for cells that were recently thawed and subjected to cell sorting^66–68^. However, CRCY also reduced expression of most UPR markers and IEGs compared to cells cultured in medium with vehicle (DMSO), indicating an effect of CRCY independent of cell culture (Supplemental Figure 2).

### High-throughout screen with low cytokines

We performed a high-throughput screen (HTS) of 2,240 FDA-approved and/or bioactive compounds to identify drugs that would enhance expansion of HSCs from UCB in CRCY (Figure 2A, Supplemental Figure 1D). Initially we screened for compounds that increased HSPCs (CD34^+^ cells) in the presence of GSK-3 and mTORC inhibitors and low dose cytokines (CRCY). We identified 74 compounds that increase the number of CD34^+^ cells > 1.4-fold and > 4 standard deviation units above the mean of control cells in CRCY alone in 2 replicate screens (Figure 2B, Supplemental Table 3) and 60 of these compounds were detected in a third replicate at 10-fold lower drug concentrations. Among these 60 hits, two compounds that directly inhibit the cap-dependent translation initiation factor eIF4E (4E1RCat^69^ and 4EGI-1^70^) drew our attention because of prior work showing that suppression of cap-dependent translation is essential to maintain long-term HSCs in mice^2,11,55,56^. 4E1RCat is a rhodanine-based compound that inhibits eIF4E interaction with the eIF4G scaffold and 4EGI-1 is a thiazole derivative that also inhibits eIF4E assembly into the eIF4F complex. To validate these translation inhibitors further, we cultured human CD34^+^ cells from UCB in CRCY with 4E1RCat, the structurally related rhodanine-derivative 4E2RCat, or with 4EGI-1 for 7 days and then performed flow cytometry to detect pHSCs. pHSCs cultured in CRCY with 4E1RCat, 4E2RCat, or 4EGI-1 expanded up to ∼10-fold compared to the number of pHSCs at Day 0 (Figure 2C), demonstrating dose-dependent expansion in the presence of eIF4E inhibitors. pHSC expansion was also significantly higher in CRCY+4E1RCat (hereafter referred to as CRCE) compared to CRCY alone. Therefore, we focused on 4E1RCat for functional studies of HSC expansion.

**Figure 2.**
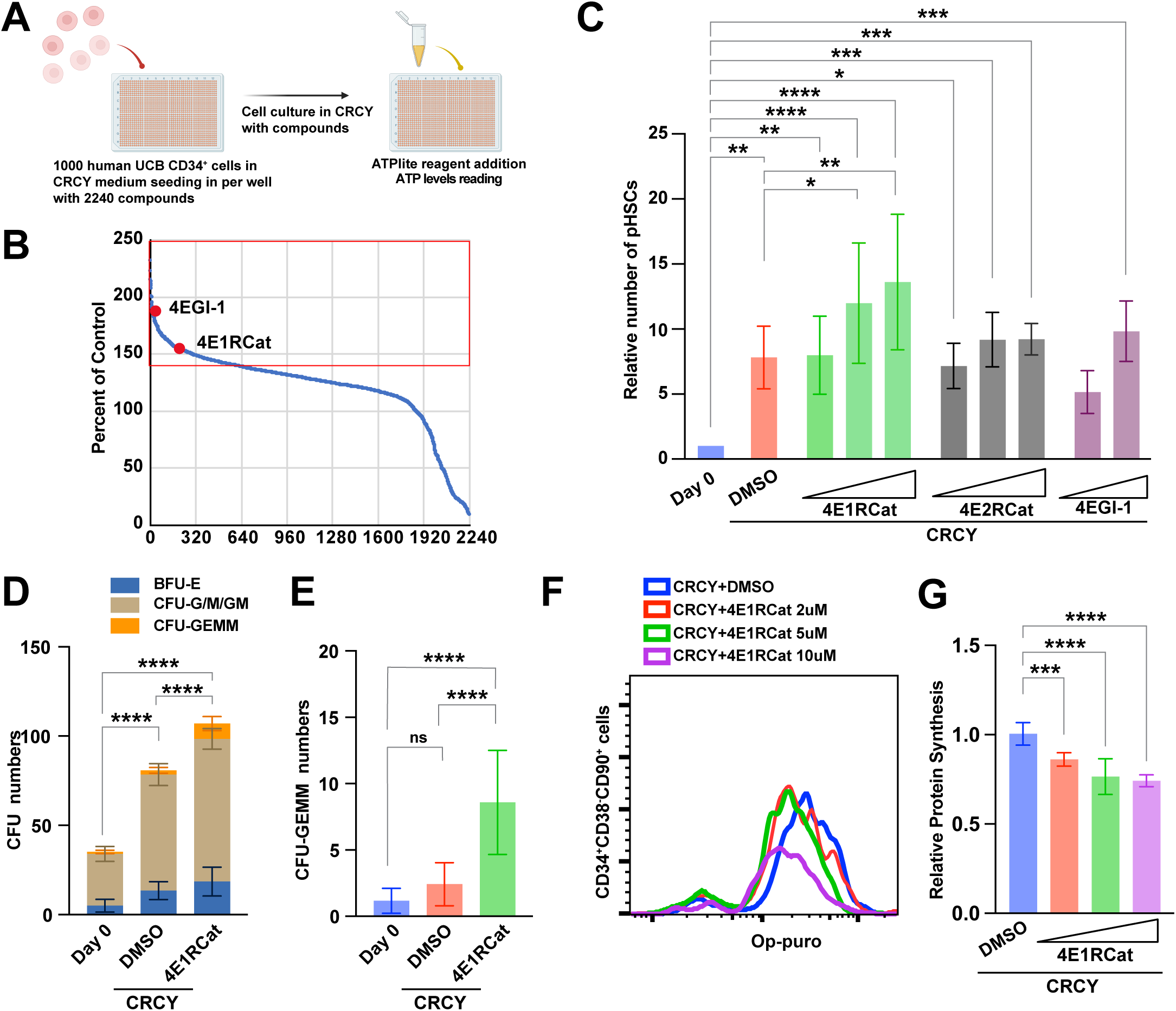
Small molecule inhibitors of translation initiation enhance ex vivo expansion of human pHSCs. **A.** HTS workflow: Human UCB CD34^+^ cells were added to 384 well dishes with vehicle (DMSO) or test compound from the Selleck bioactive compound library (>2240 compounds) in CRCY, cultured for 4 days, and cell number measured by ATP bioluminescence. **B.** Waterfall plot representing cell number as a percentage of control (DMSO) for each compound. Screen was performed twice at 1 µM and once at 0.1 µM and compounds identified as > 140% of control (Red box) in all 3 screens were selected for further study. 4E1RCat and 4EGI-1 are highlighted by red dots. **C.** CD34^+^ cells from UCB were cultured for 7 days in CRCY with vehicle (DMSO) or with increasing concentrations of 4E1RCat (500 nM, 2 µM, 10 µM), 4E2RCat (100 nM, 1 µM, 10 µM), or 4EGI-1 (1 µM, 10 µM) and then CD34^+^CD38^-^CD45RA^-^CD90^+^CD49f^+^ (pHSCs) cells were detected by flow cytometry. The number of pHSCs after 7 days of culture is shown relative to the number at Day 0 (freshly isolated cells). Data for Day 0, DMSO, and 4E1RCat represent the mean values from 6 biological replicates (6 samples of mixed donors). Data for 4E2RCat and 4EGI-1 show mean of 3 replicates (3 distinct mixed donors). The total CD34+ cell counts are shown in Supplemental Figure 13A. **D.** Colony forming units (CFU) were measured in uncultured (Day 0) CD34^+^ cells and cells cultured for 7 days in CRCY with vehicle (DMSO) or 4E1RCat (2 µM). Total number of mixed cell lineage CFUs or CFU-GEMM (granulocyte, erythrocyte, monocyte, megakaryocyte), CFU-G/M/GM (granulocyte, macrophage, granulocyte/macrophage) and BFU-E (burst forming units-erythroid) are shown in the left panel. **E.** Data from panel D showing multipotent progenitors as CFU-GEMMs with an expanded y-axis. Data show results from 3 distinct mixed-donor UCB samples. **F.** Inhibition of translation by 4E1RCat: CD34^+^CD38^-^CD90^+^ cells were cultured in CRCY and increasing concentrations of 4E1RCat (0 µM, 2 µM, 5 µM, 10 µM) and translation was measured by OP-Puro incorporation and flow cytometry. **G.** OP-Puro fluorescence is shown as relative median fluorescence intensity. For all panels, * indicates *p* < 0.05, ** indicates *p* < 0.01, *** indicates *p* < 0.001, **** indicates *p* < 0.0001, and NS indicates not significant. Statistical significance was calculated by one-way ANOVA for all panels.

**Figure 3.**
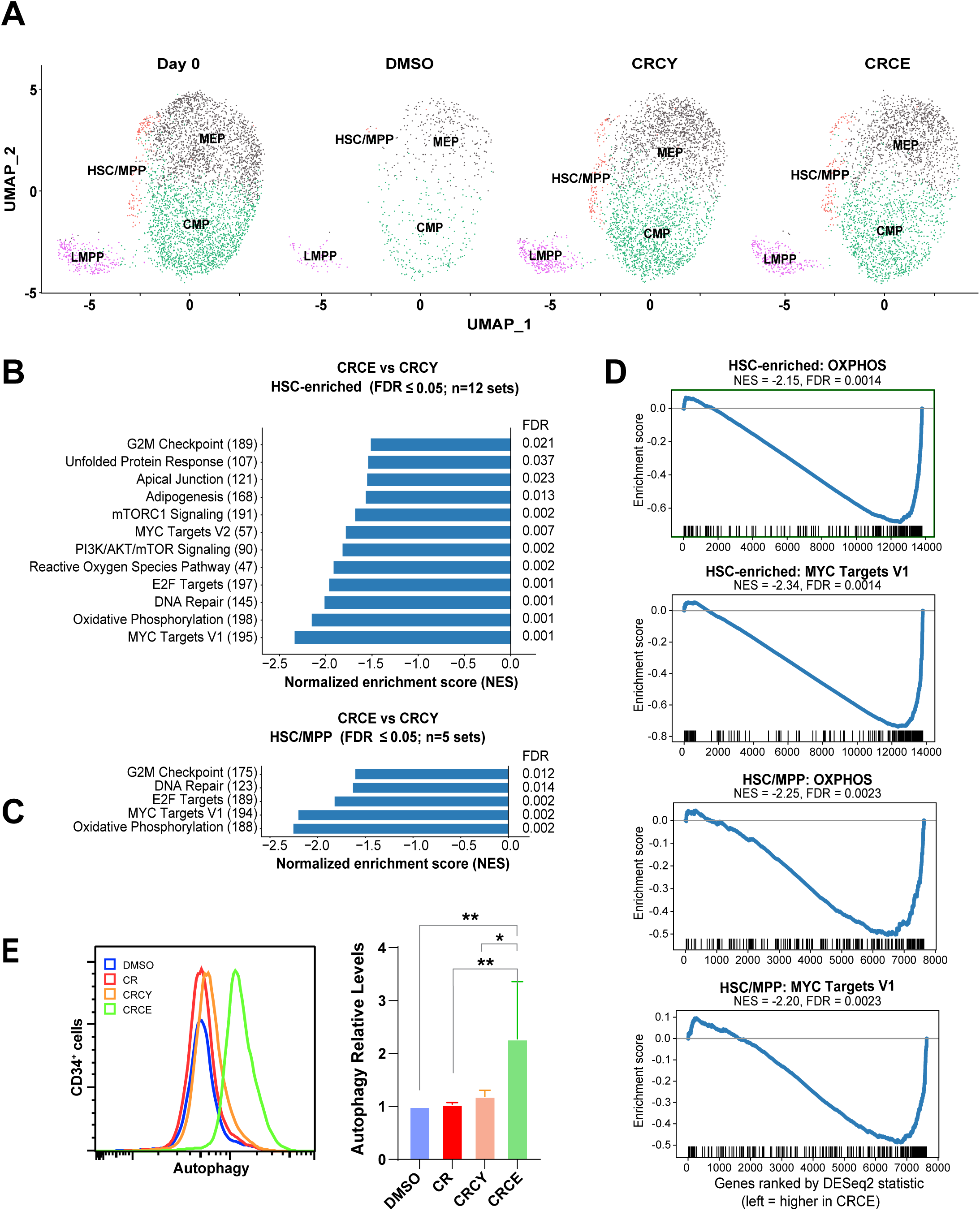
CRCE maintains an HSC/MPP signature and suppresses OXPHOS, MYC, and cell cycle gene expression programs. HSC-enriched CD34^+^CD38^-^CD45RA^-^CD90^+^ cells purified from UCB CD34^+^ cells were cultured in control medium+vehicle (DMSO), CRCY, or CRCE for 2 days and subjected to single cell transcriptomic analysis along with Day 0 (uncultured) cells. **A.** UMAP: HSC/MPP, CMP, MEP, LMPP and early erythroid commitment clusters were defined as in Figure 1. **B-D**. Gene set enrichment analysis (GSEA) of CRCE vs CRCY. Pre-ranked GSEA of the MSigDB Hallmark collection comparing CRCE with CRCY in CD34+CD38−CD45RA−CD90+ cells (“all cells”; panel B) or in the HSC/MPP cluster (C). Genes were ranked by the DESeq2 Wald statistic for CRCE vs CRCY; negative normalized enrichment scores (NES) indicate gene sets that are lower in CRCE. Bars show NES for Hallmark gene sets with FDR ≤ 0.05 (12 in “all cells”, 5 in the HSC/MPP cluster; Benjamini–Hochberg method across all Hallmark gene sets tested). The number of genes from each set found in the data is in parentheses, and FDR values are shown at right. (D) Enrichment plots for MYC targets V1 and oxidative phosphorylation in all cells (top) and HSC/MPP (bottom). E. Flow cytometric detection of autophagic vacuoles: CD34+ cells from UCB were cultured in control medium (DMSO), CR, CRCY, or CRCE for 24 hours and then analyzed by flow cytometry with a commercial autophagy detection kit to label autophagic vacuoles. Left: fluorescence for cells in control medium (DMSO), CR, CRCY, and CRCE. Right: Median fluorescence intensity (MFI) normalized to DMSO control. CD34+ cells were also stimulated with thapsigargin as a positive control for ER stress (Supplemental Figure 3B).* indicates p < 0.05; ** indicates p < 0.01 (one way ANOVA, N = 3 independent donors). GEO SubSeries Accession number GSE353932.

As the CD34^+^ population (HSPCs) contains HSCs and hematopoietic progenitor cells, we performed colony formation assays to assess the capacity of expanded HSPCs to differentiate into multilineage hematopoietic cell types. The number of colony-forming units (CFUs) generated from CD34^+^ cells cultured in CRCY or CRCE increased 2-3 fold compared to uncultured cells (Figure 2D). The number of multipotent progenitors that generate granulocyte, erythroid, macrophage, and megakaryocyte lineages (CFU–GEMM) was significantly increased in the CRCE group (Figure 2E). Thus, CRCE increases pHSCs and progenitor cell populations ex vivo.

We observe pHSC expansion with 4E1RCat at 2 µM, which is below the reported IC_50_ for inhibition of translation^69^. To confirm that 4E1RCat inhibits global translation in CD34^+^ cells under these conditions, we measured protein synthesis by incorporation of the fluorescent puromycin analog O-propargyl-puromycin (OP-Puro) ^2,71^. 4E1RCat decreased global translation in CD34^+^CD38^-^CD90^+^ populations in a dose-dependent manner, with maximal inhibition between 2 µM and 10 µM (Figure 2F and 2G).

To assess transcriptomic changes associated with ex vivo HSC expansion, we repeated scRNA-seq on CD34^+^CD38^-^CD45RA^-^CD90^+^ cells from UCB comparing cells at day 0 to cells cultured for 2 days in CRCE, CRCY, or vehicle control (DMSO). UMAP visualization showed that culture in CRCE maintains the HSC/MPP population at a proportion similar to Day 0 cells and cells cultured in CRCY (Figure 3) whereas the HSC/MPP population is reduced in control medium (DMSO, Figure 3A, see also Figure 1A). Culture in CRCE was associated with a reduction in UPR/stress response genes similar to CRCY when compared to Day 0 cells or cells cultured in medium alone (DMSO; Supplemental Figure 3A) and this was confirmed by quantitative reverse transcription PCR (RT-qPCR) for selected UPR and IEG genes in CD34^+^ cells (Supplemental Figure 3B,C).

We then asked whether 4E1RCat altered coordinated transcriptional programs by performing gene set enrichment analysis (GSEA) of HSC-enriched cells, comparing cells cultured in CRCE to culture in CRCY, as these conditions differ only by the inclusion of 4E1RCat. GSEA revealed a significant reduction in expression of gene sets associated with oxidative phosphorylation (OXPHOS), c-MYC targets, and cell cycle regulators (Figure 3B, Supplemental Table 4) for cells cultured in CRCE. Most genes in the MYC (V1) and OXPHOS programs were significantly reduced (Supplemental Figure 4). The HSC/MPP cluster also showed reduced expression of OXPHOS and MYC target genes despite the lower number of cells for analysis (Figure 3C. As translation initiation inhibitors are known to activate autophagy^72–74^, we measured autophagy by flow cytometry in CD34^+^ cells cultured in CRCE or CRCY and observed a dose-dependent increase in autophagic vacuole content in cells receiving 4E1RCat (Figure 3E, Supplemental Figure 5).

### Expansion of functional HSCs

As a rigorous test of LT-HSC expansion by culture in CRCE, we performed limiting dilution repopulation assays (LDA) in immunocompromised mice. CD34^+^ cells at Day 0 or after 7 days of culture in CRCY or CRCE were injected at varying cell doses into busulfan-conditioned non-obese diabetic severe combined immunodeficient IL-2Rγ^null^ (NSG) mice ^51,75^. Bone marrow aspirates were collected at 20 weeks after transplantation and human cell chimerism was assessed by flow cytometry. The number of positive chimeric mice, defined as > 0.1% donor-derived human cells in bone marrow, was significantly higher in the group receiving CD34^+^ cells cultured in CRCE compared to the uncultured group or the group cultured in CRCY (Figure 4A; Supplemental Figure 6). The number of severe combined immunodeficiency (SCID)-repopulating cells (SRCs), a measure of the fraction of functionally engrafting LT-HSCs in cells cultured in CRCE (1 in 187) was 5.7-fold higher than cells cultured in CRCY (1 in 1074; *p* = 0.0005) and 4.3-fold higher than uncultured CD34^+^ cells (1 in 794; *p* = 0.0001) (Figure 4B). The absolute number of SRCs (and the number of SRCs per day 0 equivalent) also increased in CRCE (Supplemental Figure 7). No significant difference in LT-HSC number was observed with CRCY compared to the uncultured group (Figure 4A, B), consistent with HSC maintenance we reported previously for CR medium^51^. Multilineage reconstitution was detected at 20 weeks in the CRCE group, as a substantial percentage of bone marrow cells were donor-derived, including T cells (CD3^+^), B cells (CD19^+^), myeloid cells (CD33^+^), and hematopoietic progenitor cells (CD34^+^CD38^-^) (Figure 4C-F), indicating that CRCE confers multilineage reconstitution compared to the uncultured group. Multilineage reconstitution was also observed at 29 weeks (Supplemental Figure 8).

**Figure 4.**
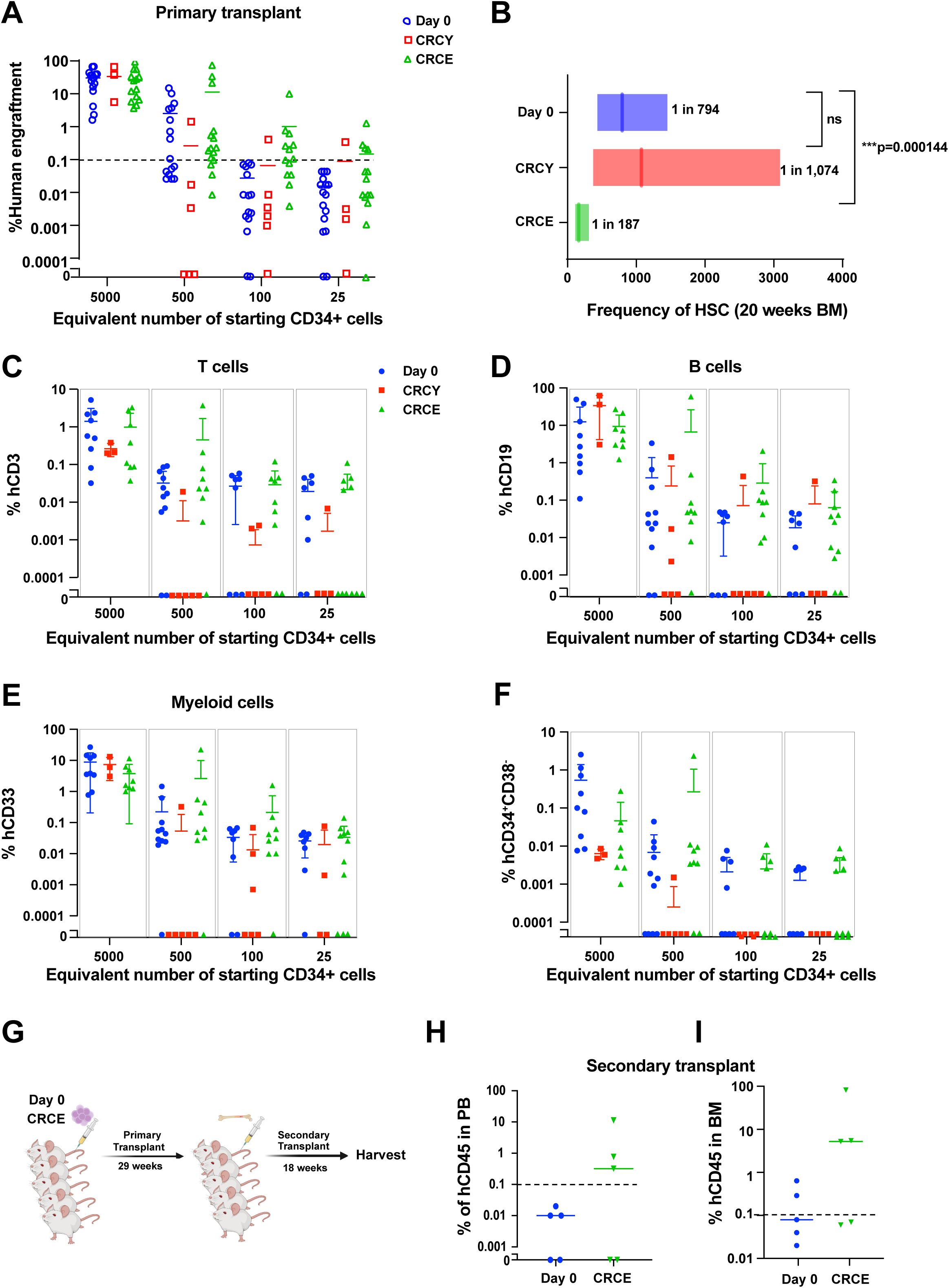
CRCE promotes HSC expansion. **A.** Limiting dilution analysis (LDA) of human UCB CD34^+^ cells at Day 0 or after 7 days of culture in CRCY+DMSO or CRCE. Dose of cells injected is based on the number of CD34^+^ cells on day 0 (day 0 equivalent). Chimerism was measured as human CD45^+^ cells at 20 weeks in bone marrow with engraftment defined as ≥ 0.1% hCD45^+^. Data are from 3 independent experiments using pooled UCB donors. **B.** HSC frequency with 95% CI calculated with ELDA software. *p* = 0.0001 for CRCE compared to fresh cells. No significant difference for CRCY compared to Day 0 cells. Total cell numbers and number of LT-HSCs (SRCs) are shown in Supplemental Figure 7. **C-F.** Lineage analysis (from 2 independent transplants using pooled UCB donors): The percentage of human T cells (CD3^+^ cells), B cells (CD19^+^ cells), myeloid cells (CD33^+^ cells), and lineage negative (CD34^+^CD38^-^) cells is shown for recipients injected with cells from ex vivo cultures corresponding to 5000, 500, 100, 25 CD34^+^ cells on Day 0. **G.** Scheme for secondary transplantation from primary recipients that had received Day 0 cells or cells cultured in CRCE for 7 days. **H.** The percentage engraftment of human CD45^+^ cells in peripheral blood (16 weeks) in secondary recipients. **I.** The percentage engraftment of human CD45^+^ cells in the bone marrow (18 weeks) in secondary recipients. Lineage analysis in Supplemental Figure 9.

To further demonstrate the capacity for long-term self-renewal after ex vivo expansion in CRCE, we performed serial transplantation into secondary recipients using bone marrow cells harvested from five donors per condition at 29 weeks post-transplant (Figure 4G-I, Supplemental Figure 9). Human chimerism was detected in the peripheral blood (PB) of secondary recipients in the 4E1RCat-treated group but not the uncultured group at 16 weeks post-transplant (Figure 4H). Bone marrow from secondary recipients at 18 weeks achieved a higher percentage of human chimerism (hCD45^+^), hematopoietic progenitor cells (CD34^+^CD38^-^), B cells (CD19^+^), myeloid cells (CD33^+^), megakaryocytic cells (CD41^+^), and erythroid cells (CD235a^+^) compared to Day 0 (uncultured) donor cells in secondary recipients (Figure 4I, Supplemental Figure 9). Thus, culture of UCB CD34^+^ cells in CRCE promotes the expansion of HSCs capable of long-term regeneration.

### Maintenance and expansion of CRISPR modified adult CD34^+^ cells

A major obstacle to therapeutic gene editing for inherited hematopoietic disorders such as sickle cell disease (SCD) is the loss of functional HSCs after culture in cytokine-rich media. Because durable clinical benefit requires long-term engraftment of edited LT-HSCs, any loss of these cells during culture limits therapeutic efficacy. Our approach could overcome this obstacle by maintaining LT-HSCs during ex vivo manipulation. Human fetal red blood cells express primarily γ-globin chains that are paired with two α-globin chains to form fetal hemoglobin (α_2_γ_2_; HbF). After birth the γ-globin (*HBG1/2*) genes are transcriptionally silenced by repressors such as *BCL11A* and the β-globin gene (*HBB*) is activated to produce adult hemoglobin (α_2_β_2_; HbA)^76–78^. Elevated HbF levels due to genetic variation or through therapeutic HbF inducers attenuate the severity of SCD. Thus, induction of HbF by targeting the *BCL11A* repressor has been a long-standing goal in the field. Although a recent report indicated that erythroid specific loss of *BCL11A* may impair erythroid precursor expansion ex vivo and repopulation of erythroid cells in human/mouse xenografts^79^, it has nevertheless shown promising outcomes in the treatment of SCD patients^80,81^. However, the limited number of viable CD34^+^ cells that can be mobilized from patients presents a major obstacle to this approach, and this is exacerbated by the loss of functional HSCs after ex vivo manipulations that typically involve exposure to high levels of multiple cytokines. Our low cytokine conditions (with GSK-3 and mTORC1 inhibitors) could address these limitations by improving the ex vivo maintenance of HSCs and potentially increasing their numbers prior to autologous transplantation.

To test whether CRCE can maintain or expand gene-edited HSCs, we targeted the *BCL11A*+58 erythroid-specific enhancer, using CRISPR/Cas9-based RNP editing in mobilized human CD34^+^ cells from adult donors and then cultured in CRCY or CRCE for 7 days (Figure 5A). The absolute number of pHSCs increased 10-to 15-fold in non-edited and *BCL11A*+58-edited pHSCs compared to uncultured cells (Figure 5B), demonstrating ex vivo expansion of edited adult pHSCs. Colony formation assays demonstrated that the capacity for hematopoietic differentiation also increased 3-to 5-fold in edited CD34^+^ cells cultured in CRCY or CRCE; CRCE significantly increased the number of burst-forming unit-erythroid (BFU-E), granulocyte and macrophage CFUs (CFU-G/M/GM), and multipotent progenitors (CFU–GEMM) compared to CRCY or uncultured cells (Figure 5C,D). RT-qPCR confirmed that *BCL11A* expression in edited cells cultured for 7 days was reduced to ∼30% of expression in unedited cells at Day 0 (*p* < 0.01; Figure 5E), although exposure to 4E1RCat also reduced *BCL11A* expression independently in non-edited cells. Nevertheless, the reduction in *BCL11A* expression in edited cells cultured in CRCE was significantly lower than in non-edited cells (p = 0.027; Supplemental Figure 10). Importantly, *BCL11A*+58-edited cells cultured in CRCE maintained the increase in the percentage of HbF+ cells and the increase in *HBG* mRNA expression observed in edited cells not subjected to culture (Figure 5F, G). Similarly, culture in CRCE (or CRCY) did not disrupt erythroid maturation (Figure 5H). These observations demonstrate that culture in CRCE does not interfere with *BCL11A*+58 editing, HbF induction, or erythroid differentiation ex vivo.

**Figure 5.**
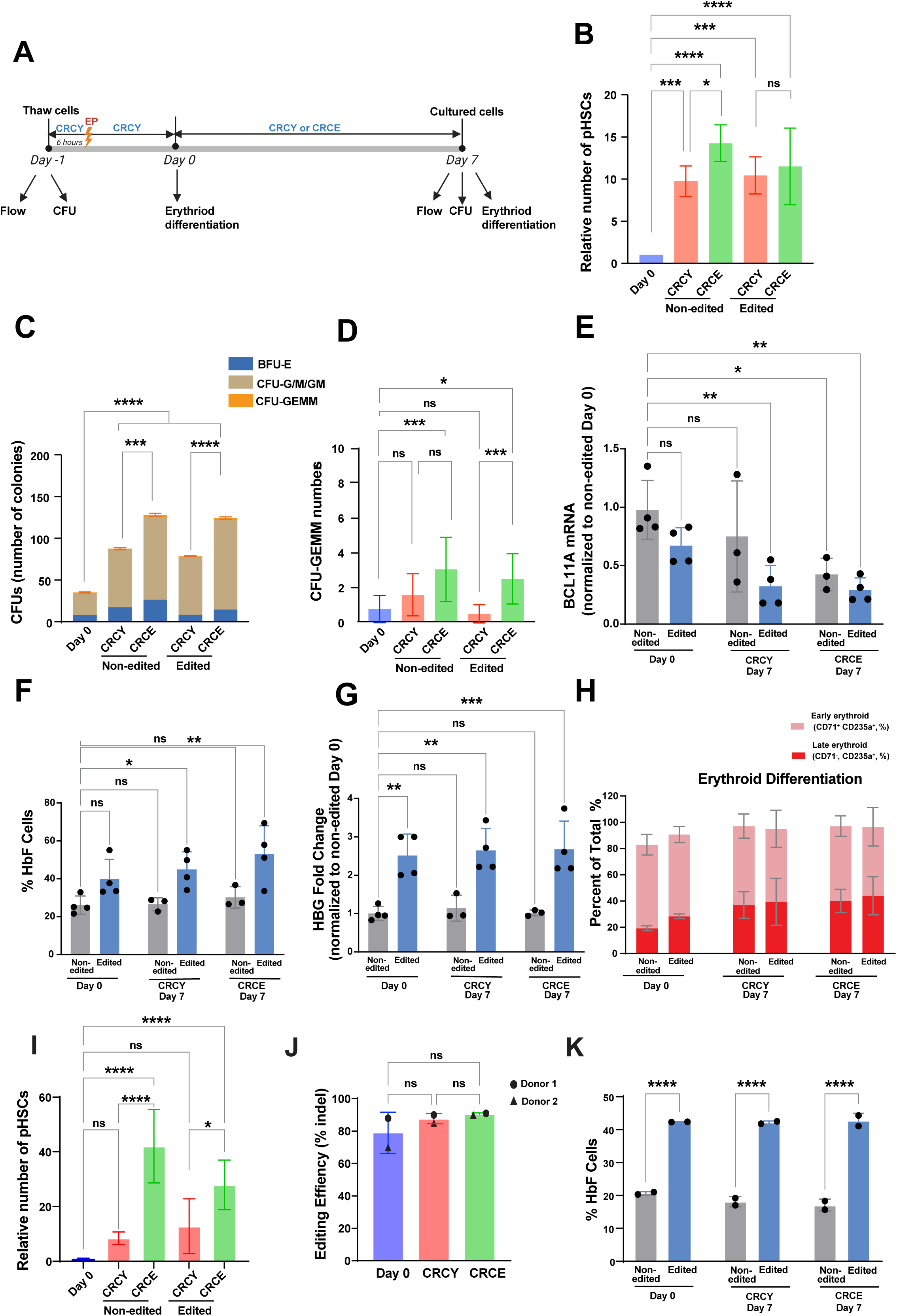
Ex vivo expansion of CRISPR modified CD34^+^ cells. **A**. Experimental plan and timeline. Mobilized adult CD34+ cells were thawed on Day-1 and subjected to flow cytometry (flow) and colony formation (CFU). Cells were placed in CRCY and electroporated (EP) at 6 hours, then returned to CRCY. An aliquot of cells was removed at Day 0 for erythroid differentiation assay and then culture in CRCY or CRCE (note [4E1RCat] = 100 nM for these experiments) was initiated. On Day 7, cells were harvested for flow cytometry, colony formation, and erythroid differentiation. **B.** Mobilized adult CD34^+^ cells were edited by CRISPR/Cas9 targeting of the *BCL11A*+58 enhancer and then edited cells and non-edited control cells were cultured in CRCY or CRCE. (Absolute number of CD34^+^ cells is shown in Supplemental Figure 13B). The number of pHSCs (CD34^+^CD38^-^CD45RA^-^CD90^+^CD49f^+^) at Day 7 in culture with CRCY or CRCE was normalized to the number at Day 0. (mean number pHSCs = 181±20 at Day 0). **C.** Colony formation was assessed as in Figure 2 for uncultured (Day-1) CD34^+^ cells and cells cultured for 7 days in CRCY or CRCE. The data represent replicates from 3 adult donors. **D.** Data from panel B with expanded y-axis to show the number of multipotent progenitors as CFU-GEMMs. **E-H.** After electroporation, CD34^+^ cells from Day 0 (±EP) and Day 7 (±EP) of culture in CRCY or CRCE were differentiated in three-phase erythroid culture and collected at Day 13 for detection of **(E)** *BCL11A* mRNA by RT-qPCR; (**F)** %HbF+ cells (F-cells) by flow cytometry; (**G**) *HBG*% mRNA expression by RT-qPCR; and (**H**) erythroid differentiation by flow cytometry. Data represent the mean of replicates from 3-4 adult healthy donors. **I.** The *BCL11A*+58 erythroid enhancer was edited in SCD patient-derived CD34^+^ cells. Edited and non-edited SCD CD34^+^ cells were cultured for 7 days in CRCY or CRCE. (The absolute number of pHSCs at day 0 was 115 for donor 1 and 153 for donor 2) **J, K.** Edited and non-edited cells were subjected to erythroid differentiation and then assessed for BCL11A+58 editing efficiency (J) and %HbF+ cells (F-cells) by flow cytometry (K). The data represent the mean of replicates of patient-derived CD34+ cells from 2 SCD donors. For all panels, * indicates *p* < 0.05, ** indicates *p* < 0.01, *** indicates *p* < 0.001, **** indicates *p* < 0.0001. NS, not significant (one-way ANOVA).

To assess whether CRCE maintains or expands edited primary cells from patients with SCD, we isolated CD34^+^ cells from deidentified SCD patients undergoing therapeutic red cell exchange, performed CRISPR-Cas9-RNP editing of the *BCL11A*+58 erythroid enhancer, and cultured both edited and non-edited cells with CRCY and CRCE. The absolute number of pHSCs increased 13-fold in CRCY and nearly 28-fold in CRCE (Figure 5I, Supplemental Figure 13). Editing efficiency was ∼80-90% and was not significantly affected by CRCE in two independent donors (Figure 5J). In addition, high levels of HbF induction were maintained following CRCE expansion of edited cells (Figure 5K). These data indicate that HbF induction after editing of *BCL11A*+58 is equivalent, and potentially more robust, in SCD patient-derived CD34^+^ cells cultured in CRCE (Figure 5K) compared to healthy donors (Figure 5F).

To test whether our culture conditions maintain long-term HSCs after CRISPR editing, we cultured edited CD34^+^ cells in CRCY or CRCE for 7 days and transplanted them into busulfan-conditioned NBSGW mice^82^ (Figure 6A). Uncultured CD34^+^ cells from the same donor were transplanted as well. The number of day 7 cells injected was based on day 0 equivalent cells so that an equal proportion of the culture for either day 0 or day 7 was transplanted. The NBSGW strain was selected as it provides improved erythroid engraftment compared to NSG mice^82–84^. Bone marrow was harvested at 20 weeks and human cell engraftment was assessed. Human cell chimerism for CRISPR edited cells cultured in CRCY or CRCE was equivalent to engraftment of freshly isolated (uncultured and unedited) cells (Figure 6B), indicating maintenance of HSCs after CRISPR editing. We also observed multilineage reconstitution, with robust engraftment of primitive hematopoietic stem and progenitor cells (CD34^+^CD38^-^), lymphocyte, myeloid, megakaryocytic, and erythroid lineages that was similar for unedited and edited/cultured cells (Figure 6C-H), although engraftment of CD3^+^ T cells was low for all conditions (Figure 6F). A recent report has similarly shown that UM171-based culture enhances fitness and engraftment of sickle cell disease patient HSCs transduced with lentivirus expressing short hairpin RNA to suppress *BCL11A*^85^. We then measured HbF (by flow cytometry) and *HBG* expression (by RT-qPCR) in erythroid cells harvested from the bone marrow of transplanted mice at 20 weeks. CRISPR editing of the *BCL11A*+58 erythroid enhancer increased the mean percentage of HbF^+^ erythroid cells by ∼3-fold (*p* < 0.05) and increased *HBG* mRNA expression by nearly 5-fold compared to unedited cells (*p* < 0.05) when cells were cultured in CRCE (Figure 6I, J). These data demonstrate maintenance of genome edited HSCs from adult donors under low cytokine conditions with CRCY and CRCE medium and provide a potential therapeutic method to ensure robust transplantation of CD34^+^ products in SCD patients undergoing gene therapy.

**Figure 6.**
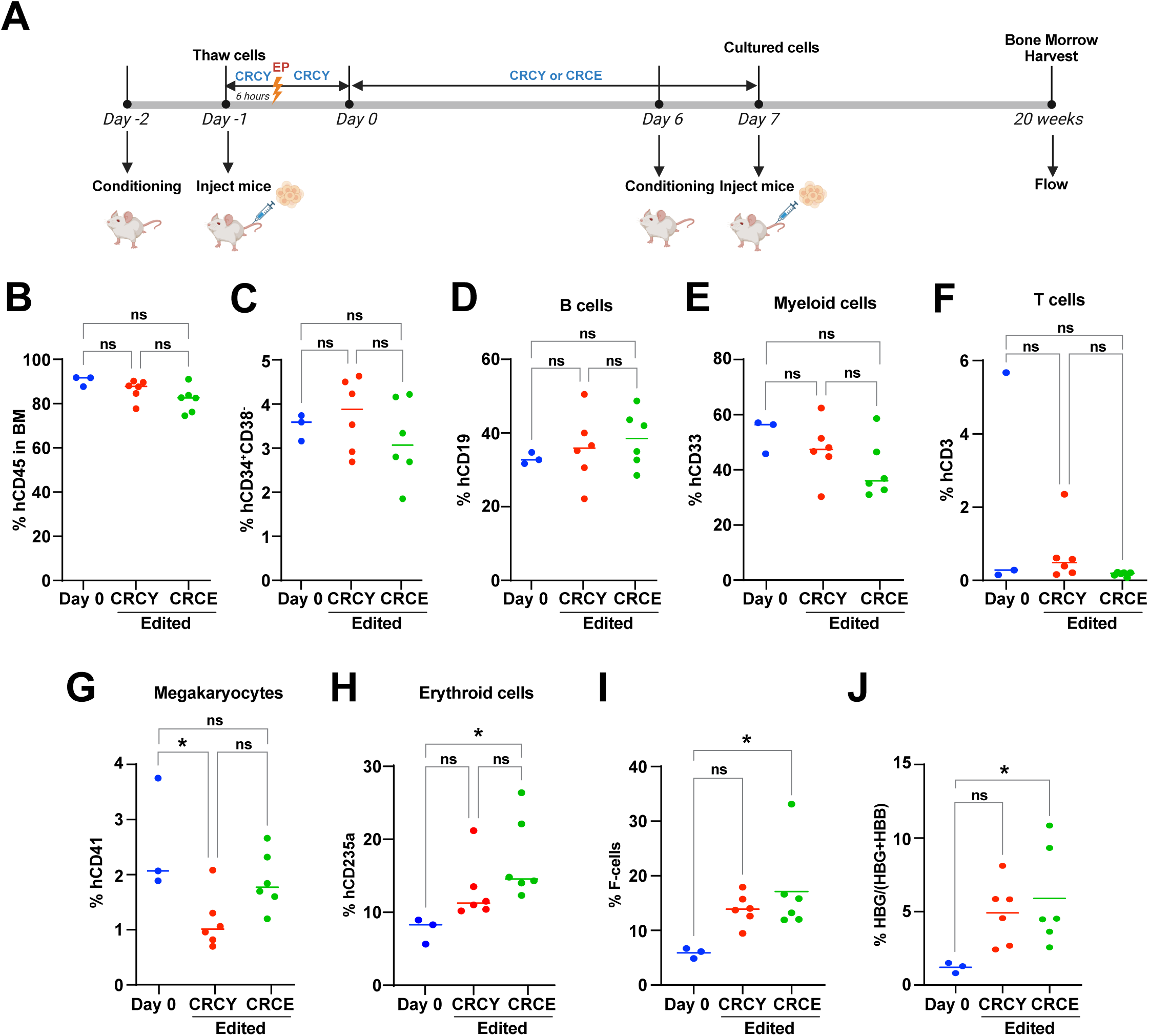
Engraftment in NBSGW mice of edited adult HSCs after culture in CRCY with or without 4E1RCat. **A.** Experimental plan and timeline for transplant of *BCL11A*+58 edited adult CD34+ cells in NBSGW mice. Uncultured CD34+ cells and cells that were electroporated with *BCL11A* sgRNA and then cultured in CRCY or CRCE for 7 days were transplanted into busulfan conditioned NBSGW mice and bone marrow was harvested after 20 weeks. **B.** Total human cell engraftment as % human CD45^+^ cells. (The absolute number of engrafted human cells/mouse is shown in Supplemental Figure 13D). **C.** hCD34^+^CD38^-^ (lineage negative) hematopoietic progenitor cells, **D.** hCD19^+^ B cells, **E.** hCD33^+^ myeloid cells, **F.** hCD3^+^ T cells, **G.** hCD41^+^ megakaryocytic cells, **H**. hCD235a^+^ erythroid lineage cells. **I.** HbF+ cells (F-cells) measured by flow cytometry and represented as percent of stage-matched hCD71^+^ hCD235a^+^ cells harvested from bone marrow at 20 weeks post transplant (mice do not express HbF). **J.** Human %*HBG* mRNA expression (RT-qPCR with human-specific primers, expressed as *HBG*/(*HBG*+*HBB*) x100%) in total marrow cells harvested at 20 weeks from NBSGW mice transplanted with uncultured control cells (n=3 mice), cells cultured in CRCY (n=6 mice), or CRCE (n=6 mice). * indicates *p* < 0.05 (one-way ANOVA).

To test whether CRCE can maintain or expand CRISPR/Cas9-edited HSCs in other clinically relevant editing modalities, we targeted the *BCL11A*+58 erythroid-specific enhancer using clinically scalable MaxCyte electroporation (EP) with a TadA-based cytosine base editor (CBE-EP) or lipid nanoparticle (LNP) transduction of conventional CRISPR/Cas9. These were compared directly to EP with standard CRISPR/Cas9-RNP-based approaches (Lonza 4D nucleofection, as in Figure 5). All groups were cultured in CRCY or CRCE for 7 days. Compared to uncultured cells, the absolute number of pHSCs increased 10-to 15-fold with conventional CRISPR/Cas9 editing delivered by EP or by LNPs and ∼40-fold with CBE-EP (Supplemental Figure 11A). Day 0 and Day 7 cells were subjected to 3-phase erythroid differentiation^62,63^ and assessed for expression of *BCL11A* and *HBG* mRNAs and percentage of HbF^+^ cells. Compared to Day 0 unedited cells, *BCL11A*+58-edited cells from all 3 groups cultured for one week in CRCY or CRCE demonstrated reduced *BCL11A* (Supplemental Figure 11B) and increased *HBG* mRNA expression (Supplemental Figure 11D). Importantly, elevated *HBG* expression and %HbF+ cells were maintained in edited cells cultured for 7 days compared to Day 0 edited cells (Supplemental Figure 11C,D). Erythroid maturation was also unaffected after culture in CRCY or CRCE (Supplemental Figure 11E). These data indicate that our HSC culture system maintains adult HSCs independently of the delivery technology and may be broadly applicable for therapeutic gene editing.

## Discussion

Ex vivo expansion of human HSCs from UCB has tremendous therapeutic potential for hematopoietic malignancies and bone marrow failure, but a major challenge has been the loss of self-renewing HSCs when cells are stimulated with hematopoietic cytokines. Similarly, gene therapy for inherited blood disorders such as sickle cell disease and β-thalassemia has been limited by the low recovery of functional HSCs, which may also be due, in part, to high cytokine exposure during ex vivo manipulations. Culture in cytokine-rich medium suppresses autophagy, increases oxidative phosphoryilation, and increases protein synthesis, which must be restricted to maintain the capacity for self-renewal in HSCs^54,55^. These changes are detrimental to the expansion of functional, long-term HSCs^1–5,10,11,54–59,86,87^. Our approach limits cytokine exposure (both concentration and duration) and reduces translation initiation, achieving > 4-fold expansion of long-term HSCs based on limiting dilution assays in immunocompromised mice. This scale of HSC expansion is more than sufficient to allow the use of the many stored UCB units that are just under the threshold of CD34^+^ cell counts for clinical use. Indeed, a two-fold expansion should, in principle, be equivalent to the clinically used double cord approach, and would avoid the problem of immunological extinction of cells from one of the two donors. These conditions also maintain adult human HSCs after CRISPR/Cas9-mediated gene editing of the *BCL11A*+58 enhancer, with sustained *HBG* and HbF induction in edited human erythroid cells at 20 weeks after transplantation, potentially overcoming a major obstacle to gene therapy for sickle cell disease, β-thalassemia, and other inherited blood disorders.

Other promising approaches for ex vivo HSC expansion are at various stages of development^26–40,47–50,85^. Our method is distinct from those approaches because we have reduced cytokine exposure substantially. However, our data reveal interesting parallels with two of the more advanced expansion protocols, UM171 and Omidubicel. UM171, a “molecular glue degrader” that induces proteosomal degradation of the LSD1–CoREST corepressor complex^45,88^, also reduces global translation, suppresses MYC target gene expression, increases expression of a lysosomal/autophagy gene expression program referred to as the CLEAR (Coordinated Lysosomal Expression And Regulation) network^89^; this in turn limits HSC activation and enhances self-renewal^90,91^. Furthermore, nicotinamide, a key ingredient in the Omidubicel process, also stimulates autophagy ^92–94^. Mitofusin agonists also enhance expansion of HSCs from UCB by suppressing translation, inhibiting mTORC1, and increasing autophagy^47^. Here we show that 4E1RCat similarly reduces translation, suppresses OXPHOS, MYC, and cell cycle gene expression programs, and increases autophagic vacuole content. Culture in CRCE also increases expression of multiple CLEAR network genes (Supplemental Figure 12). These parallels may suggest a common downstream mechanism for these distinct HSC expansion methods or at least reflect an LT-HSC state that is enhanced by each of these approaches.

Single-cell transcriptomic analysis of cord blood cells enriched for HSCs (CD34^+^CD38^-^CD45RA^-^ CD90^+^) reveals several interesting features. Importantly, the HSC/MPP population is maintained when cells are cultured with inhibitors of GSK-3 and mTORC1 (CR medium), but not with vehicle control medium, consistent with our prior functional studies showing that CR maintains LT-HSCs in cytokine-free culture^12,51^. HSC/MPPs were also maintained when low-dose cytokines were added (Figures 1A) and when 4E1RCat (CRCE) was included (Figures 3 and 4). Thus, an HSC/MPP transcriptomic signature is maintained in parallel with long-term self-renewal in both cytokine-free and low cytokine conditions that include CR (e.g. CR, CRCY, and CRCE). ScRNA-seq also showed that the unfolded protein response sensor *HSPA5*, which encodes the chaperone GRP78 (BiP)^65^, is enriched in the HSC/MPP population. Multiple IEGs, including several *JUN* family transcription factors, are also significantly enriched in HSC/MPPs compared to other cell types. These observations are consistent with prior work showing that UPR components, including *HSPA5*, are enriched in HSCs and essential for maintaining HSC self-renewal during hematopoietic stress^1,54,56,95,96^. However, sc-RNA-seq and RT-qPCR also showed that UPR markers and IEGs are high in day 0 (uncultured) CD34+ cells (HSPCs), CD34^+^CD38^-^CD45RA^-^CD90^+^ cells, and the HSC/MPP population relative to cells cultured in CR-containing media, consistent with the well-known stress of freeze-thawing and cell sorting^67,68^. These stress markers were also reduced in cells cultured in CR-containing media compared to cells cultured in control medium (DMSO), correlating with improved overall survival and maintenance of LT-HSCs in CR, CRCY, and CRCE.

Among its many functions, mTORC1 activates translation but the mRNA targets of mTORC1 regulation are surprisingly limited. mRNAs with 5’ polypyrimidine tracts are particularly sensitive to inhibition by rapamycin^97^. Similarly, concentrations of 4E1RCat that enhance HSC expansion only partially inhibit translation. This could indicate that a subset of mRNAs has higher sensitivity to 4E1RCat, but, alternatively, the partial reduction in translation with 4E1RCat may improve HSC maintenance by reducing overall proteostatic stress. Consistent with this interpretation, partial reduction in unfolded proteins in the endoplasmic reticulum improves HSC fitness and, conversely, increased misfolded protein impairs in vivo repopulation in human:mouse xenografts^54,56^. Similarly, small increases in the rate of protein synthesis impair HSC function^55^. Thus, the HSC expansion we observe with modest reduction in global protein synthesis by 4E1RCat may be due to a reduction in proteostatic stress. It is also possible that the combination of rapamycin, an mTORC1 inhibitor, and 4E1RCat, an inhibitor of eIF4F assembly, have additive effects to suppress translation. However, the rapamycin in CR (5nM) is well below typically used concentrations and by itself does not cause detectable inhibition of translation at this concentration; furthermore, other translation inhibitors cannot replace rapamycin to maintain HSCs in CR-containing media^12^, suggesting that mTORC1 inhibition contributes to HSC maintenance through a distinct mechanism. In contrast, 4E1RCat does modestly reduce overall translation and, as we show here, substantially increases autophagy, similar to the effects of other eIF4F inhibitors^72–74^. The increase in autophagy induced by 4E1RCat is consistent with prior work showing that increased autophagy plays an essential role in maintaining LT-HSCs ex vivo^5,87^.

FDA-approved gene therapy approaches for SCD and β-thalassemia include lentivirus-mediated expression of modified β-globin^98,99^ and CRISPR-editing of the erythroid-specific *BCL11A*+58 enhancer to enhance HbF expression^81^. However, ex vivo manipulation and exposure to hematopoietic cytokines are associated with proliferative stress that impairs HSC fitness and engraftment, leading to loss of functional HSCs after culture in cytokine-rich media. Alternative methods include in vivo editing using lipid nanoparticle delivery of mRNAs^100,101^ and approaches to reduce proliferative stress and increase HSC fitness during ex vivo culture, as shown here by reducing cytokine exposure and reducing translation ex vivo in CRISPR-edited HSCs, and also shown recently by culture of lentivirus-transduced HSCs in UM171-based medium^85^. Importantly, we also observe increased HbF expression in human erythroid cells derived from *BCL11A*-edited HSCs cultured in CRCE. Furthermore, the level of HbF induction we observe here (∼2-fold increase in HbF+ cells) is associated with at least 50% reduction in cellular sickling^63,102^, consistent with the well-established concept that small changes in F-cell number can have much broader phenotypic benefit. Furthermore, clinical studies and meta-analyses show that patients with a ∼1.5-2-fold increase in HbF% and F-cell% have a substantial reduction in vaso-occlusive crises^103^. Therefore, the >2-fold induction of HbF that we achieve in healthy and SCD donors is well within the range for clinically meaningful effects.

In summary, we have identified a protein synthesis inhibitor that, in combination with low cytokine exposure, confers >four-fold expansion of human HSCs from umbilical cord blood. This degree of expansion is modest by intention, and is more than sufficient to make a large number of stored UCB units available for HSCT. This degree of expansion would also surpass the quantity of CD34^+^ cells used clinically in “double cord” HSCT. This approach may benefit patients from genetic and ethnic backgrounds that are not well represented in bone marrow registries and those who are not suitable for haploidentical transplants. Furthermore, the approach may overcome a major obstacle to gene therapy for inherited blood disorders by maintaining mobilized adult HSCs after gene editing.

## Acknowledgements

We thank the UPenn High-throughput screening core, RRID: SCD_022379, for their assistance with the high-throughput screen. We thank Dr. Ana Domingo Muelas for help with graphics, Dr. Ivan Maillard for comments on the manuscript, Drs Martin Carroll, Nicolas Skuli, and Martin Keough, Anthony Secreto, and the staff of the Stem Cell and Xenograft Core at the University of Pennsylvania for guidance on xenograft experimental design.

## Conflict of Interest

SAP is a consultant for Genetix Biotherapeutics, Agios Pharmaceuticals, and Fulcrum Therapeutis. SAP had a recent sponsored research agreement with Blueprint Medicines and PSK had an independent sponsored research agreement with Blueprint Medicines that is no longer active and is unrelated to the research reported here.

Data Sharing: Single cell RNA-seq data were deposited in the NCBI Gene Expression Omnibus and are accessible through GEO SuperSeries accession number GSE353936. All other data are contained within the manuscript and supplemental data.

## Author contributions

CL designed experiments, performed research, analyzed data, wrote the paper, and edited the paper.

HSh designed experiments, performed research, analyzed data, and edited the paper.

DB designed experiments, performed research, and analyzed data.

ML analyzed data.

MIB performed research and analyzed data. XY analyzed data.

DCS provided guidance on HTS, performed experiments, and analyzed data.

SC provided guidance on HTS and analyzed data.

XL performed research and analyzed data.

FH assisted with data analysis and provided vital resources.

HSa assisted with experimental design and performed experiments.

LB assisted with experimental design and data analysis, performed experiments

M-GA assisted with experimental design and provided vital resources.

LB assisted with experimental design and data analysis, performed experiments, and provided vital resources.

NK performed research and analyzed data.

YK performed research and analyzed data.

NS performed research and analyzed data.

SR assisted with experimental design and data analysis, and provided viral resources.

JRA-D assisted with data analysis and provided vital resources.

GAB provided vital reagents and resources.

BDG assisted with data analysis and provided vital resources.

SAP assisted with experimental design, performed research, analyzed data, provided vital resources, and edited the paper.

JH assisted with experimental design and data analysis and provided vital resources.

PSK designed experiments, analyzed data, wrote and edited the paper, supervised the overall project, and provided vital resources.

## Supplemental Data Files

1. Supplemental Figures 1-14 with legends.

2. Supplemental Methods

3. SupplementalTable 1: Differential gene expression in specific populations

4. SupplementalTable 2: GO metascape

5. SupplementalTable 3: HTS data summary

6. Supplemental Table 4: GSEA

**Supplemental Figure 1.**
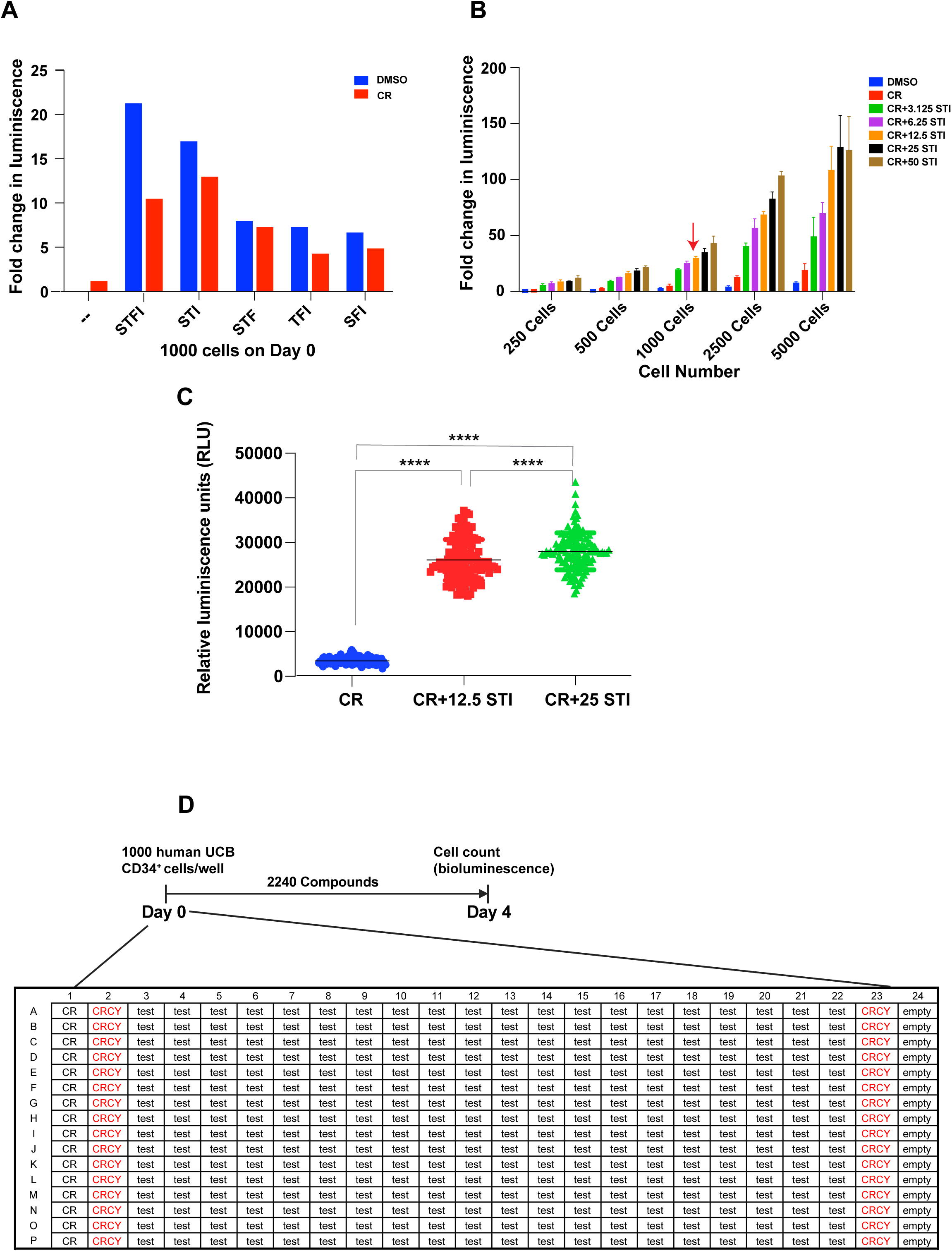
Optimization of screening conditions. **A.** Human UCB CD34^+^ cells were added to 384 well dishes in StemSpan SFEM with (red) or without (blue) CR and varying combinations of cytokines as indicated. S: SCF (100 ng/ml), T; Thrombopoietin (100 ng/ml), F: Flt3 ligand (100 ng/ml), I: IL3 (10 ng/ml). Cells were cultured for 4 days and cell number measured using a bioluminescence assay (ATPlite). Relative light units (RLU) are shown. Red arrow indicates the combination (STI) selected for the screen. **B.** Varying numbers of CD34^+^ cells/well and varying concentrations of STI were added to a 384-well dish; cells were cultured for 4 days and cell number measured by bioluminescent assay. Red arrow indicates optimal cell number (1000 cells) and concentration of STI (12.5 ng/ml SCF, 12.5 ng/ml TPO, and 1.25 ng/ml IL3; subsequently referred to as CRCY) selected for HTS. **C.** Validation of screening conditions with CR vs CRCY at two concentrations. **D.** Screen was performed with 1000 cells/well in CRCY in 384 well dishes with 32 control wells (CRCY+DMSO) per plate (lanes 2 and 23) and Selleck library compounds (test) in lanes 3-22. Cells were cultured for 4 days without medium change and cell number was assessed by ATP bioluminescent assay.

**Supplemental Figure 2:**
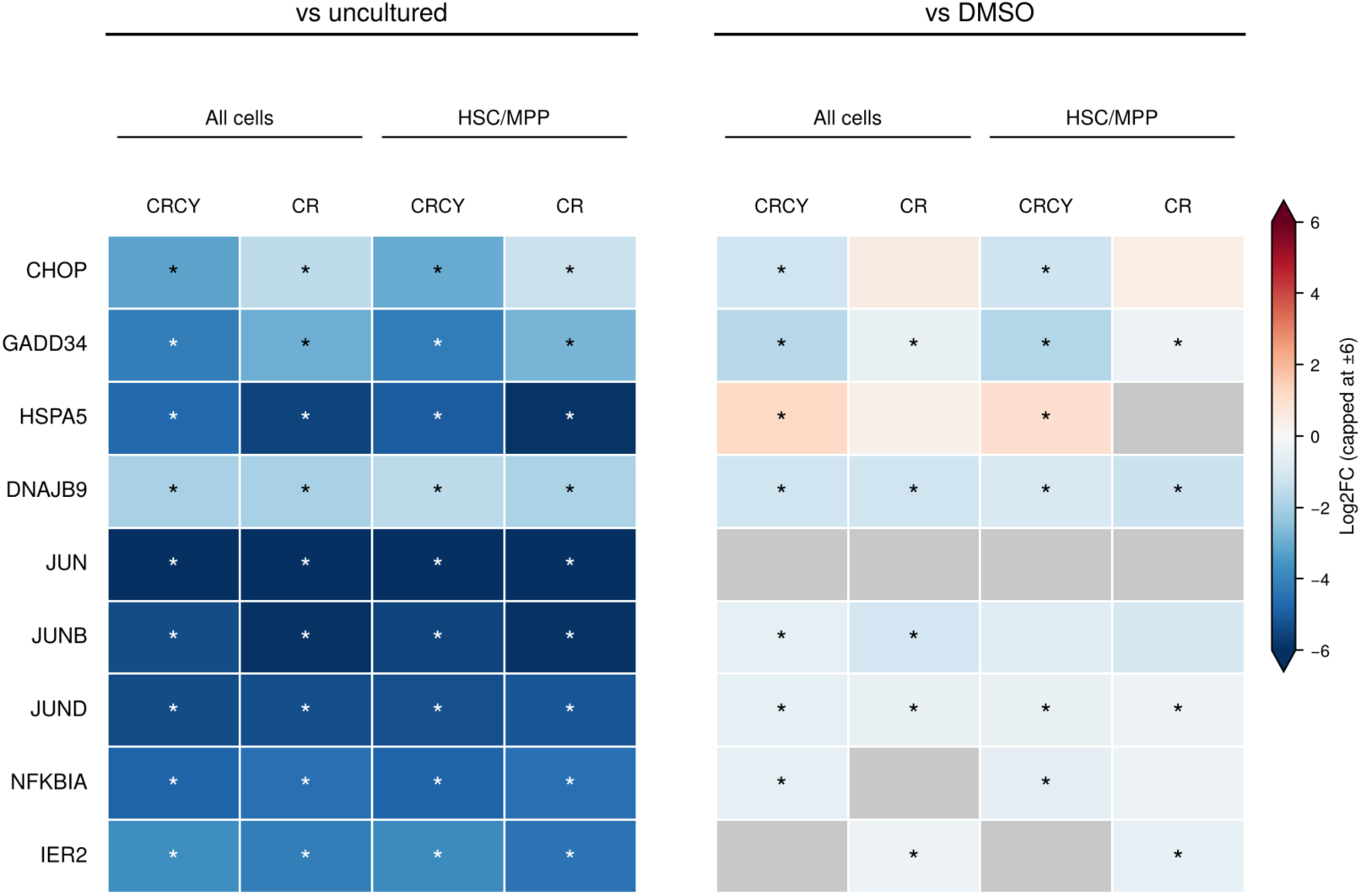
scRNA-seq analysis of UPR markers and immediate early response genes (IEGs). As in Figure 1, HSC enriched CD34^+^CD38^-^CD45RA^-^CD90^+^ cells sorted from CD34+ UBC were cultured in basal medium (Stem-span) with vehicle (DMSO), CR, or CRCY for 2 days and then subjected to 10X Genomics scRNA-seq along with freshly thawed cells from the same donor (day 0 = uncultured). Log2 fold change for CRCY or CR vs uncultured and CRCY or CR vs basal medium with DMSO (DMSO) in the CD34^+^CD38^-^CD45RA^-^CD90^+^ population (all cells) or in the HSC/MPP cluster is shown with values truncated at ±6. An asterisk indicates adjusted p ≤ 0.05, although all but one cell that achieved significance had p < 0.001. Gray indicates nonadjusted p > 0.05 or not in table because gene was below FindMarkers thresholds (|log2FC| < 0.25 or expressed in < 10% of cells). The data show marked reduction in UPR markers and IEGs compared to uncultured cells, consistent with known elevation of stress markers in freeze/thawed and FACS sorted cells. Most UPR markers and IEGs are reduced in CR, CRCY, or CRCE when compared to cells cultured in basal medium+vehicle (DMSO), although the low number of cells in the DMSO group limited statistical power.

**Supplemental Figure 3:**
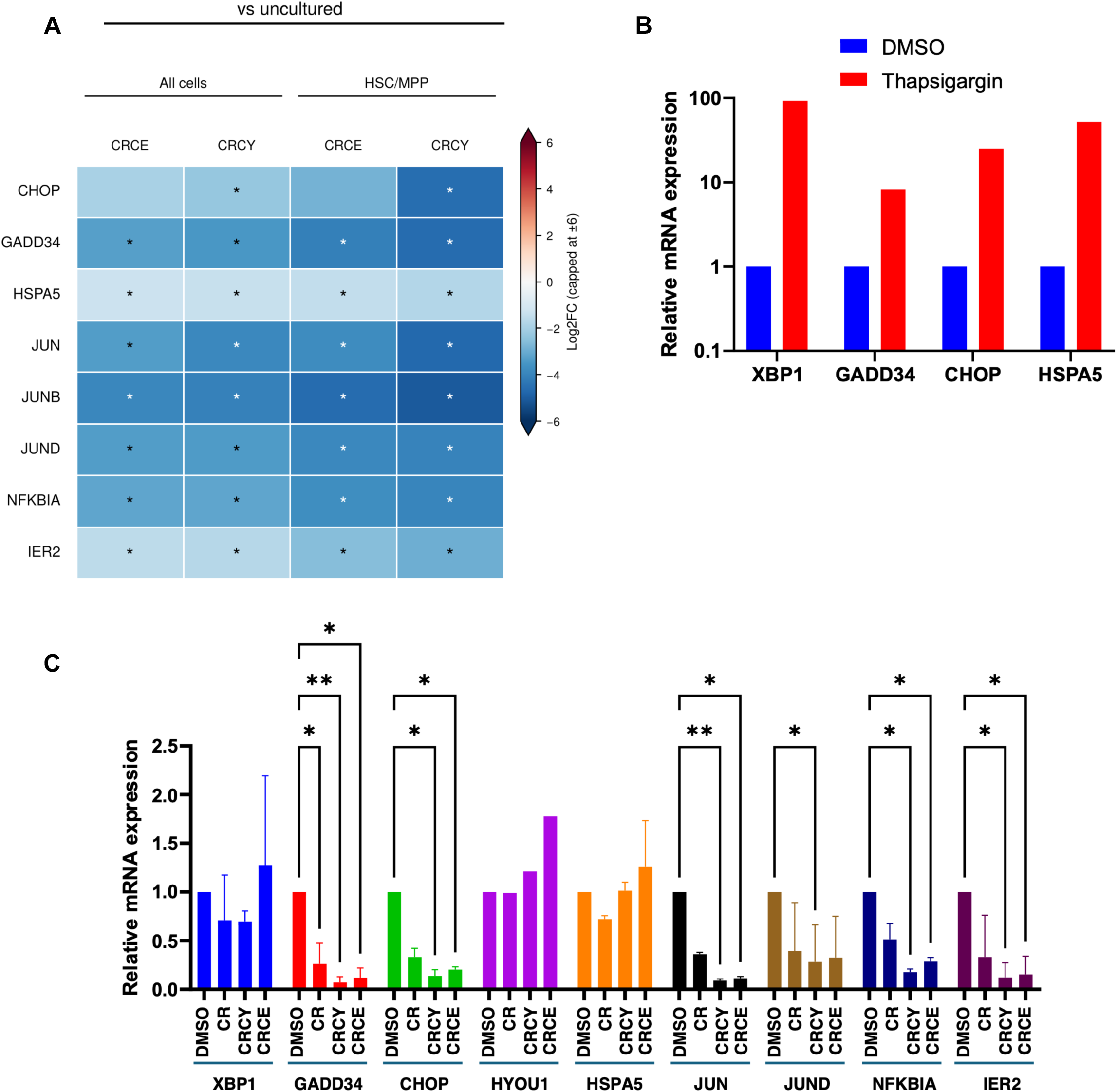
Expression of UPR markers and immediate early response genes (IEGs). **A.** scRNA-seq: As in Figure 1 and 3, HSC enriched CD34+CD38-CD45RA-CD90+ cells sorted from CD34+ UBC were cultured in basal medium with vehicle (DMSO), CRCY, or CRCE for 2 days and then subjected to 10X Genomics scRNA-seq along with day 0 (uncultured) cells from the same donor. Log2FC is shown. An asterisk indicates adjusted p ≤ 0.05 (although most p values were < 0.001). As in Supplemental Figure 2, gray indicates nonadjusted p > 0.05 or below FindMarkers thresholds (|log2FC| < 0.25 or expressed in < 10% of cells). These data confirm marked reduction in UPR markers and IEGs compared to uncultured cells, consistent with known elevation of stress markers in freeze/thawed and FACS sorted cells. Most UPR markers and IEGs are also reduced when compared to cells cultured for 2 days in basal medium+vehicle (DMSO), although the low number of cells in the DMSO group limited statistical power and detection of several markers. **B.** Positive control for UPR markers: A group of CD34^+^ cells was cultured for 22 hours in basal medium+vehicle (DMSO) and treated with 300nM thapsigargin for 2 hours prior to collection at 24h. XBP1 splicing and expression of CHOP, GADD34, and HSPA5 were assessed by RT-qPCR. As expected, thapsigargin activated IRE1/XBP1, PERK/ATF4, and ATF6 pathways. **C.** CD34+ cells at day 0 (uncultured) or cultured for 24 hours in basal medium (Stem-span) with vehicle (DMSO), CR, CRCY, or CRCE were collected for RT-qPCR. IRE1/XBP1 activation was assessed by RT-qPCR for the spliced form of XBP1. PERK/ATF4 pathway was assessed by expression of CHOP (DDIT3) and GADD34 (PPP1R15A). ATF6/XBP1 regulate HSPA5. Immediate early genes JUN, JUND, NFKBIA, and IER2 were also measured by RT-qPCR. Data are from 2 independent UCB donors. Only values with p < 0.05 are marked: * indicates p<0.05, ** p<0.01 (two way ANOVA). Compared to vehicle (DMSO) control, CHOP and GADD34 were significantly reduced in CR compared to vehicle control, and further reduced in CRCY and CRCE, while XBP1 and HSPA5 were not significantly affected. All of the IEGs were also significantly reduced in CR, CRCY, and CRCE. UPR and IEG markers were markedly reduced compared to day 0/uncultured cells (not shown), likely representing stress associated with freeze/thawing and initiation of culture.

**Supplemental Figure 4.**
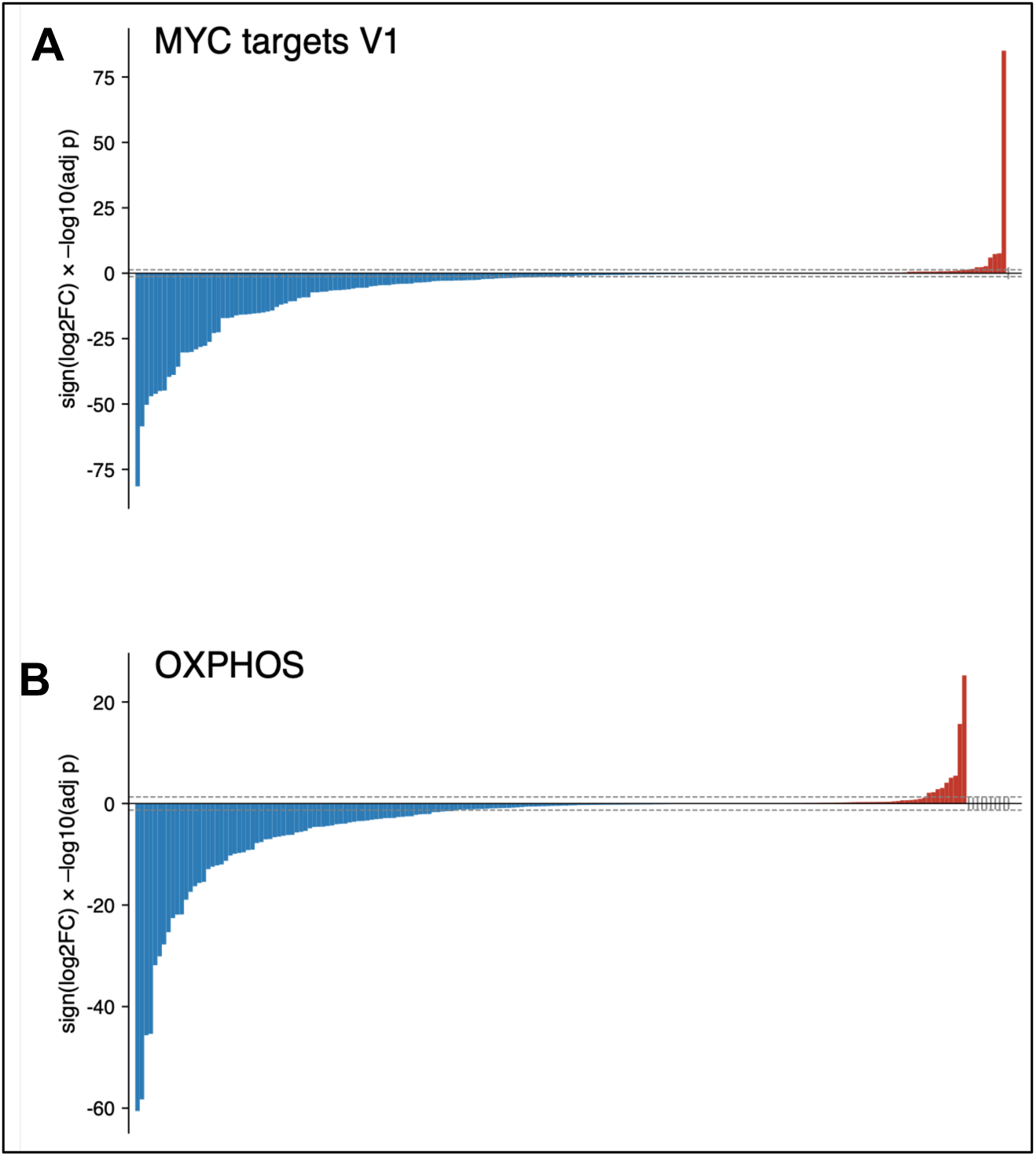
Gene-level changes in MYC and oxidative phosphorylation (OXPHOS) gene programs between CRCE and CRCY. Each bar represents one gene of the Hallmark MYC targets V1 gene set (A, 195 genes) or the Hallmark OXPHOS gene set (B, 198 genes) in CD34+CD38−CD45RA−CD90+ cells (all cells). The height of each bar is the sign of the log2 fold change multiplied by −log10 of the DESeq2 adjusted p value for CRCE vs CRCY, and genes are ordered from most negative to most positive. Blue bars are genes lower in CRCE, and red bars are genes higher in CRCE. Dashed lines mark adjusted p = 0.05. 148 of 195 MYC target genes were lower in CRCE, with 94 significantly lower and 10 significantly higher; 140 of 198 OXPHOS genes were lower in CRCE, with 73 significantly lower and 9 significantly higher. DESeq2 replicates were random splits of cells, so p values are descriptive.

**Supplemental Figure 5:**
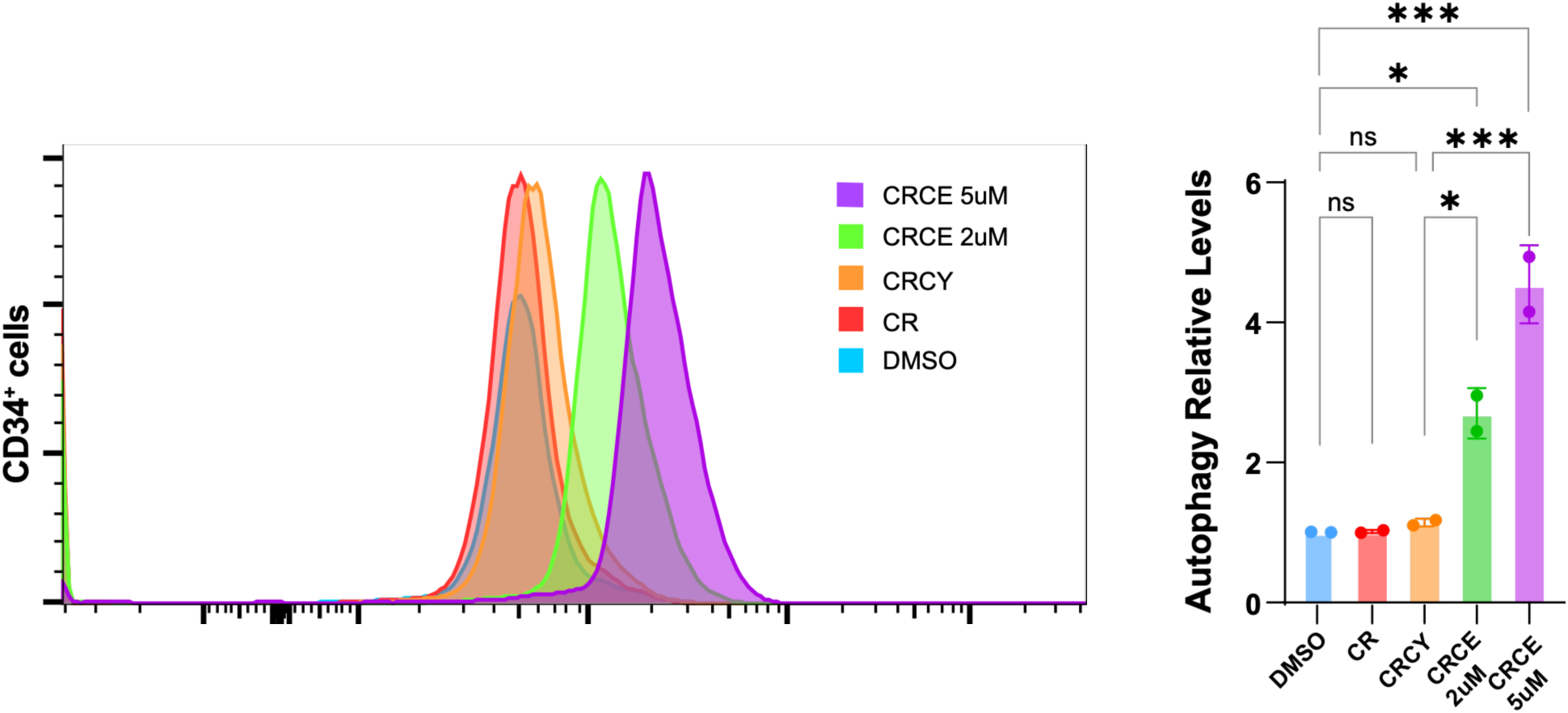
Does-dependent increase in autophagic vacuole content with 4E1RCat. CD34+ cells from UCB were cultured in control medium (DMSO), CR, CRCY, CRCE (CRCY + 2µM 4E1RCat), or CRCY + 5 µM 4E1RCat for 24 hours and autophagic vacuole content was measured by flow cytometry as in Figure 3E. A. Flow plot from one donor is representative of replicates from 3 donors. B. MFI reflecting autophagic vacuole content. 4E1Rcat increased autophagic vacuoles at 2 µM and this further increased at 5 µM. * indicates p < 0.05, *** indicates p < 0.005. Data are from one donor; a similar dose-dependent increase was observed in 2 other donors.

**Supplemental Figure 6.**
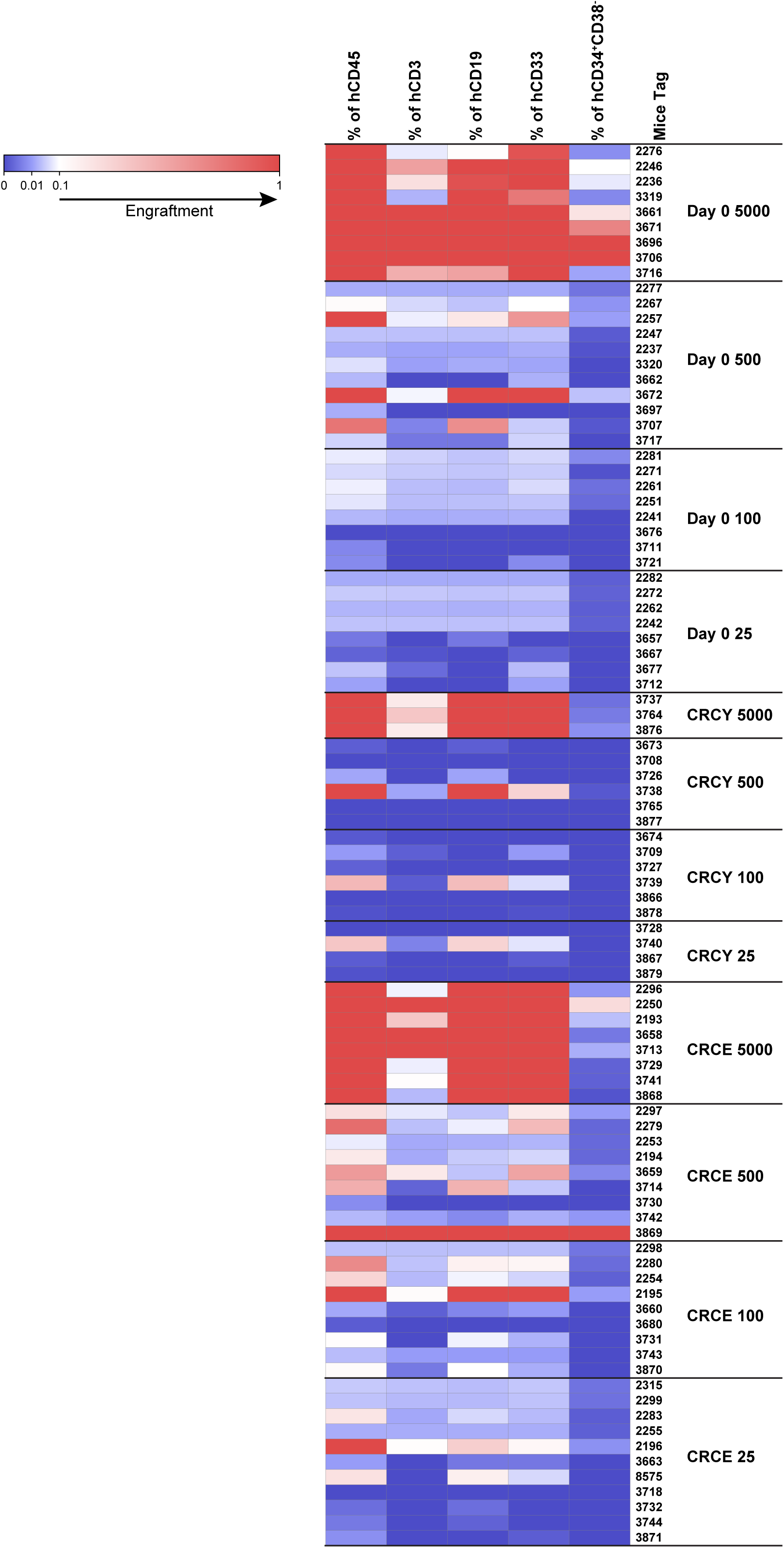
Multilineage Contribution of HSCs expanded in CRCE in primary LDA at 20 weeks. The percentage of human CD45^+^ cells, T cells (CD3^+^), B cells (CD19^+^), myeloid cells (CD33^+^), and CD34^+^CD38^-^ cells in each primary recipient mouse at 20 weeks in bone marrow are represented by a heatmap. Red indicates percentage engraftment (human cells) is > 0.1%. Blue indicates human cell engraftment < 0.1%. White indicates percentage engraftment = 0.1%. Data show results from expansion/transplants using 2 distinct mixed donor UCB samples.

**Supplemental Figure 7:**
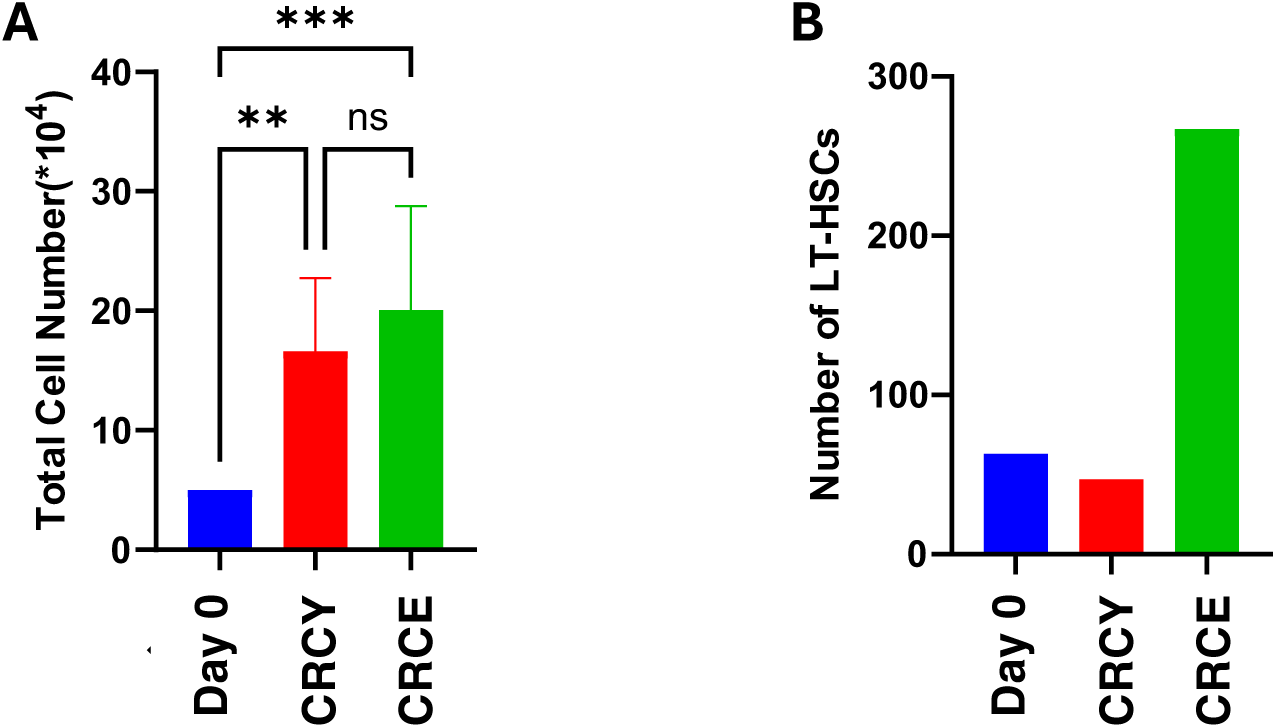
Number of CD34^+^ cells and LT-HSCs (SRCs) at Day 0 and after 7 days. A. total cell count at Day 0 (50,000 for all groups) or after 7 days of culture in CRCY or CRCE. B. Total number of LT-HSCs is shown based on frequency calculated by ELDA x cell number at day 0. The number of LT-HSCs was 63 at day 0, 47 at day 7 in CRCY, and 267 at day 7 in CRCE. Addition of 4E1RCat therefore expanded LT-HSCs 5.7-fold compared to CRCE and 4.3-fold compared to day 0.

**Supplemental Figure 8.**
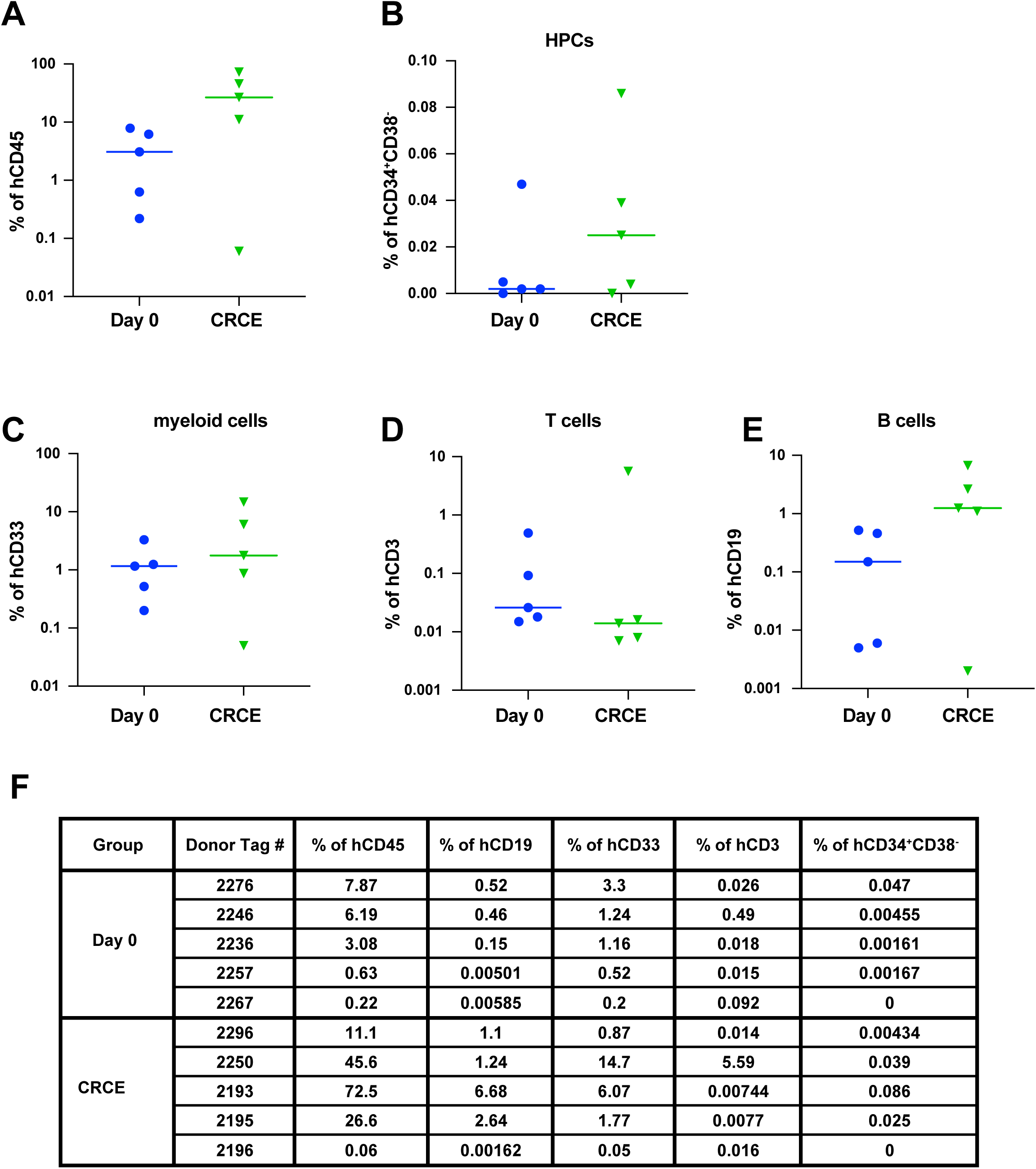
Multilineage Contribution of HSCs expanded in CRCE in primary recipients at 29 weeks.A-E. The percentage of human CD45^+^ cells, CD34^+^CD38^-^ cells, myeloid cells (CD33^+^), T cells (CD3^+^), and B cells (CD19^+^) at day 29 post-transplant in bone marrow of mice used as donors for secondary transplant. Mice had received either uncultured (Day 0) CD34^+^ cells or cells cultured in CRCE for 7 days. There was no significant difference between Day 0 and CRCE/Day 7 groups in panels A-E. **F.** Tableshowing percent engraftment of each population in each mouse at 29 weeks.

**Supplemental Figure 9.**
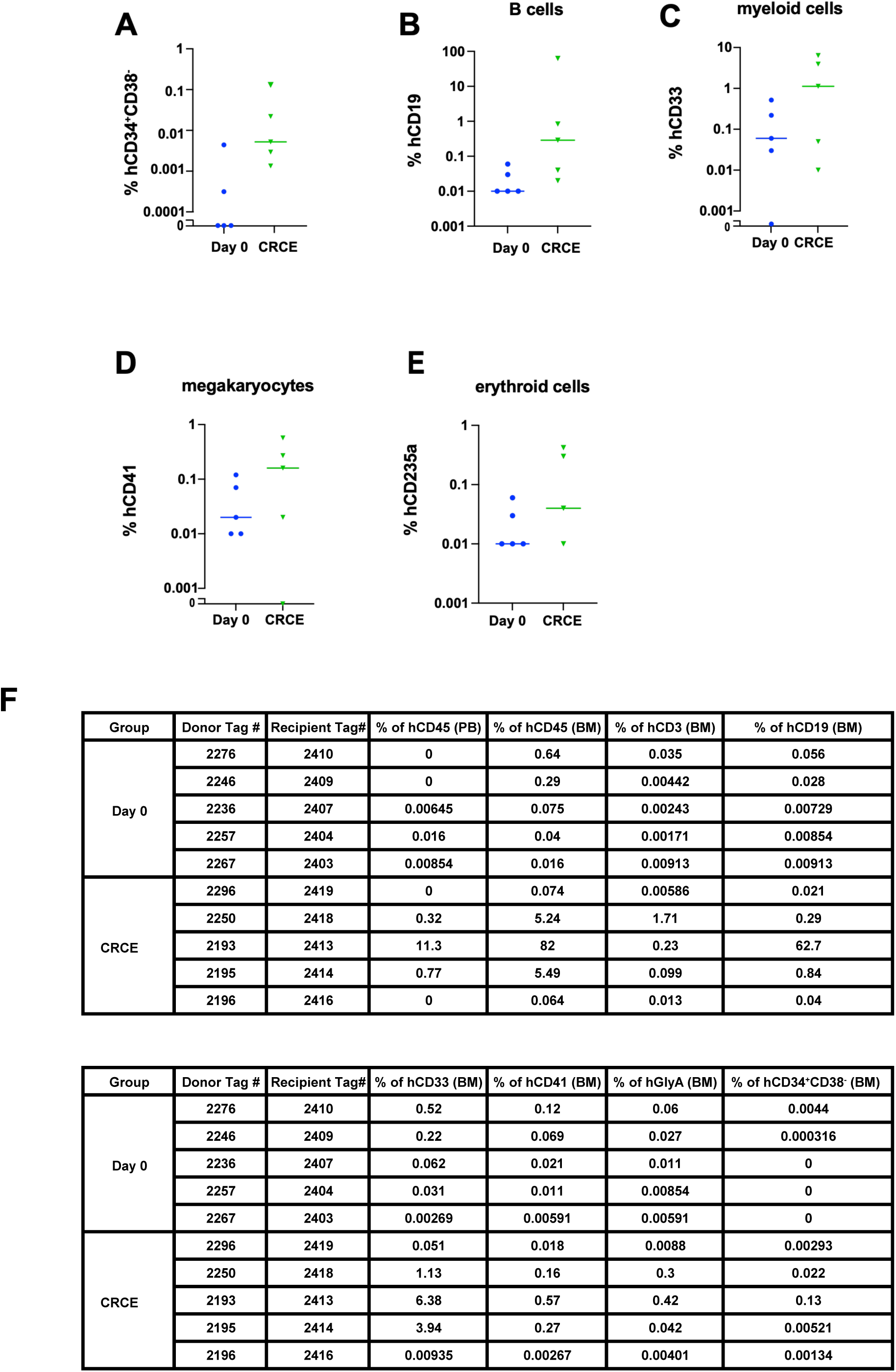
Lineage analysis of serial transplant of LT-HSCs expanded with CRCE. See. **Figure 4G-I**. Bone marrow harvested from primary recipients at 29 weeks was transplanted into conditioned secondary recipients and harvested at 18 weeks as described in Figure 4G-I. The percentage (log_10_ scale) of human **A.** lineage negative cells (CD34^+^CD38^-^), **B.** B cells (CD19^+^), **C.** myeloid cells (CD33^+^), **D.** megakaryocytes (CD41^+^), and **E.** erythroid cells (CD235a^+^) in the bone marrow (18 weeks) of secondary recipients. Samples with 0% engraftment are shown at the baseline (with broken y-axes). **F.** Table showing per mouse percentage of human CD45^+^ cells in peripheral blood (PB) at 16 weeks and in bone marrow (BM) at 18 weeks. The percentage of human CD45^+^ T cells (CD3^+^), B cells (CD19^+^), myeloid cells (CD33^+^), megakaryocytic cells (CD41^+^), erythroid cells (GlyA^+^) and CD34^+^CD38^-^ cells is also shown in the table.

**Supplemental Figure 10:**
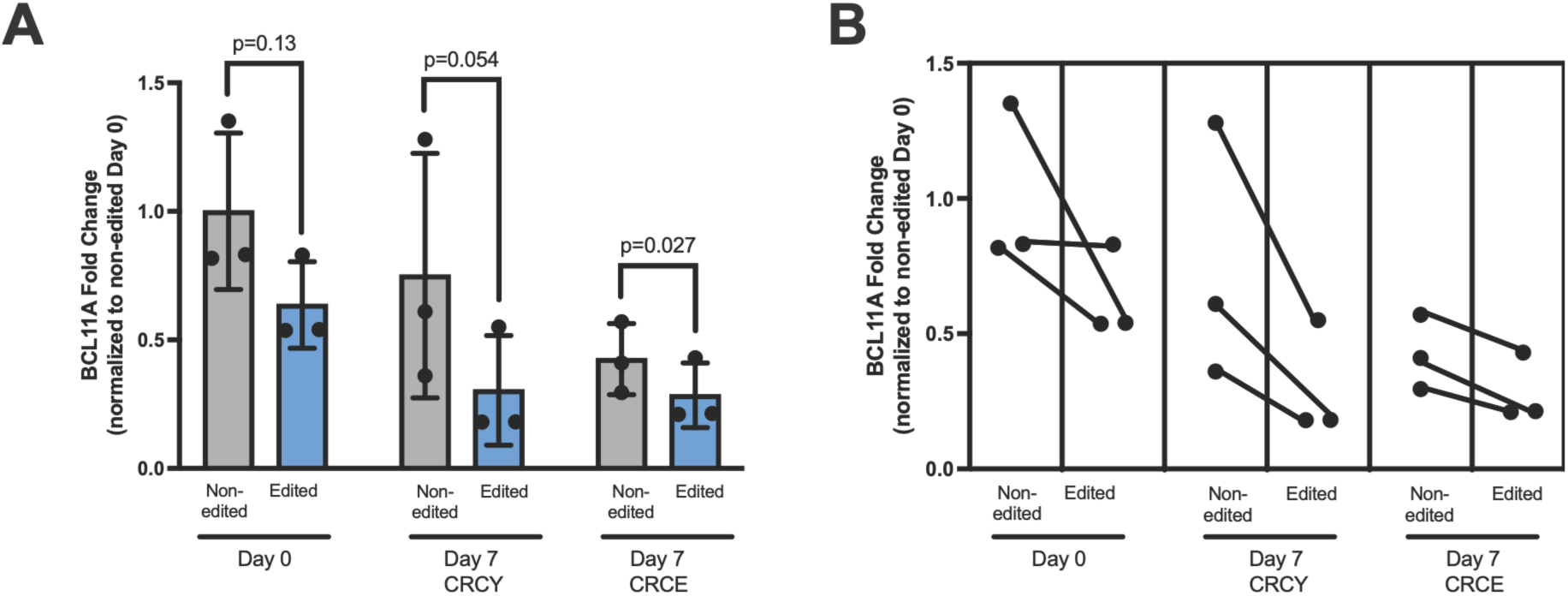
Pairwise comparison of *BCL11A*+58 enhancer editing in healthy donor-derived CD34^+^ cells expanded ex vivo. BCL11A expression data from Figure 6E was compared pairwise for samples with matched edited and non-edited data in uncultured (Day 0) cells or cells cultured for 7 says in CRCY or CRCE. A. The reduction of *BCL11A* transcript levels in *BCL11A*+58-edited samples compared to unedited samples was statistically significant in the CRCE group (*p* = 0.027) and approached significance in the CRCY group (*p* = 0.054; one-tailed paired t-test). B. Paired dot plot identifying *BCL11A* transcript levels for each pair of samples in panel A (unedited vs *BCL11A*+58 edited). *BCL11A* transcript levels were normalized to Day 0 non-edited sample for each donor; N = 3 independent donors.

**Supplemental Figure 11.**
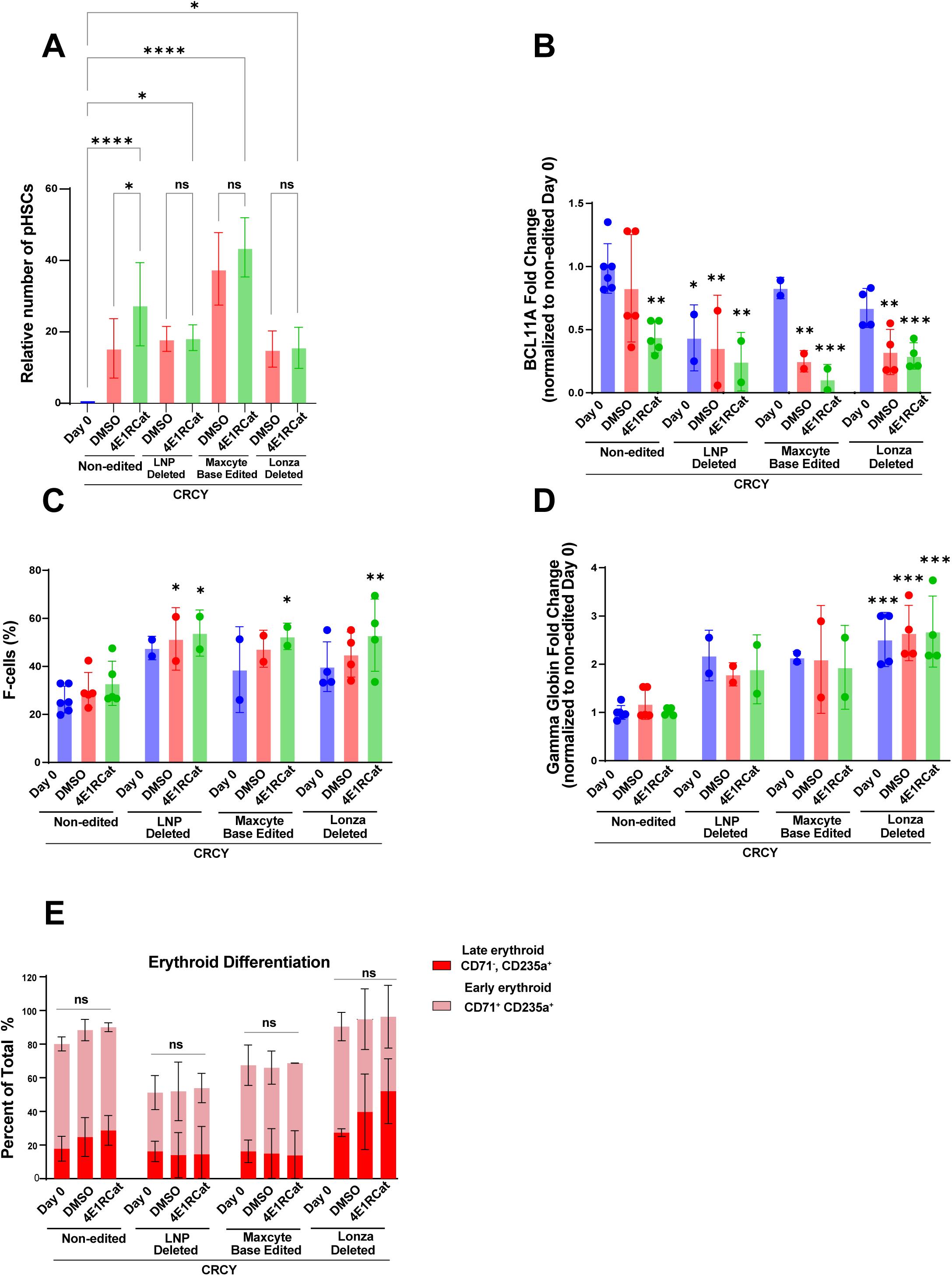
Expansion of adult HSCs after editing with conventional CRISPR/Cas9 (Lonza-EP), base editing (CBE-EP), or LNPtransduction. Mobilized adult CD34^+^ cells were edited by CRISPR/Cas9 targeting of the *BCL11A+58* enhancer to compare conventional CRISPR/Cas9 using electroporation (Lonza-EP), as in Figure 4, to transduction of Cas9/gRNA RNPs with lipid nanoparticles (LNP) and to transduction of aTadA-based cytosine base editor using MaxCyte EP(CBE-EP).After transduction, cells were cultured in CRCYwith vehicle (indicated as “DMSO”) or CRCYwith 4E1RCat (indicated as “4E1RCat”) as in Figure 4. **A.** The number of pHSCs (CD34^+^CD38^-^CD45RA^-^CD90^+^CD49f^+^) at Day 7 relative to Day 0 is shown. Expansion of edited pHSCs (compared to day 0) is similar to unedited cells, and therefore not impaired by any of the above editing approaches. The data for non-edited and Lonza-EP represent the mean of replicates from 4-6 adult donors; the data for Maxcyte and LNP are from 2 adult donors. * indicates *p* < 0.05, ** indicates *p* < 0.01, *** indicates *p* < 0.001, **** indicates *p* < 0.0001. NS, not significant (one-wayANOVA)**. B-E.** CD34^+^ cells from Day 0 (unedited), immediately after transduction (electroporation or exposure to LNPs) with *BCL11A+58* sgRNA(Day 0 edited), or after transduction and culture in CRCY with or without 4E1RCat (Day 7 edited), were differentiated in three-phase erythroid culture and collected at Day 13 for detection of **(B)** *BCL11A* mRNAby RT-qPCR (normalized to Day 0 non-edited), demonstrating reduction in *BCL11A* expression in all edited groups cultured in either CRCY or CRCY+4E1RCat compared to unedited, day 0 cells; (**C)** HbF expression (F-cells) by flow cytometry (as percent of CD71+CD235a+ erythroid cells), showing an increase in % F-cells in cells cultured in CRCY+4E1RCat after editing by Lonza-EP, CBE-EP, or LNP. Importantly, culture in CRCY±4E1RCat did not impair induction of F-cells; (**D**) *HBG* mRNAexpression by RT-qPCR (normalized to Day 0 non-edited), again showing that culture in CRCY or CRCY+4E1RCat does not impair induction of HBG. For panels B-D, **\*** indicates p < 0.05, ** indicates p < 0.01, and *** indicates p < 0.001 as compared to Day 0 non-edited control (one-way ANOVA). (**E**) Erythroid differentiation by flow cytometry (as percent of total cells) is not impaired by culture in CRCY or CRCY+4E1RCat. NS indicates not significant within each editing condition group (two-way ANOVA).

**Supplemental Figure 12:**
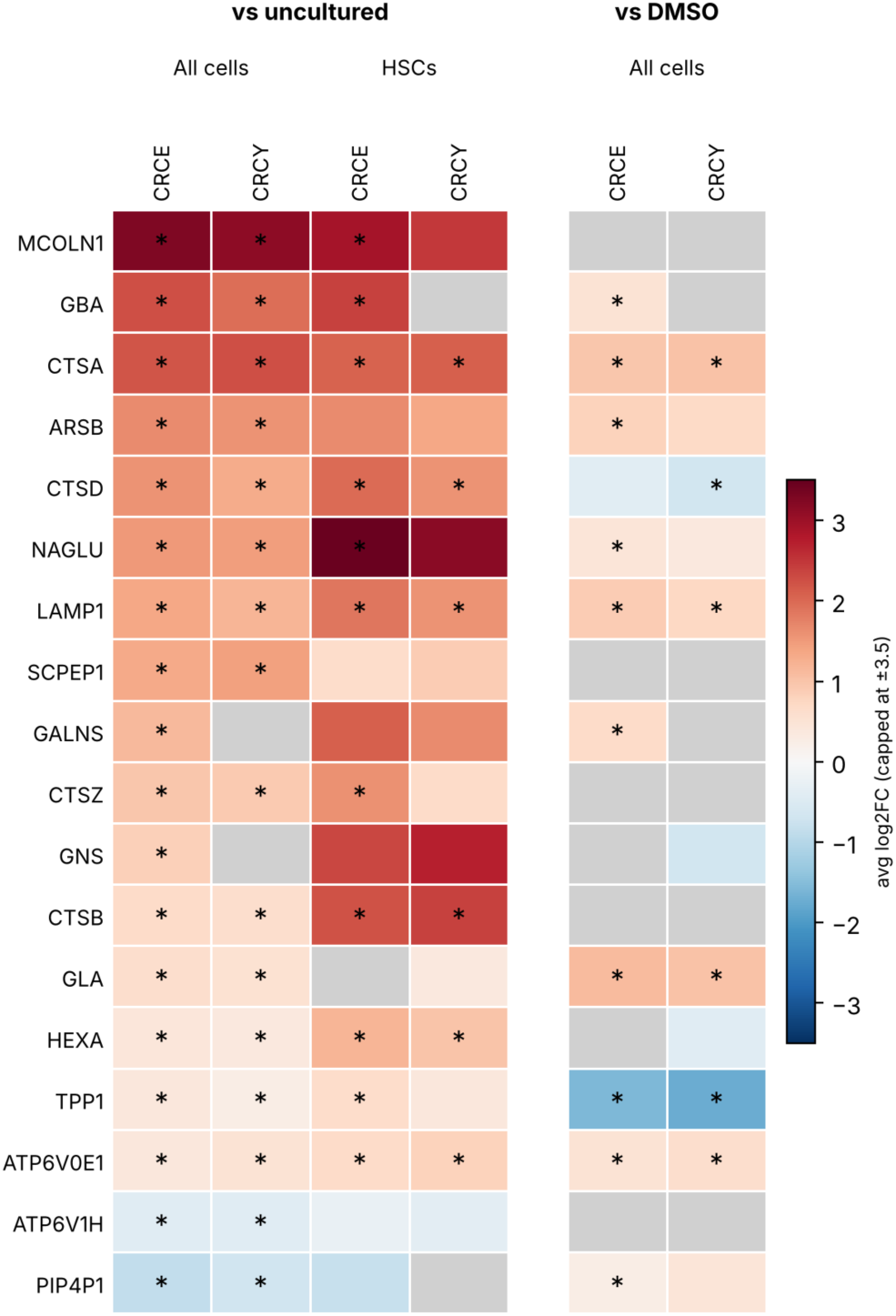
CLEAR Network genes in HSC-enriched cells and HSC/MPP cluster after culture in CRCE or CRCY: scRNA-seq analysis of TFEB target genes associated with lysosomal biogenesis and function (Taken from Sardiello et al Science 2009 PMID 19556463). HSC-enriched CD34^+^CD38^-^CD45RA^-^CD90^+^ cells sorted from CD34+ UBC were cultured in CRCY or CRCE for 2 days and then subjected to scRNA-seq along with freshly thawed and FACS sorted cells from the same donor (day 0 = uncultured), described in detail in Figure 3 and Supplemental Figure 3. Log2[fold change] (Log2FC) for CRCY or CRCE vs uncultured in the CD34^+^CD38^-^CD45RA^-^CD90^+^ population (all cells) or in the HSC/MPP cluster is shown with values truncated at ±3. Log2FC vs basal medium+vehicle (DMSO) for the CD34^+^CD38^-^CD45RA^-^CD90^+^ population is also shown. An asterisk indicates adjusted p ≤ 0.05. Gray indicates nonadjusted p > 0.05 or not in present in the table because it was below FindMarkers thresholds (|log2FC| < 0.25 or expressed in < 10% of cells). The data show an increase in most of the listed CLEAR network genes compared to uncultured cells in both the total HSC-enriched population (all cells) and the HSC/MPP cluster. A similar trend was observed for a subset of these genes when comparing CRCE and CRCY to cells cultured in basal medium+vehicle (DMSO), but the low number of cells in the DMSO group and in the HSC/MPP cluster limited statistical power.

**Supplemental Figure 13:**
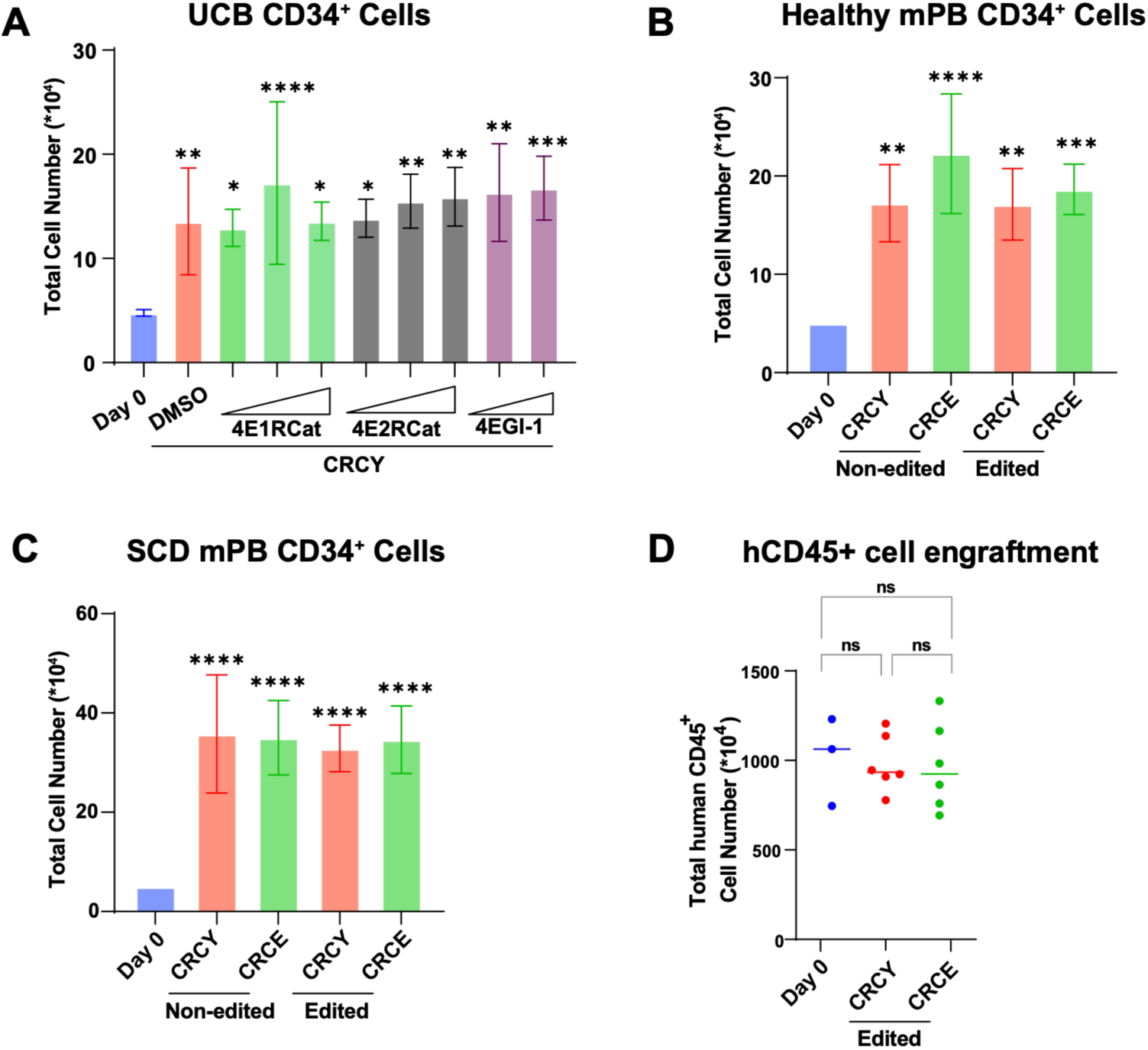
Total CD34^+^ cell counts before (Day 0) and after (Day 7) culture in CRCY and CRCE. A. From Figure 2, expansion in CRCY with vehicle (DMSO) or increasing doses of 4E1RCat, 4E2RCat, or 4EGI-1. B, C. From Figure 6, mobilized peripheral blood (mPB) CD34^+^ cells at Day 0 or day 7 culture of non-edited vs edited cells cultured in CRCY or CRCE. D. From Figure 7, absolute number of engrafted human CD45+ cells. Mean for Day 0 = 1.01 x 10^7^ hCD45^+^ cells/mouse; day 7 CRCY = 0.98 x 10^7^ hCD45^+^ cells/mouse; day 7 CRCE = 0.97 x 10^7^ hCD45^+^ cells/mouse. (* indicates p < 0.05; ** p < 0.01, *** p < 0.005, **** p < 0.001; “ns”: not significant; for all panels: one-way ANOVA).

**Supplemental Figure 14.**
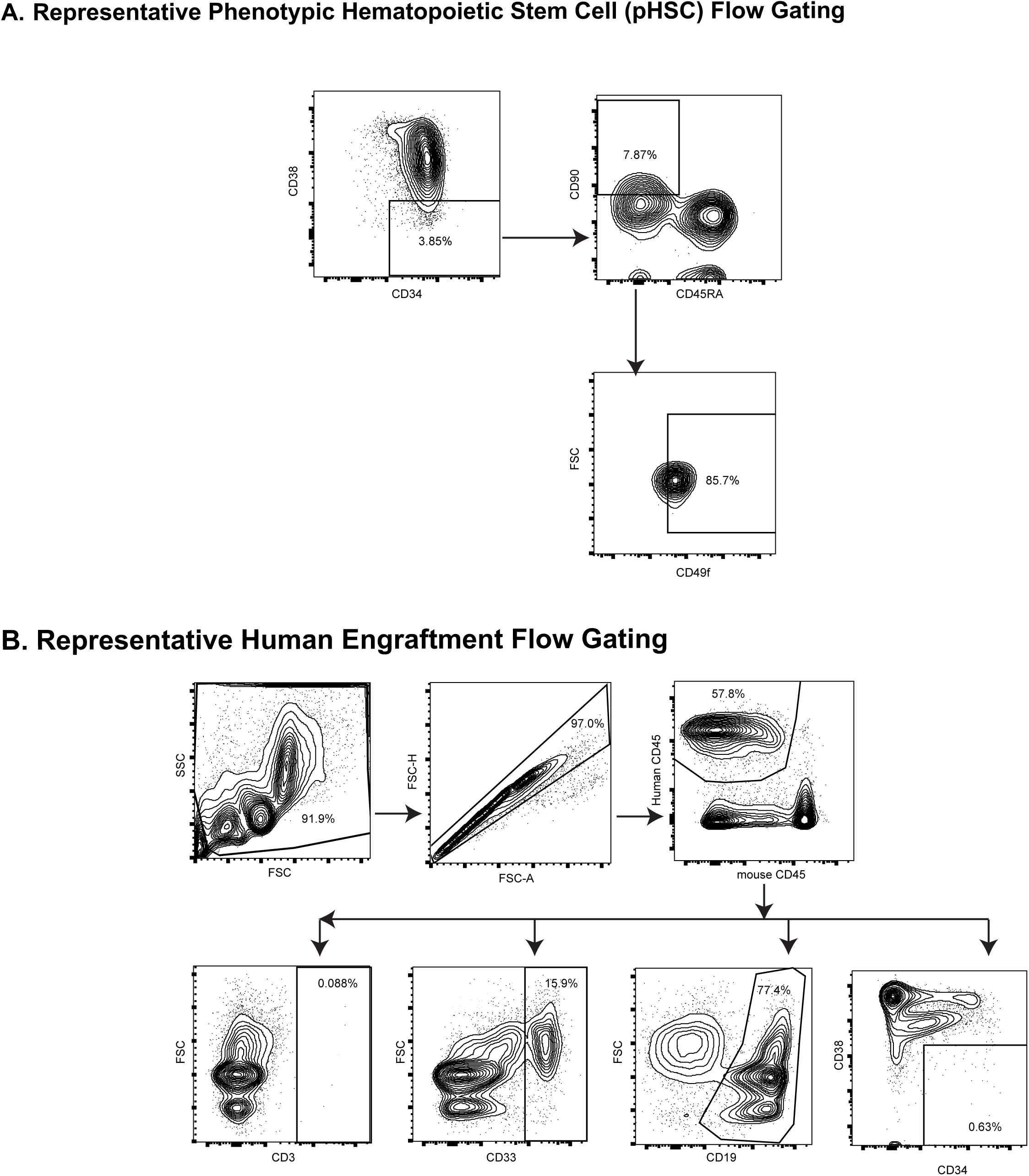
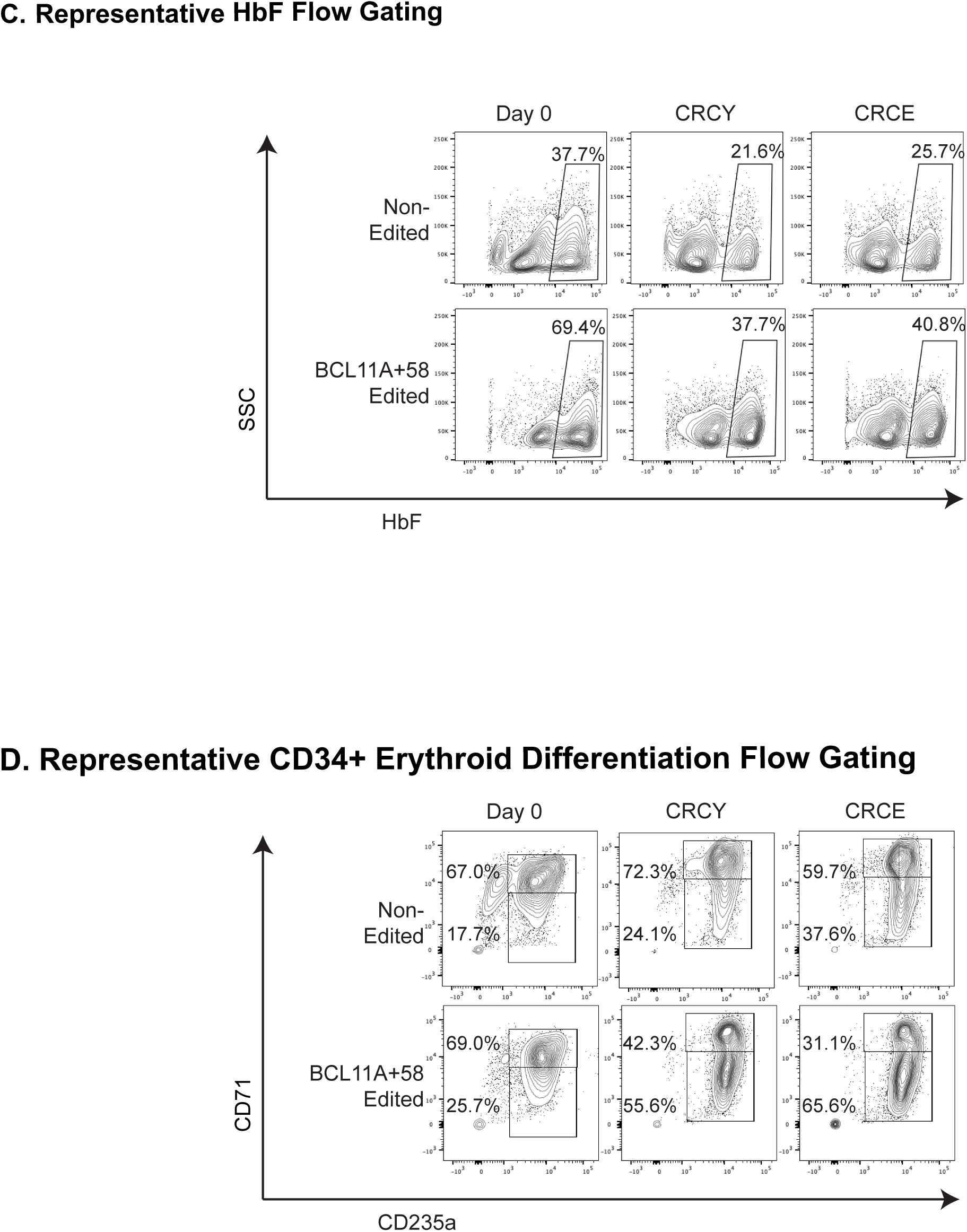
Gating Strategies

## Supplemental Methods

### Antibodies for flow cytometry

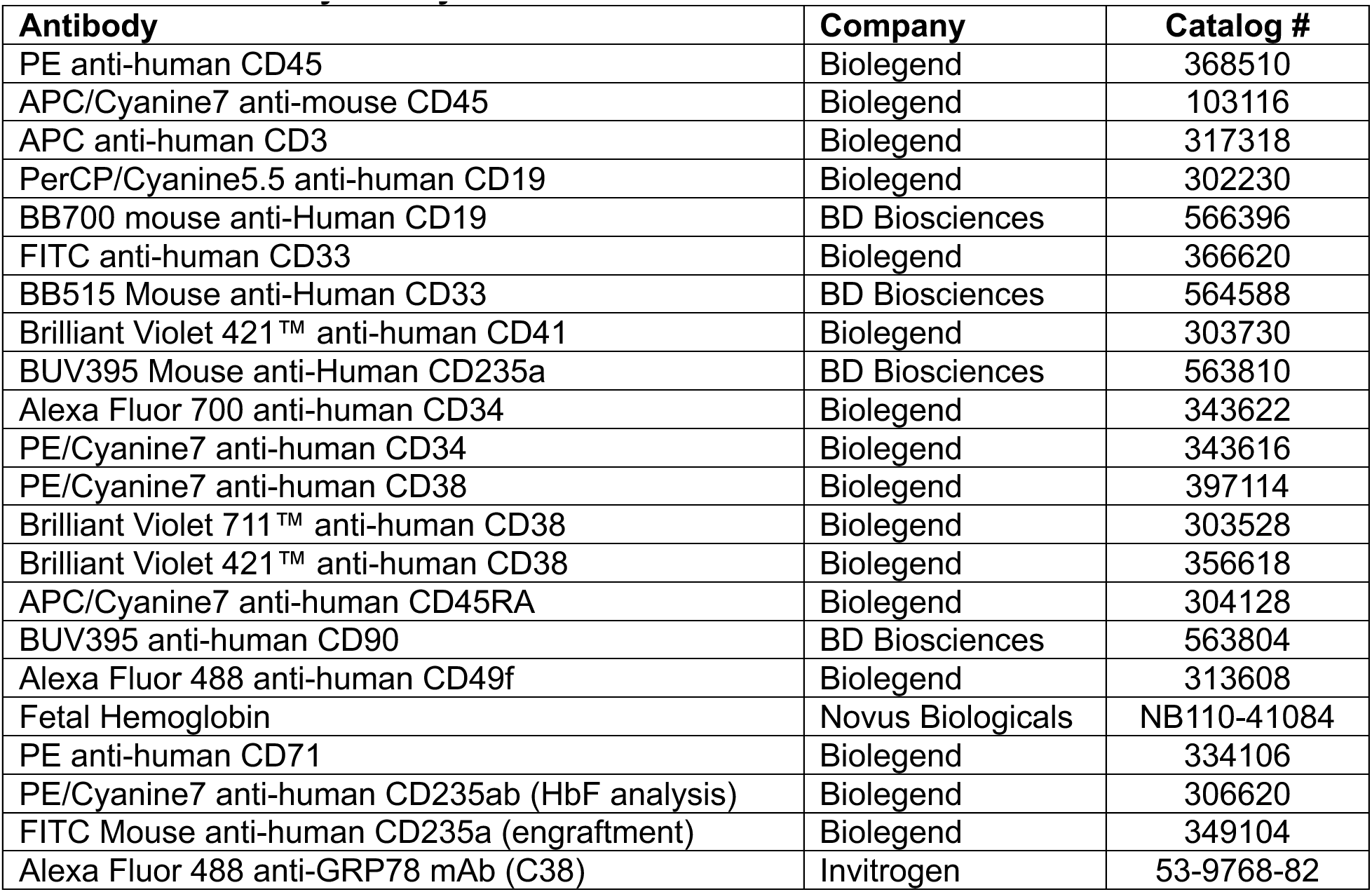

### Human umbilical cord blood CD34^+^ cell culture

Human CD34^+^ cells from umbilical cord blood pooled from 10 donors per vial were obtained from the StemCell Technologies (CAT# 70008) and cultured in StemSpan™ SFEM II (StemCell Technologies). CHIR99021 and rapamycin (Cayman Chemical) reconstituted in DMSO were added to final concentrations of 3 μM (CHIR99021) and 5 nM (rapamycin) for all experiments (designated CR medium). Low concentration cytokine medium with CR (CRCY) included 12.5 ng/ml human stem cell factor (SCF) (PeproTech Cat# 300-07), 12.5 ng/ml human Thrombopoietin (TPO) (StemCell Technologies, Cat# 78210), 1.25 ng/ml human IL3 (StemCell Technologies Cat# 78040). 4E1RCat (Selleckchem), 4E2RCat (MedchemExpress), and 4EGI-1(Selleckchem) were reconstituted in DMSO and used at the concentrations indicated. CD34^+^ cells were distributed into 96-well U-bottom plates at 50,000 cells per well with 200 μl medium. Except as described for the HTS, one-half volume of medium was replaced every other day. After 7 days (37°C, 5%CO_2_), the total culture product was harvested, and cells were washed and detected by flow cytometry or transplanted into NSG mice.

### Culture of mobilized peripheral blood CD34^+^ cells

Human CD34^+^ cells from peripheral blood of healthy adult donors were obtained from the Cooperative Centers of Excellence in Hematology Core at the Fred Hutchinson Cancer Center. Cells were thawed and cultured in CRCY±4E1RCat as described above. For erythroid differentiation, an established 3-phase erythroid culture system was used^1^. For Phase I, cells were cultured for 8 days in Iscove’s Modification of DMEM (IMDM) (Mediatech, #MT10016CV) supplemented with 100 ng/mL human SCF (Peprotech, #300-07), 1 ng/mL IL-3 (Peprotech, #200-03), 3 units/mL erythropoietin (Amgen, #55513-144-10), 200 μg/mL holo-transferrin (Sigma, #T4132), 5% human AB serum (Sigma, #H4522), 2% penicillin/streptomycin (ThermoFisher, #15140122), 10 μg/mL heparin (Sigma, #H3149), and 10 μg/ml insulin (Sigma, #I9278). IL-3 was then withdrawn (Phase 2) and cells were cultured for 5 days. For Phase III, the cells were cultured for 2 days with IMDM supplemented with 3 units/mL erythropoietin, 2% penicillin/streptomycin, 1 mg/mL holo-transferrin, 10 μg/ml insulin, 5% human A/B plasma and 10 μg/mL heparin.

### Transplantation into NSG and NBSGW mice

Transplants into non-obese diabetic severe combined immunodeficient IL-2Rγ^null^ (NSG) and NOD.Cg-*Kit^W-41J^ Tyr*^+^ *Prkdc^scid^ Il2rg^tm1Wjl^*/ThomJ (NBSGW) mice were performed by the Stem Cell and Xenograft Core Facility at Perelman School of Medicine at the University of Pennsylvania). All animal experiments were performed in accordance with guidelines approved by the Institutional Animal Care and Use Committee (IACUC) at the University of Pennsylvania. Limiting dilution assay transplants were performed on cells from 3 independent mixed donor human UCB samples and data were combined for analysis. Transplant recipients were 8-to 10-week-old females. For primary transplants, Day 0 or cultured human CD34^+^ cells were injected into NSG mice conditioned with 30mg/kg busulfan 24 hours prior by intravenous (IV) injection. For Day 0 cells, injected cell doses were 5000, 500, 100, and 25 cells per mouse. For Day 7 samples, cell doses were based on the Day 0 equivalent^2^ or starting number of cells (a proportion of the culture at Day 7 corresponding to 5000, 500, 100, or 25 cells at Day 0). Bone marrow was collected by aspiration at 20 weeks and red blood cells were lysed with Ammonium Chloride Solution (STEMCELL Technologies). Mononuclear cells were stained with PE anti-human CD45 antibody, APC/Cyanine7 anti-mouse CD45 antibody, APC anti-human CD3 antibody, PerCP/Cyanine5.5 anti-human CD19 antibody, FITC anti-human CD33 antibody, Alexa Fluor 700 anti-human CD34 antibody, and PE/Cyanine7 anti-human CD38 antibody, and human cell engraftment and multilineage reconstitution were assessed by flow cytometry. The frequency of HSCs was calculated using Extreme Limiting Dilution Analysis (ELDA) software (https://bioinf.wehi.edu.au/software/elda/) from the Bioinformatics Division, the Walter and Eliza Hall Institute of Medical Research).

For serial transplantation, bone marrow was harvested from primary recipients by terminal harvest and extrusion from femurs and tibias at 29 weeks, and 10 × 10^6^ bone marrow cells were transplanted into each busulfan conditioned secondary NSG recipient. After 18 weeks, bone marrow was collected and analyzed with PE anti-human CD45 antibody, APC/Cyanine7 anti-mouse CD45 antibody, APC anti-human CD3 antibody, BB515 Mouse anti-Human CD33, BB700 mouse anti-Human CD19, Brilliant Violet 421™ anti-human CD41 antibody, BUV395 Mouse anti-Human CD235a antibody, PE/Cyanine7 anti-human CD34 antibody, and Brilliant Violet 711™ anti-human CD38 antibody. Analyses were performed on LSRFortessa flow cytometers.

To assess engraftment of CRISPR-edited, mobilized adult CD34^+^ cells, freshly isolated (Day-1) cells or cells cultured for 7 days in CRCY+DMSO (vehicle control) or CRCE after electroporation were transplanted into NBSGW mice pre-conditioned with 15mg/kg busulfan 24 hours prior to injection. NBSGW mice were selected because they support improved erythroid engraftment compared to NSG mice. An equal starting cell number was used for Day-1 and Day 7 injections. Thus, for Day-1 cells, 100,000 cells were injected and for Day 7 cells, a volume of the culture corresponding to 100,000 cells at Day-1 sample was injected. Bone marrow was harvested for analysis at 20 weeks.

### Fluorescence activated cell sorting (FACS) and flow cytometric analysis

For phenotypic HSC (pHSC) detection, freshly thawed (Day 0) or cultured human CD34^+^ cells are incubated with Alexa Fluor 700 anti-human CD34 antibody (see antibody table), Brilliant Violet 421™ anti-human CD38 antibody, APC/Cyanine7 anti-human CD45RA antibody, Alexa Fluor 488 anti-human CD49f antibody, BUV395 anti-human CD90, and LIVE/DEAD™ Fixable Aqua Dead Cell Stain (CAT# L34965, ThermoFisher Scientific). Analyses were performed on LSRFortessa flow cytometers (Becton Dickinson). Data were analyzed using FlowJo 10.9.0. Sorting was performed on the BD Influx™ cell sorter.

### Assessment of rate of translation

Human CD34^+^ cells were cultured as described above for 24 h, and then O-propargylpuromycin (OP-Puro; Click-iT™ Plus OPP Alexa Fluor™ 647 Protein Synthesis Assay Kit, ThermoFisher Scientific) was added (10 μM) to the medium for an additional 60 min. Cells were washed with PBS and stained with surface markers indicated. BD Cytofix/Cytoperm™ Fixation/Permeabilization Kit (CAT# 554714) were used for the fixation and permeabilization of cells. Cells were incubated in 250 µl BD Fix/Perm solution for 20 min covered on ice. Then cells are incubated in 1ml of 1x BD Perm/Wash buffer at room temperature for 15 minutes. The azide-alkyne cycloaddition was performed using the Click-iT™ Plus OPP Alexa Fluor™ 647 Protein Synthesis Assay Kit as directed by the manufacturer. After the 30-min reaction, cells were washed in PBS supplemented with 2.5 % FBS, then stained for surface markers indicated and analyzed by LSRFortessa flow cytometers (Becton Dickinson). Data were analyzed using FlowJo 10.9.0.

### Flow cytometric detection of autophagic vacuoles

Human CD34^+^ cells from UCB were cultured in StemSpan medium + vehicle (DMSO), CR, CRCY, CRCE (2 µM 4E1RCat), or CRCY + 5µM 4E1RCat for 24 hours and then analyzed by flow cytometry with a commercial kit (Abcam #ab139484) to label autophagic vacuoles.

### Colony-forming unit (CFU) assay

For CFU assays, 100 day 0 or cultured CD34^+^ cells were mixed with 1 ml semi-solid methylcellulose medium MethoCult™ H4435 Enriched (STEMCELL Technologies) by shaking vigorously for one minute, incubated for 10 minutes to dissipate bubbles, and plated in 6-well SmartDish™ plates (STEMCELL Technologies) using a blunt-end needle and syringe. Sterilized water or phosphate buffered saline were added to empty wells to provide humidity. After 14 days, colonies were counted by STEMvision™ Hematopoietic Colony Counter (STEMCELL Technologies).

### CRISPR–Cas9 RNP Electroporation (Lonza-EP)

CRISPR-Cas9 RNP electroporation was performed as previously described^1,3^. Briefly, for in vitro studies shown in Supplementary Figure 6A-B, RNP complexes were assembled by combining 300 pmol 2′-O-methylated (2’O-Me) and phosphorthioate (PS) end-modified sgRNA (purchased from Synthego) and 50 pmol HiFi SpCas9 protein (IDT #1081061) and incubated at room temperature for 10 minutes. CD34^+^ cells (200,000 cells) in CRCY were electroporated 6 hours after thawing in 25 µL total volume using the P3 Primary Cell 4D-NucleofectorTM X Kit S (#V4XP-3032) on the Amaxa 4D Nucleofector (Lonza) with program DZ-100. For larger-scale electroporations used for xenotransplantation shown in Figure 4 and Supplementary Figure 6C, RNP complexes were assembled by combining 1200 pmol modified sgRNA (Synthego) and 400 pmol HiFi SpCas9 protein (IDT #1081061) and incubated at room temperature for 10 minutes. CD34^+^ cells (1.6E6 cells) in CRCY were electroporated 6 hours after thawing in 100 µL total volume using the P3 Primary Cell 4D-NucleofectorTM X Kit L (#V4XP-3024) on the Amaxa 4D Nucleofector (Lonza) with program DS-130. After electroporation, cells were transferred to CRCY and allowed to recover overnight. Viable cells were then counted, centrifuged, resuspended in CRCY, placed into 96 well dishes at 50,000 cells/well in CRCY±4E1RCat, and cultured for 7 days as described in main methods section. The sgRNA sequence used to target the erythroid-specific BCL11A+58 enhancer was 5’-CTAACAGTTGCTTTTATCAC-3’.

### Cytosine Base Editing by Electroporation (MaxCyte CBE-EP)

Electroporation substrate (EPS) was prepared by combining TadA-derived cytosine base editor mRNA (synthesized with N1-methylpseudouridine throughout and capped with “clean cap AG” from TriLink) with end-modified sgRNA (Synthego) at a 1:1.5 (mRNA:sgRNA) ratio in Opti-MEM™ I Reduced Serum Medium (ThermoFisher Scientific #31985070) supplemented with RNasin™ Plus Ribonuclease Inhibitor (Promega #N2615, 1.2 U/µL, final). CD34⁺ cells were washed and resuspended in the same Opti-MEM/RNase inhibitor solution, then mixed with EPS to achieve a final concentration of 5 × 10⁷ cells/ml for electroporation. A total of 6 µg RNA was used per 1 × 10⁶ cells. Electroporation was performed using an ExPERT GTx (MaxCyte) with program HSC4. After electroporation, cells were transferred to CRCY and allowed to recover overnight. Cells were then counted, centrifuged, resuspended in CRCY±4E1RCat, placed into 96 well dishes at 50,000 cells/well, and cultured for 7 days as described in the main methods section. The sgRNA sequence used to target the erythroid-specific *BCL11A+58* enhancer was 5’-TTTATCACAGGCTCCAGGAA-3’.

### CRISPR–Cas9 mRNA Lipid Nanoparticle Editing (LNP)

Preclinical grade, and endotoxin free N1 methylpseudouridine modified and codon optimized CRISPR–Cas9 mRNA was purchased from RNA technologies (Montreal, Canada, OTS-EDIT-005). HPLC grade 2’-O-methyl ribonucleosides, and phosphorotiaoate modified *BCL11A* sgRNA was purchased from IDT (custom made). The mRNA and the gRNA were mixed at 1:1 ratio, and encapsulated into lipid nanoparticles using the Ignite Nanoassembler microfluidic device (Precision Nanosystems). The nucleic acid containing solution was acidified (pH 4) using sodium acetate, and mixed at 3:1 (vol:vol) with an ethanolic mixture containing SM-102 (ionizable lipids), DSCP (Helper lipid), Cholesterol (structural lipid), and DMG-PEG-2000 LNP at a lipid mole ratio of 50:10:38.5 and 1.5%. The LNP mixture was buffer exchanged using an AMICON-Ultra 15 centrifugation filter, concentrated, sterile filtered, frozen and stored at-80C prior to use^4^. LNP size, polydispersity, and surface charge were determined using dynamic light scattering on a Malvern Zetasizer Ultra Red Encapsulation. Encapsulation efficiency, and RNA content (concentration) were calculated by Quant-iT RiboGreen RNA Assay (Thermo Fisher #E11490), as described^5^. LNPs were added to CD34⁺ cells (1.5 × 10⁶ cells/mL) to achieve final RNA (Cas9 mRNA + gRNA) concentration of 1.5 µg/mL (1µg/10^6^ cells). Apolipoprotein E3 (Peprotech #350-02-500UG) was added to a final concentration of 0.1 µg/mL. Cells were cultured overnight, then counted, centrifuged, resuspended in CRCY, placed into 96 well dishes at 50,000 cells/well in CRCY±4E1RCat, and cultured for 7 days as described in methods. The sgRNA sequence used to target the erythroid-specific *BCL11A+58* enhancer was 5’-CTAACAGTTGCTTTTATCAC-3’.

### HbF and erythroid differentiation flow cytometry

HbF and erythroid differentiation analyses were performed as previously described^3^; briefly, 1.5 million cells at day 15 of culture were washed in PBS, fixed with 0.05% glutaraldehyde (Sigma #G6257), permeabilized with 0.1% Triton X-100 (Life Technologies #HFH10) and stained with AF647 HbF (Novus Biologicals #NB110-41084) at 1:200 dilution and PE-CD71 and PE-Cy7 CD235a at 1:100 dilution. Flow cytometry was carried out on a FACSCanto or LSRFortessa analyzer (Becton Dickinson) and analyzed with FlowJo 10.9.0 software.

### RT-qPCR

For RNA extraction, cells were harvested in TRIzol (ThermoFisher Scientific, #15596018) and the aqueous phase was recovered. 100% ethanol (0.7 volumes) was added to the aqueous phase and RNA was purified with the RNeasy Mini kit (Qiagen, #74106) with a DNase digestion step on the column with the RNase-free DNase set (Qiagen, #79254). RNA (100-200ng) was reverse-transcribed with iScript Reverse Transcription Supermix (BioRad, #1708841). Quantitative PCR (qPCR) was performed with cDNA from 2 ng RNA using the 2x Power SYBR Green Master Mix (Life Technologies, #4367660). Data were normalized to RPS18. PCR primers for *HBG* were forward: GAACTGCACTGTGACAAGCTG; reverse: TTTGCCGAAATGGATTGCCA and for *HBB* were forward: GCACGTGGATCCTGAGAACTT; reverse: GCGAGCTTAGTGATACTTGTGGG. *BCL11A-XL* primers were forward: CAGCGGCACGGGAAGTGGAG and reverse: CGCCCGTGTGGCTTCTCCTG. RPS 18 primers were forward: GTAACCCGTTGAACCCCATT and reverse: CCATCCAATCGGTAGTAGCG.

### High-throughput screen

The HTS screen was performed through the High-Throughput Screening Core facility at the Perelman School of Medicine. Human UCB CD34^+^ cells in CRCY (StemSpan SFEM with 3 μM CHIR99021, 5 nM rapamycin, 12.5 ng/ml human SCF, 12.5 ng/ml human TPO, and 1.25 ng/ml human IL3) were distributed into 384-well plates at 1000 cells per well where each well contained one test compound. The screen was performed with the UPENN Bioactive library, which is composed of 2240 small molecules purchased from Selleck Chemicals and was selected because it includes ∼1100 FDA approved drugs and an additional 1100 compounds with known pharmacologically activity against annotated molecular targets in current use as advanced investigational tools, chemical probes, preclinical candidates, and enzyme/receptor inhibitors identified in biomedical research literature for diverse disease indications. The library is composed of protein kinase inhibitors, G protein coupled receptor agonists and antagonists, chromatin modifying agents (e.g. HDAC, HAT, histone methyltransferase (HMT), and BET/bromodomain inhibitors), calcium, potassium, and sodium channel blockers/activators, antineoplastic agents targeting DNA repair, topoisomerases, CDKs, and other oncological targets, anti-microbial, anti-viral, protease inhibitors, protein synthesis inhibitors, and metabolic regulators. Compounds are suspended in DMSO and arrayed in columns 3 through 22 of source plates at a single concentration. Library source plates were freeze-thawed ≤10 times prior to use. The library was screened at 1µM and 0.1µM. Cells were similarly distributed into at least 16 wells/plate without test compounds as vehicle (DMSO) controls. After 4 days of culture (37°C, 5%CO_2_), cell number was measured indirectly using the Luminescence ATP Detection Assay System kit as directed by the manufacturer (PerkinElmer). Luminescence was detected by an EnVision Xcite multi-plate reader (PerkinElmer). Compounds inducing > 1.4-fold increase in CD34^+^ cells that were also > 4 standard deviation units above the mean cell number for vehicle controls in 2 replicate screens at 1µM were subjected to a third screen at 100nM.

### Single-cell RNA-seq

CD34^+^CD38^-^CD45RA^-^CD90^+^ cells were purified from CD34^+^ UCB cells by fluorescence activated cell sorting (FACS) and cultured in StemSpan SFEM with vehicle control (DMSO), CR, CRCY, or CRCE. After two days, the cultured samples and freshly thawed and sorted CD34^+^CD38^-^CD45RA^-^CD90^+^ cells (uncultured/day 0) from the same lot number/pool of cells used for culture were collected and single cells were isolated using the 10XGenomics platform. Cells were collected at 2 days of culture to capture early changes in gene expression and because cells in the vehicle (DMSO) group did not survive well in prolonged culture. cDNA libraries were prepared according to the 10X Genomics user manual using Chromium Single Cell 3’ Reagent Kits v3 (10X Genomics) through the Center for Applied Genomics, Children’s Hospital of Philadelphia. Next generation sequencing was performed by Genewiz. Cell Ranger (10X Genomics) was used to process the scRNA-seq data. Cell Ranger Count aligned the sequencing reads to the human reference genome (hg38) using STAR. The output files for the two replicates were aggregated into one gene-cell expression matrix using Cell Ranger aggr with the mapped read depth normalization option. Subsequent analysis was performed using Seurat (4.3.0.1) in R. Using Read10X function, we obtained unique molecular identifiers (UMI) for each cell in Day 0, CR, CRCY, CRCE, and control medium (DMSO). scRNA-seq was performed with cells purified from two UCB donors. For the first donor, the experimental groups were: control medium (DMSO), CR, CRCY, and day 0 cells. This analysis identified 10364 cells in the Day 0 group, 11771 cells in the CR group, 10519 cells in the CRCY group, and 4884 cells in the DMSO group. Quality control was performed to remove low quality cells. We filtered out cells that contained less than 700 unique feature counts and cells that contained more than 10% mitochondrial counts. After filtering, Day 0 contained 8571 cells, CR contained 8683 cells, CRCY contained 10282 cells, and DMSO contained 3080 cells. scRNA-seq data were deposited in the NCBI Gene Expression Omnibus^6^ and are accessible through GEO SubSeries accession number GSE248311. For the second donor (2^nd^ set), the groups were: control medium (DMSO), CRCY, CRCE, and Day 0 cells. Data for the second set are accessible through GEO SubSeries accession number GSE353932. Both groups are linked under GEO SuperSeries GSE353936.

SCTransform function (method = “glmGamPoi”) was applied to normalize and scale the gene expression within each condition. The effect of mitochondria was removed using vars.to.regress = c(“percent.mt”). SelectIntegrationFeatures function (nfeatures = 3000) was performed to select features for integration. PrepSCTIntegration function was used to prepare SCTransform gene expression for integration and FindIntegrationAnchors function (reference = UC) and IntegrateData function detected anchors and integrated cells from each condition. RunPCA function was used to generate principal component analysis (PCA) and the RunUMAP function was used to reduce the dimensions of the integrated dataset into 2-D space. The FindNeighbors function was performed to find k-nearest neighbours (KNN) for each cell, and FindCluster function was performed to group cells basis of KNN into cluster using the Louvain algorithm (resolution = 0.1,0.3,0.5,0.8,1,2,3), and resolution = 0.3 was picked in clustering cells. Cells were annotated based on gene markers defined by Zheng et al^7^ and the HumanPrimaryCellAtlasData library. After running PrepSCTFindMarkers function, we used FindAllMarkers function to identify differentially expressed genes among each cluster. HSC/MPP signature genes were found using FindMarkers function. Finally, we used FindConservedMarkers function to confirm HSC/MPP signature genes among all conditions. scRNA-seq data were deposited in the NCBI Gene Expression Omnibus^6^ and are accessible through GEO Series accession number GSE248311.

### Gene ontology analysis

Gene ontology analysis was performed using Metascape (https://metascape.org).

### Gene set enrichment analysis

Pre-ranked GSEA^8^ was performed against the MSigDB Hallmark collection (h.all v2026.1)^9^. Genes were ranked by the DESeq2 Wald statistic^10^ for CRCE vs CRCY, calculated separately for CD34+CD38−CD45RA−CD90+ cells and the HSC/MPP cluster, using [number] random cell splits as replicates; p values are therefore descriptive. Enrichment scores used weighted statistics (p = 1), gene sets with fewer than 15 genes in the ranked list were excluded, and significance was assessed with 10,000 gene-set permutations. NES was normalized to the mean of same-sign permuted scores, and FDR was calculated by the Benjamini–Hochberg method across all Hallmark sets tested. Gene sets with FDR ≤ 0.05 are reported.

## Use of Artificial Intelligence

GSEA was performed with the assistance of Claude (Anthropic) under the authors’ supervision and verified independently by the authors using the Broad GSEA desktop application (v4.4.0, 10,000 permutations). Heat maps derived from scRNA-seq data were generated by Claude under the authors’ direction and were manually reviewed and verified by the authors.

## Statistical methods

Statistical analysis was performed using Prism version 10 software. Comparisons of multiple treatment groups were analyzed by one-way ANOVA. Comparisons of two treatment groups were analyzed by 2-tailed Student’s t test. Results were considered significant when P < 0.05. Statistical analysis of limiting dilution assays was performed using ELDA software.

