## Supplemental Data and Methods for "Expansion of human hematopoietic stem cells by inhibiting translation"

### Supplemental Data Files
### Expansion of human hematopoietic stem cells by inhibiting translation

Chenchen Li, Hanna Shin, Dheeraj Bhavanasi, Mai Liu, Maria I. Briones, Xiang Yu, Xiaolei Liu, Friederike A. Herbst, Hooda Said, Mohamad-Gabriel Alameh, Laura Breda, Narasaiah Kovuru, Yeipyeng Kwa, Nithin Sam Ravi, David C. Schultz, Sara Cherry, Stefano Rivella, Juan R. Alvarez-Dominguez, Gerd A. Blobel, Brian D. Gregory, Scott A. Peslak, Jian Huang, Peter S. Klein

**Table of Contents:**

1. Supplemental Figures 1-14 with legends.
2. Supplemental Methods
3. SupplementalTable 1: Differential gene expression in specific populations
4. SupplementalTable 2: GO metascape
5. SupplementalTable 3: HTS data summary
6. Supplemental Table 4: GSEA

Supplemental Figure 1

A

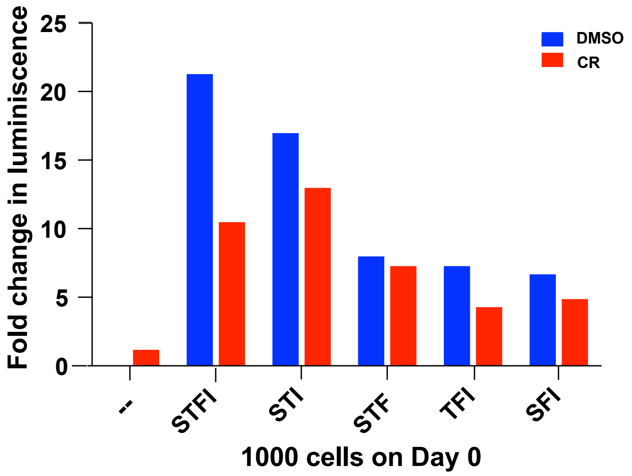


B

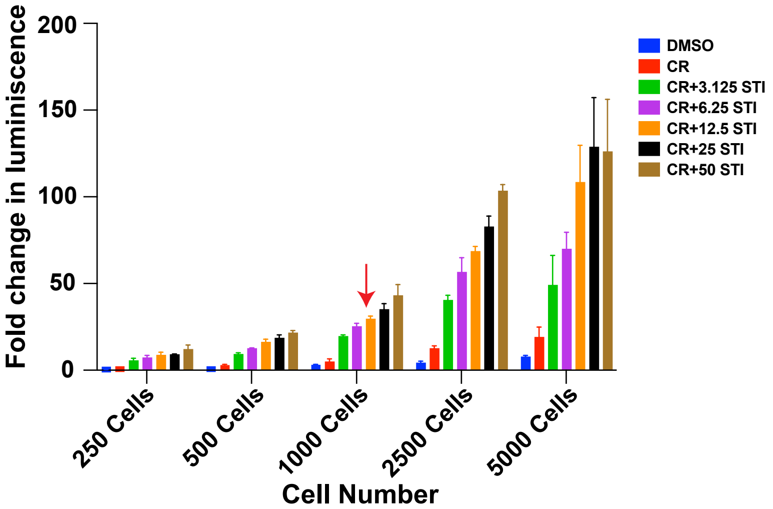


C

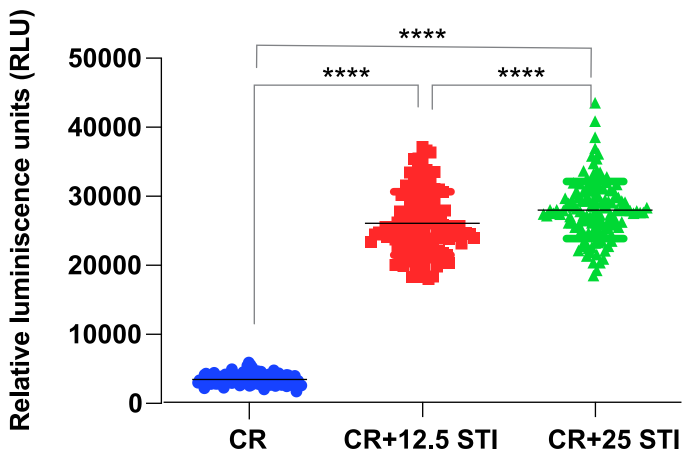


D

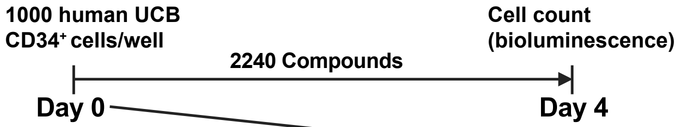


| | 1 | 2 | 3 | 4 | 5 | 6 | 7 | 8 | 9 | 10 | 11 | 12 | 13 | 14 | 15 | 16 | 17 | 18 | 19 | 20 | 21 | 22 | 23 | 24 |
|---|---|---|---|---|---|---|---|---|---|---|---|---|---|---|---|---|---|---|---|---|---|---|---|---|
| A | CR | CRCY | test | test | test | test | test | test | test | test | test | test | test | test | test | test | test | test | test | test | test | test | CRCY | empty |
| B | CR | CRCY | test | test | test | test | test | test | test | test | test | test | test | test | test | test | test | test | test | test | test | test | CRCY | empty |
| C | CR | CRCY | test | test | test | test | test | test | test | test | test | test | test | test | test | test | test | test | test | test | test | test | CRCY | empty |
| D | CR | CRCY | test | test | test | test | test | test | test | test | test | test | test | test | test | test | test | test | test | test | test | test | CRCY | empty |
| E | CR | CRCY | test | test | test | test | test | test | test | test | test | test | test | test | test | test | test | test | test | test | test | test | CRCY | empty |
| F | CR | CRCY | test | test | test | test | test | test | test | test | test | test | test | test | test | test | test | test | test | test | test | test | CRCY | empty |
| G | CR | CRCY | test | test | test | test | test | test | test | test | test | test | test | test | test | test | test | test | test | test | test | test | CRCY | empty |
| H | CR | CRCY | test | test | test | test | test | test | test | test | test | test | test | test | test | test | test | test | test | test | test | test | CRCY | empty |
| I | CR | CRCY | test | test | test | test | test | test | test | test | test | test | test | test | test | test | test | test | test | test | test | test | CRCY | empty |
| J | CR | CRCY | test | test | test | test | test | test | test | test | test | test | test | test | test | test | test | test | test | test | test | test | CRCY | empty |
| K | CR | CRCY | test | test | test | test | test | test | test | test | test | test | test | test | test | test | test | test | test | test | test | test | CRCY | empty |
| L | CR | CRCY | test | test | test | test | test | test | test | test | test | test | test | test | test | test | test | test | test | test | test | test | CRCY | empty |
| M | CR | CRCY | test | test | test | test | test | test | test | test | test | test | test | test | test | test | test | test | test | test | test | test | CRCY | empty |
| N | CR | CRCY | test | test | test | test | test | test | test | test | test | test | test | test | test | test | test | test | test | test | test | test | CRCY | empty |
| O | CR | CRCY | test | test | test | test | test | test | test | test | test | test | test | test | test | test | test | test | test | test | test | test | CRCY | empty |
| P | CR | CRCY | test | test | test | test | test | test | test | test | test | test | test | test | test | test | test | test | test | test | test | test | CRCY | empty |

**Supplemental Figure 1. Optimization of screening conditions. A.** Human UCB $CD34^+$ cells were added to 384 well dishes in StemSpan SFEM with (red) or without (blue) CR and varying combinations of cytokines as indicated. S: SCF (100 ng/ml), T; Thrombopoietin (100 ng/ml), F: Flt3 ligand (100 ng/ml), I: IL3 (10 ng/ml). Cells were cultured for 4 days and cell number measured using a bioluminescence assay (ATPlite). Relative light units (RLU) are shown. Red arrow indicates the combination (STI) selected for the screen. **B.** Varying numbers of $CD34^+$ cells/well and varying concentrations of STI were added to a 384-well dish; cells were cultured for 4 days and cell number measured by bioluminescent assay. Red arrow indicates optimal cell number (1000 cells) and concentration of STI (12.5 ng/ml SCF, 12.5 ng/ml TPO, and 1.25 ng/ml IL3; subsequently referred to as CRCY) selected for HTS. **C.** Validation of screening conditions with CR vs CRCY at two concentrations. **D.** Screen was performed with 1000 cells/well in CRCY in 384 well dishes with 32 control wells (CRCY+DMSO) per plate (lanes 2 and 23) and Selleck library compounds (test) in lanes 3-22. Cells were cultured for 4 days without medium change and cell number was assessed by ATP bioluminescent assay.

#### Supplemental Figure 2

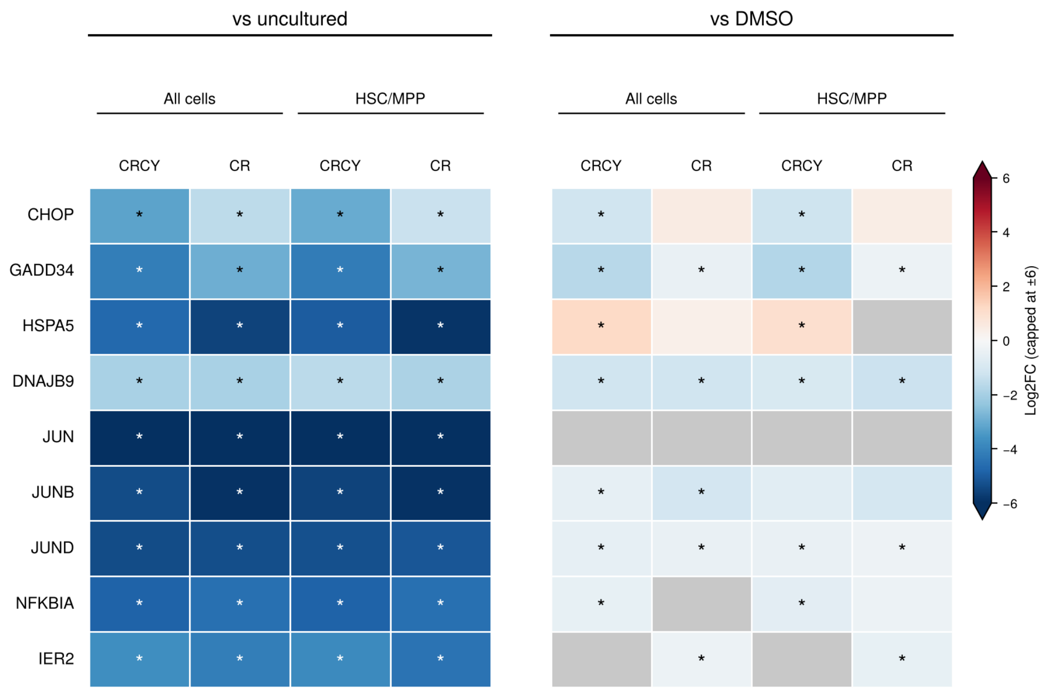


**Supplemental Figure 2: scRNA-seq analysis of UPR markers and immediate early response genes (IEGs).** As in Figure 1, HSC enriched $CD34^+CD38^-CD45RA^-CD90^+$ cells sorted from CD34+ UBC were cultured in basal medium (Stem-span) with vehicle (DMSO), CR, or CRCY for 2 days and then subjected to 10X Genomics scRNA-seq along with freshly thawed cells from the same donor (day 0 = uncultured). Log2 fold change for CRCY or CR vs uncultured and CRCY or CR vs basal medium with DMSO (DMSO) in the $CD34^+CD38^-CD45RA^-CD90^+$ population (all cells) or in the HSC/MPP cluster is shown with values truncated at ±6. An asterisk indicates adjusted $p \leq 0.05$, although all but one cell that achieved significance had $p < 0.001$. Gray indicates nonadjusted $p > 0.05$ or not in table because gene was below FindMarkers thresholds ($|\log 2FC| < 0.25$ or expressed in < 10% of cells). The data show marked reduction in UPR markers and IEGs compared to uncultured cells, consistent with known elevation of stress markers in freeze/thawed and FACS sorted cells. Most UPR markers and IEGs are reduced in CR, CRCY, or CRCE when compared to cells cultured in basal medium+vehicle (DMSO), although the low number of cells in the DMSO group limited statistical power.

#### Supplemental Figure 3

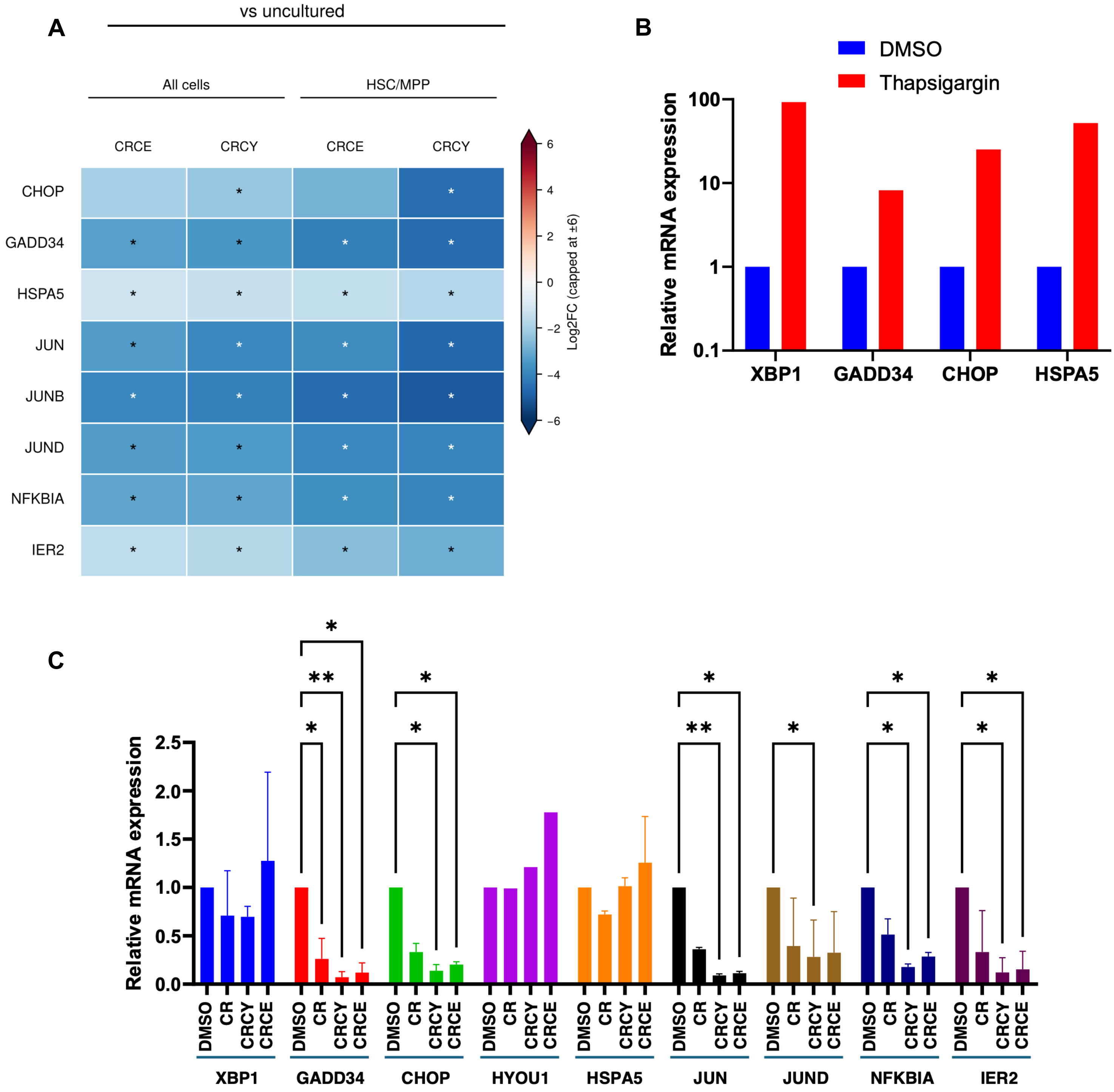


**Supplemental Figure 3: Expression of UPR markers and immediate early response genes (IEGs).**
A. scRNA-seq: As in Figure 1 and 3, HSC enriched CD34+CD38-CD45RA-CD90+ cells sorted from CD34+ UBC were cultured in basal medium with vehicle (DMSO), CRCY, or CRCE for 2 days and then subjected to 10X Genomics scRNA-seq along with day 0 (uncultured) cells from the same donor. Log2FC is shown. An asterisk indicates adjusted $p \leq 0.05$ (although most p values were $< 0.001$). As in Supplemental Figure 2, gray indicates nonadjusted $p > 0.05$ or below FindMarkers thresholds ($|\log2FC| < 0.25$ or expressed in < 10% of cells). These data confirm marked reduction in UPR markers and IEGs compared to uncultured cells, consistent with known elevation of stress markers in freeze/thawed and FACS sorted cells. Most UPR markers

and IEGs are also reduced when compared to cells cultured for 2 days in basal medium+vehicle (DMSO), although the low number of cells in the DMSO group limited statistical power and detection of several markers. **B.** Positive control for UPR markers: A group of $CD34^+$ cells was cultured for 22 hours in basal medium+vehicle (DMSO) and treated with 300nM thapsigargin for 2 hours prior to collection at 24h. XBP1 splicing and expression of CHOP, GADD34, and HSPA5 were assessed by RT-qPCR. As expected, thapsigargin activated IRE1/XBP1, PERK/ATF4, and ATF6 pathways. **C.** CD34+ cells at day 0 (uncultured) or cultured for 24 hours in basal medium (Stem-span) with vehicle (DMSO), CR, CRCY, or CRCE were collected for RT-qPCR. IRE1/XBP1 activation was assessed by RT-qPCR for the spliced form of XBP1. PERK/ATF4 pathway was assessed by expression of CHOP (DDIT3) and GADD34 (PPP1R15A). ATF6/XBP1 regulate HSPA5. Immediate early genes JUN, JUND, NFKBIA, and IER2 were also measured by RT-qPCR. Data are from 2 independent UCB donors. Only values with $p < 0.05$ are marked: * indicates $p<0.05$, ** $p<0.01$ (two way ANOVA). Compared to vehicle (DMSO) control, CHOP and GADD34 were significantly reduced in CR compared to vehicle control, and further reduced in CRCY and CRCE, while XBP1 and HSPA5 were not significantly affected. All of the IEGs were also significantly reduced in CR, CRCY, and CRCE. UPR and IEG markers were markedly reduced compared to day 0/uncultured cells (not shown), likely representing stress associated with freeze/thawing and initiation of culture.

**Supplemental Figure 4**

**Gene-level changes in MYC (V1) and OXPHOS programs in CRCE vs CRCY**

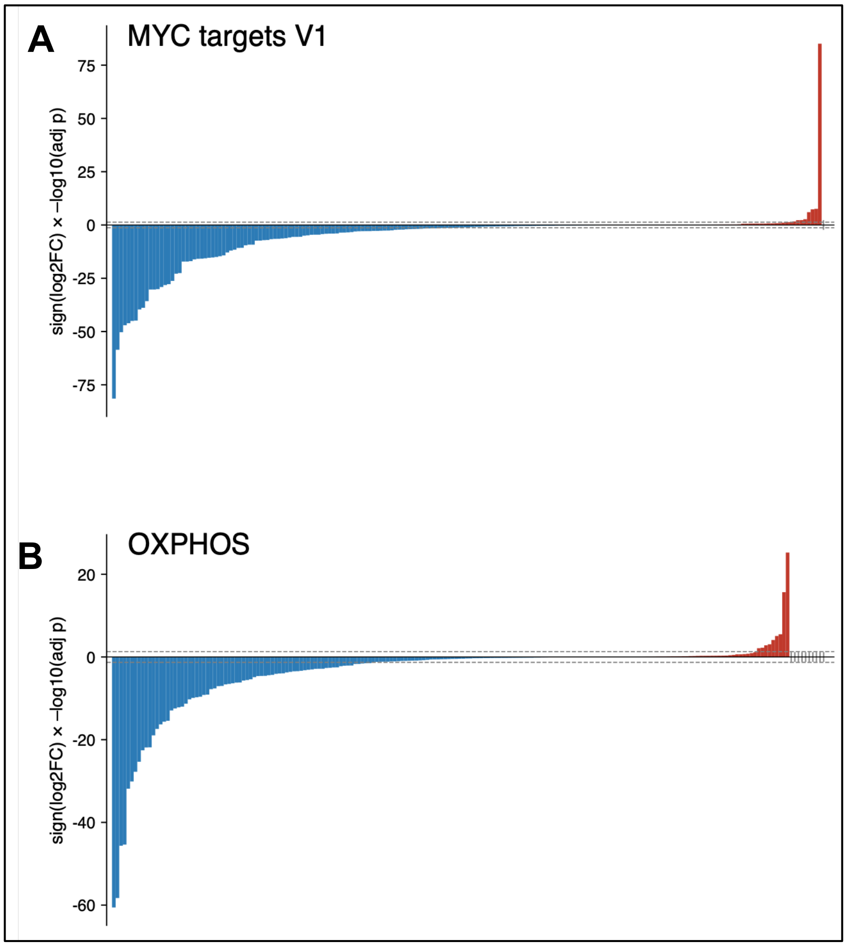


**Supplemental Figure 4. Gene-level changes in MYC and oxidative phosphorylation (OXPHOS) gene programs between CRCE and CRCY.** Each bar represents one gene of the Hallmark MYC targets V1 gene set (A, 195 genes) or the Hallmark OXPHOS gene set (B, 198 genes) in CD34+CD38−CD45RA−CD90+ cells (all cells). The height of each bar is the sign of the log2 fold change multiplied by −log10 of the DESeq2 adjusted p value for CRCE vs CRCY, and genes are ordered from most negative to most positive. Blue bars are genes lower in CRCE, and red bars are genes higher in CRCE. Dashed lines mark adjusted p = 0.05. 148 of 195 MYC target genes were lower in CRCE, with 94 significantly lower and 10 significantly higher; 140 of 198 OXPHOS genes were lower in CRCE, with 73 significantly lower and 9 significantly higher. DESeq2 replicates were random splits of cells, so p values are descriptive.

#### Supplemental Figure 5

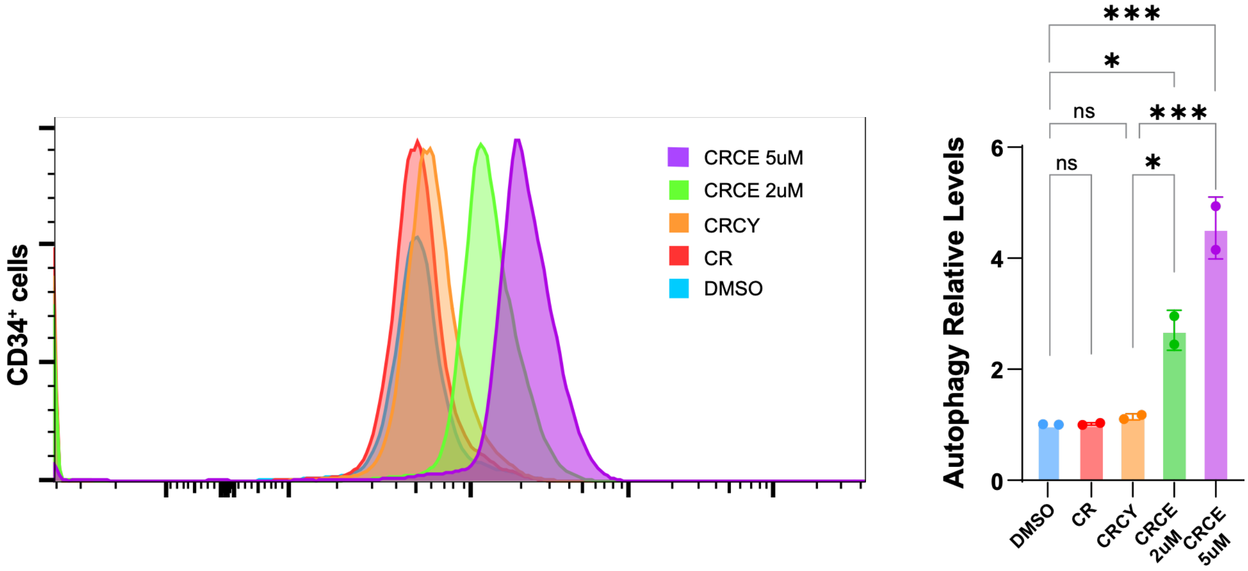


**Supplemental Figure 5: Does-dependent increase in autophagic vacuole content with 4E1RCat.** CD34+ cells from UCB were cultured in control medium (DMSO), CR, CRCY, CRCE (CRCY + 2μM 4E1RCat), or CRCY + 5 μM 4E1RCat for 24 hours and autophagic vacuole content was measured by flow cytometry as in Figure 3E. A. Flow plot from one donor is representative of replicates from 3 donors. B. MFI reflecting autophagic vacuole content. 4E1Rcat increased autophagic vacuoles at 2 μM and this further increased at 5 μM. * indicates $p < 0.05$, *** indicates $p < 0.005$. Data are from one donor; a similar dose-dependent increase was observed in 2 other donors.

#### Supplemental Figure 6

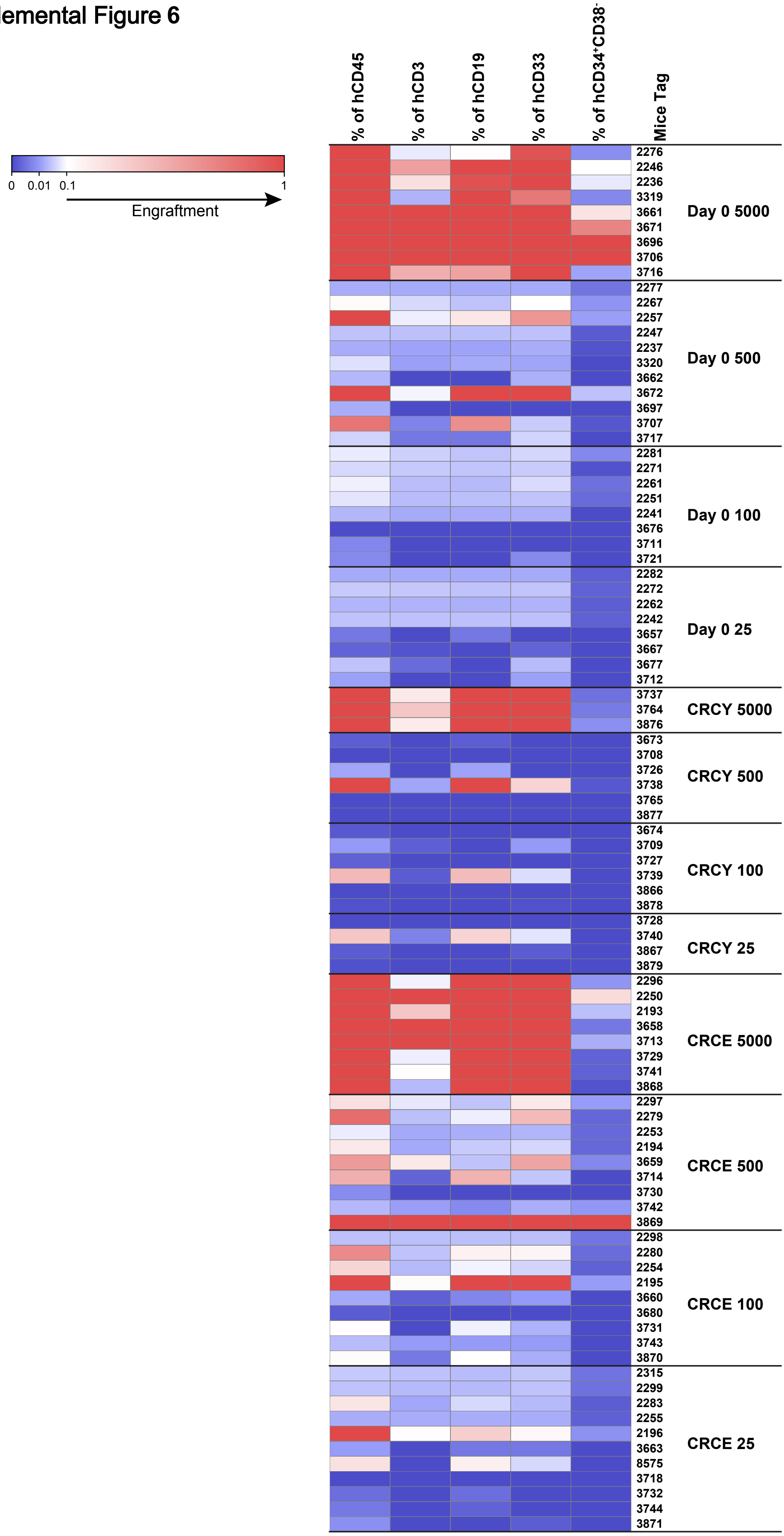

**Supplemental Figure 6. Multilineage Contribution of HSCs expanded in CRCE in primary LDA at 20 weeks.** The percentage of human CD45$^+$ cells, T cells (CD3$^+$), B cells (CD19$^+$), myeloid cells (CD33$^+$), and CD34$^+$CD38$^-$ cells in each primary recipient mouse at 20 weeks in bone marrow are represented by a heatmap. Red indicates percentage engraftment (human cells) is > 0.1%. Blue indicates human cell engraftment < 0.1%. White indicates percentage engraftment = 0.1%. Data show results from expansion/transplants using 2 distinct mixed donor UCB samples.

**Supplemental Figure 7**

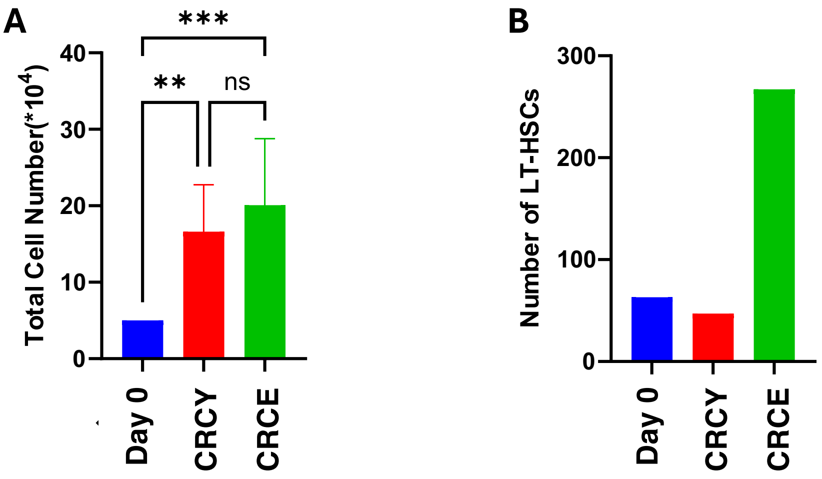


**Supplemental Figure 7: Number of $CD34^{+}$ cells and LT-HSCs (SRCs) at Day 0 and after 7 days**. A. total cell count at Day 0 (50,000 for all groups) or after 7 days of culture in CRCY or CRCE. B. Total number of LT-HSCs is shown based on frequency calculated by ELDA x cell number at day 0. The number of LT-HSCs was 63 at day 0, 47 at day 7 in CRCY, and 267 at day 7 in CRCE. Addition of 4E1RCat therefore expanded LT-HSCs 5.7-fold compared to CRCE and 4.3-fold compared to day 0.

Supplemental Figure 8

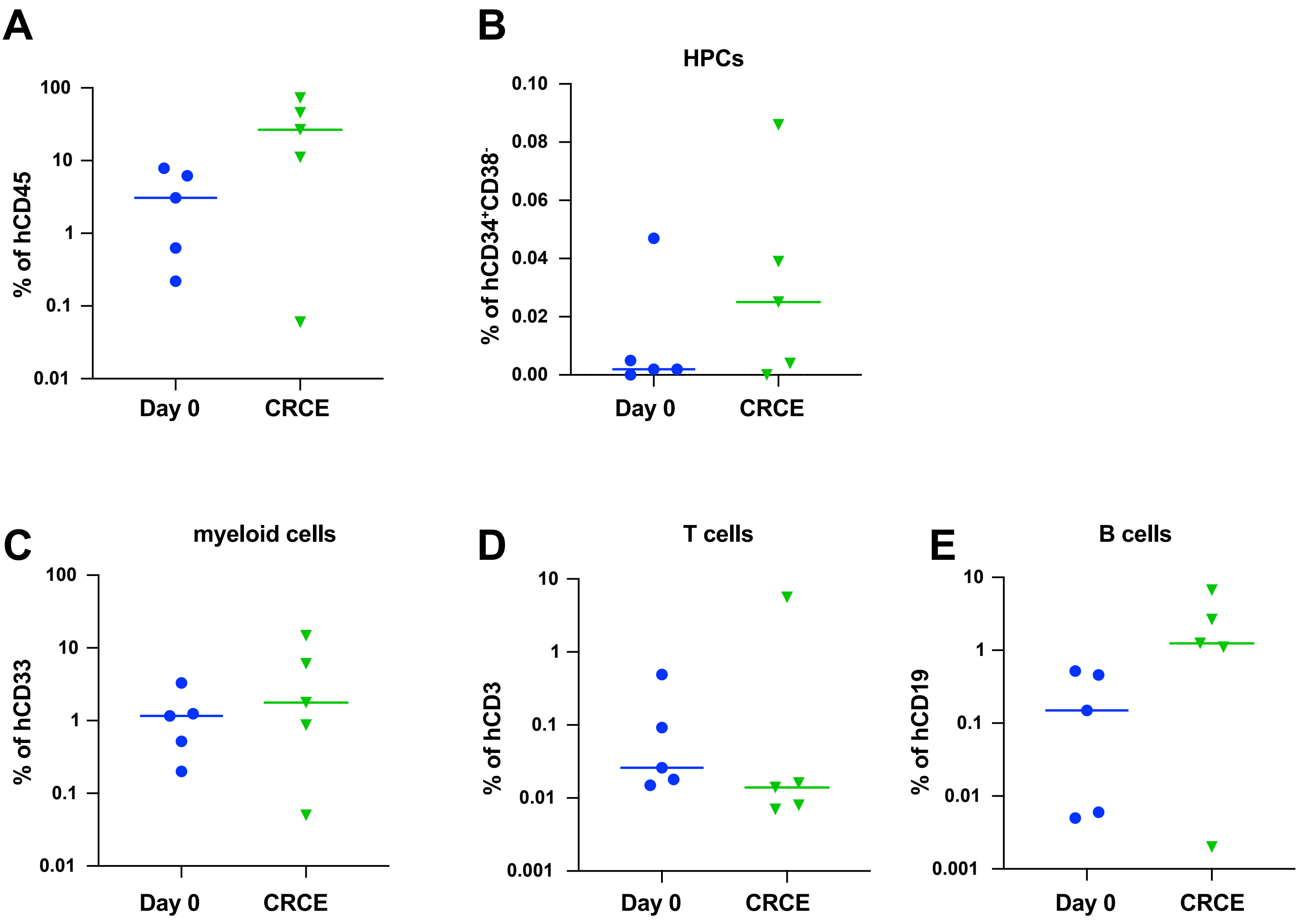


F

| Group | Donor Tag # | % of hCD45 | % of hCD19 | % of hCD33 | % of hCD3 | % of hCD34$^+$CD38$^-$ |
|---|---|---|---|---|---|---|
| Day 0 | 2276 | 7.87 | 0.52 | 3.3 | 0.026 | 0.047 |
| | 2246 | 6.19 | 0.46 | 1.24 | 0.49 | 0.00455 |
| | 2236 | 3.08 | 0.15 | 1.16 | 0.018 | 0.00161 |
| | 2257 | 0.63 | 0.00501 | 0.52 | 0.015 | 0.00167 |
| | 2267 | 0.22 | 0.00585 | 0.2 | 0.092 | 0 |
| CRCE | 2296 | 11.1 | 1.1 | 0.87 | 0.014 | 0.00434 |
| | 2250 | 45.6 | 1.24 | 14.7 | 5.59 | 0.039 |
| | 2193 | 72.5 | 6.68 | 6.07 | 0.00744 | 0.086 |
| | 2195 | 26.6 | 2.64 | 1.77 | 0.0077 | 0.025 |
| | 2196 | 0.06 | 0.00162 | 0.05 | 0.016 | 0 |

**Supplemental Figure 8. Multilineage Contribution of HSCs expanded in CRCE in primary recipients at 29 weeks.A-E.** The percentage of human $CD45^+$ cells, $CD34^+CD38^-$ cells, myeloid cells ($CD33^+$), T cells ($CD3^+$), and B cells ($CD19^+$) at day 29 post-transplant in bone marrow of mice used as donors for secondary transplant. Mice had received either uncultured (Day 0) $CD34^+$ cells or cells cultured in CRCE for 7 days. There was no significant difference between Day 0 and CRCE/Day 7 groups in panels A-E. **F.** Table showing percent engraftment of each population in each mouse at 29 weeks.

#### Supplemental Figure 9

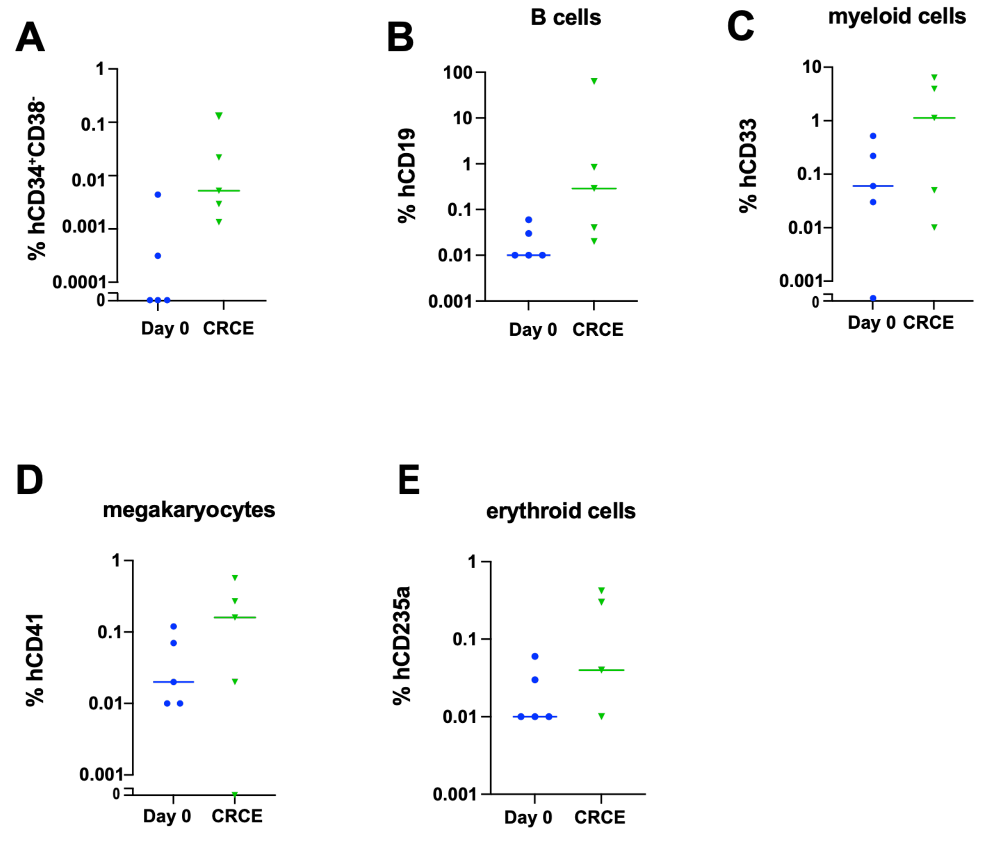


**F**

| Group | Donor Tag # | Recipient Tag# | % of hCD45 (PB) | % of hCD45 (BM) | % of hCD3 (BM) | % of hCD19 (BM) |
|---|---|---|---|---|---|---|
| Day 0 | 2276 | 2410 | 0 | 0.64 | 0.035 | 0.056 |
| | 2246 | 2409 | 0 | 0.29 | 0.00442 | 0.028 |
| | 2236 | 2407 | 0.00645 | 0.075 | 0.00243 | 0.00729 |
| | 2257 | 2404 | 0.016 | 0.04 | 0.00171 | 0.00854 |
| | 2267 | 2403 | 0.00854 | 0.016 | 0.00913 | 0.00913 |
| CRCE | 2296 | 2419 | 0 | 0.074 | 0.00586 | 0.021 |
| | 2250 | 2418 | 0.32 | 5.24 | 1.71 | 0.29 |
| | 2193 | 2413 | 11.3 | 82 | 0.23 | 62.7 |
| | 2195 | 2414 | 0.77 | 5.49 | 0.099 | 0.84 |
| | 2196 | 2416 | 0 | 0.064 | 0.013 | 0.04 |

| Group | Donor Tag # | Recipient Tag# | % of hCD33 (BM) | % of hCD41 (BM) | % of hGlyA (BM) | % of $hCD34^{+}CD38^{-}$ (BM) |
|---|---|---|---|---|---|---|
| Day 0 | 2276 | 2410 | 0.52 | 0.12 | 0.06 | 0.0044 |
| | 2246 | 2409 | 0.22 | 0.069 | 0.027 | 0.000316 |
| | 2236 | 2407 | 0.062 | 0.021 | 0.011 | 0 |
| | 2257 | 2404 | 0.031 | 0.011 | 0.00854 | 0 |
| | 2267 | 2403 | 0.00269 | 0.00591 | 0.00591 | 0 |
| CRCE | 2296 | 2419 | 0.051 | 0.018 | 0.0088 | 0.00293 |
| | 2250 | 2418 | 1.13 | 0.16 | 0.3 | 0.022 |
| | 2193 | 2413 | 6.38 | 0.57 | 0.42 | 0.13 |
| | 2195 | 2414 | 3.94 | 0.27 | 0.042 | 0.00521 |
| | 2196 | 2416 | 0.00935 | 0.00267 | 0.00401 | 0.00134 |

**Supplemental Figure 9. Lineage analysis of serial transplant of LT-HSCs expanded with CRCE. See Figure 4G-I.** Bone marrow harvested from primary recipients at 29 weeks was transplanted into conditioned secondary recipients and harvested at 18 weeks as described in Figure 4G-I. The percentage ($log_{10}$ scale) of human **A.** lineage negative cells ($CD34^{+}CD38^{-}$), **B.** B cells ($CD19^{+}$), **C.** myeloid cells ($CD33^{+}$), **D.** megakaryocytes ($CD41^{+}$), and **E.** erythroid cells ($CD235a^{+}$) in the bone marrow (18 weeks) of secondary recipients. Samples with 0% engraftment are shown at the baseline (with broken y-axes). **F.** Table showing per mouse percentage of human $CD45^{+}$ cells in peripheral blood (PB) at 16 weeks and in bone marrow (BM) at 18 weeks. The percentage of human $CD45^{+}$ T cells ($CD3^{+}$), B cells ($CD19^{+}$), myeloid cells ($CD33^{+}$), megakaryocytic cells ($CD41^{+}$), erythroid cells ($GlyA^{+}$) and $CD34^{+}CD38^{-}$ cells is also shown in the table .

#### Supplemental Figure 10

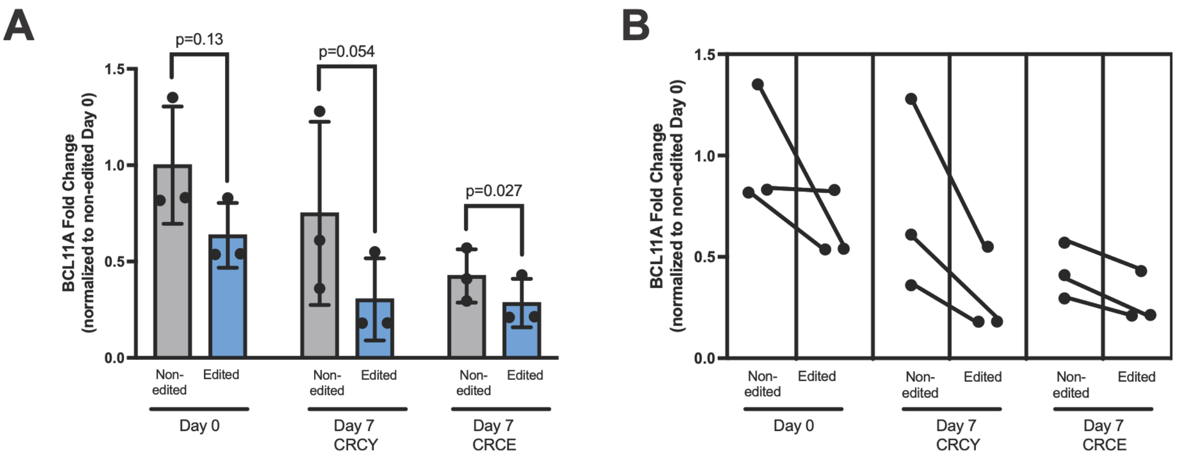


**Supplemental Figure 10: Pairwise comparison of *BCL11A*+58 enhancer editing in healthy donor-derived CD34$^{+}$ cells expanded ex vivo**. BCL11A expression data from Figure 6E was compared pairwise for samples with matched edited and non-edited data in uncultured (Day 0) cells or cells cultured for 7 says in CRCY or CRCE. A. The reduction of *BCL11A* transcript levels in *BCL11A*+58-edited samples compared to unedited samples was statistically significant in the CRCE group ($p$ = 0.027) and approached significance in the CRCY group ($p$ = 0.054; one-tailed paired t-test). B. Paired dot plot identifying *BCL11A* transcript levels for each pair of samples in panel A (unedited vs *BCL11A*+58 edited). *BCL11A* transcript levels were normalized to Day 0 non-edited sample for each donor; N = 3 independent donors.

### Supplemental Figure 11

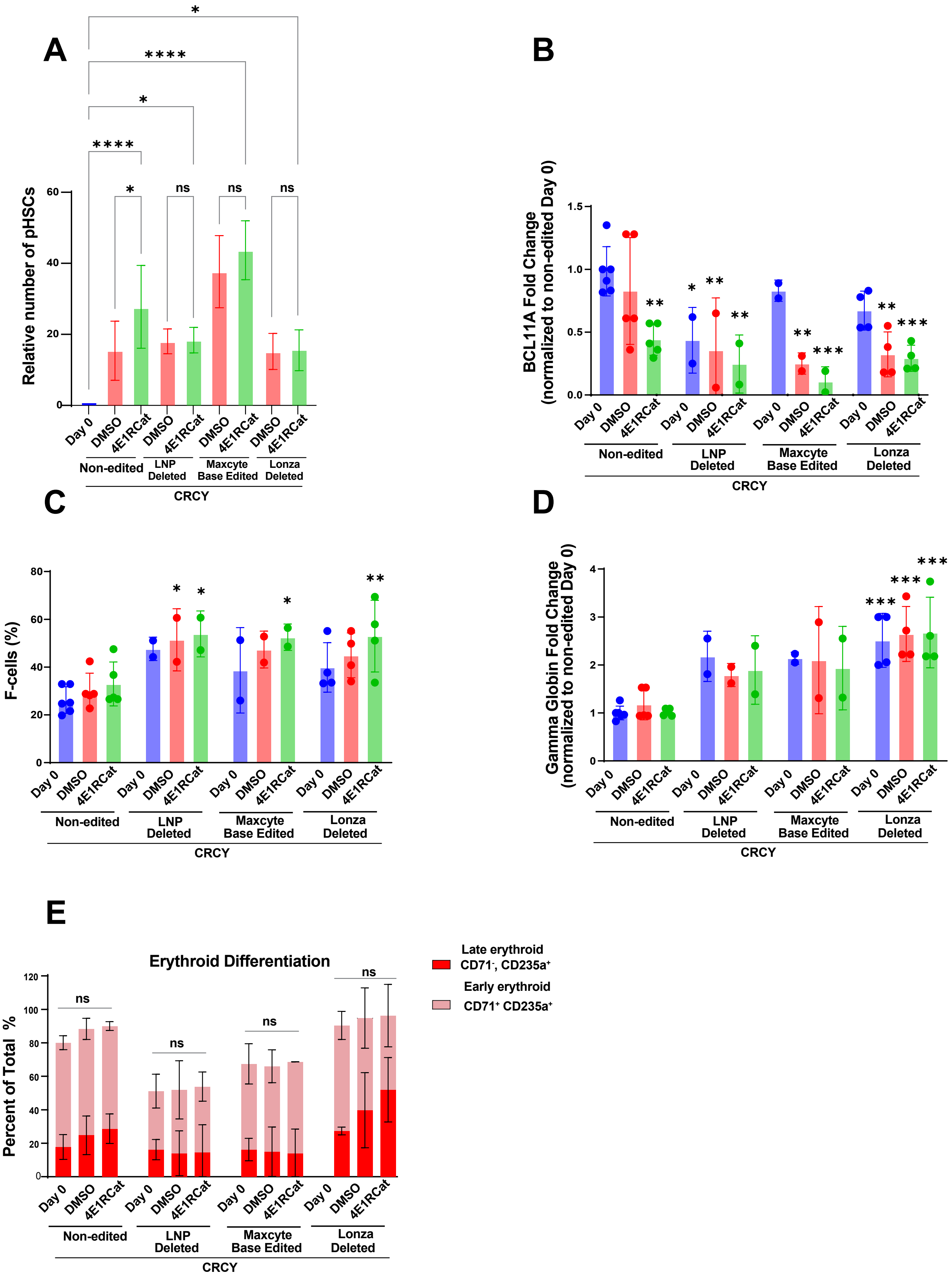

**Supplemental Figure 11. Expansion of adult HSCs after editing with conventional CRISPR/Cas9 (Lonza-EP), base editing (CBE-EP), or LNPtransduction.** Mobilized adult $CD34^+$ cells were edited by CRISPR/Cas9 targeting of the *BCL11A+58* enhancer to compare conventional CRISPR/Cas9 using electroporation (Lonza-EP), as in Figure 4, to transduction of Cas9/gRNA RNPs with lipid nanoparticles (LNP) and to transduction of aTadA-based cytosine base editor using MaxCyte EP(CBE-EP)After transduction, cells were cultured in CRCYwith vehicle (indicated as "DMSO") or CRCYwith 4E1RCat (indicated as "4E1RCat") as in Figure 4. **A.** The number of pHSCs ($CD34^+CD38^-CD45RA^-CD90^+CD49f^+$) at Day 7 relative to Day 0 is shown. Expansion of edited pHSCs (compared to day 0) is similar to unedited cells, and therefore not impaired by any of the above editing approaches. The data for non-edited and Lonza-EP represent the mean of replicates from 4-6 adult donors; the data for Maxcyte and LNP are from 2 adult donors. * indicates $p < 0.05$, ** indicates $p < 0.01$, *** indicates $p < 0.001$, **** indicates $p < 0.0001$. NS, not significant (one-wayANOVA)**. B-E.** $CD34^+$ cells from Day 0 (unedited), immediately after transduction (electroporation or exposure to LNPs) with *BCL11A +58* sgRNA(Day 0 edited), or after transduction and culture in CRCY with or without 4E1RCat (Day 7 edited), were differentiated in three-phase erythroid culture and collected at Day 13 for detection of **(B)** *BCL11A* mRNAby RT-qPCR (normalized to Day 0 non-edited), demonstrating reduction in *BCL11A* expression in all edited groups cultured in either CRCY or CRCY+4E1RCat compared to unedited, day 0 cells; (**C)** HbF expression (F-cells) by flow cytometry (as percent of CD71+CD235a+ erythroid cells), showing an increase in % F-cells in cells cultured in CRCY +4E1RCat after editing by Lonza-EP, CBE-EP, or LNP. Importantly, culture in CRCY±4E1RCat did not impair induction of F-cells; (**D**) *HBG* mRNAexpression by RT-qPCR (normalized to Day 0 non-edited), again showing that culture in CRCY or CRCY+4E1RCat does not impair induction of HBG. For panels B-D, * indicates p < 0.05, ** indicates p < 0.01, and *** indicates p < 0.001 as compared to Day 0 non-edited control (one-way ANOVA). (**E**) Erythroid differentiation by flow cytometry (as percent of total cells) is not impaired by culture in CRCY or CRCY+4E1RCat. NS indicates not significant within each editing condition group (two-way ANOVA).

#### Supplemental Figure 12

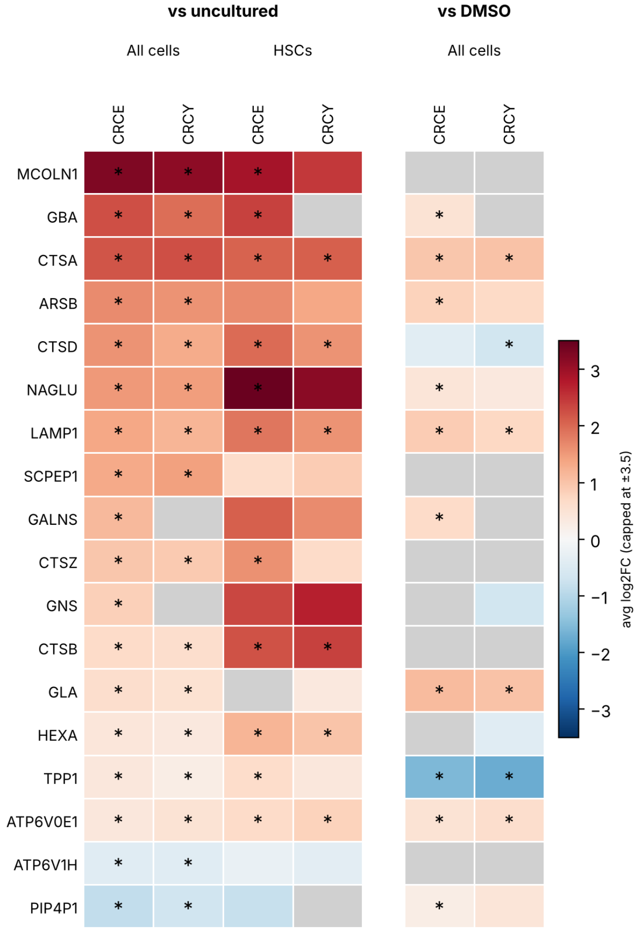


**Supplemental Figure 12: CLEAR Network genes in HSC-enriched cells and HSC/MPP cluster after culture in CRCE or CRCY:** scRNA-seq analysis of TFEB target genes associated with lysosomal biogenesis and function (Taken from Sardiello et al Science 2009 PMID 19556463). HSC-enriched $CD34^{+}CD38^{-}CD45RA^{-}CD90^{+}$ cells sorted from CD34+ UBC were cultured in CRCY or CRCE for 2 days and then subjected to scRNA-seq along with freshly thawed and FACS sorted cells from the same donor (day 0 = uncultured), described in detail in Figure 3 and Supplemental Figure 3. Log2[fold change] (Log2FC) for CRCY or CRCE vs uncultured in the $CD34^{+}CD38^{-}CD45RA^{-}CD90^{+}$ population (all cells) or in the HSC/MPP cluster is shown with values truncated at ±3. Log2FC vs basal medium+vehicle (DMSO) for the $CD34^{+}CD38^{-}CD45RA^{-}CD90^{+}$ population is also shown. An asterisk indicates adjusted $p \leq 0.05$. Gray indicates nonadjusted $p > 0.05$ or not in present in the table because it was below FindMarkers thresholds ($|log2FC| < 0.25$ or expressed in < 10% of cells). The data show an increase in most of the listed CLEAR network genes compared to uncultured cells in both the total HSC-enriched population (all cells) and the HSC/MPP cluster. A similar trend was observed for a subset of these genes when comparing CRCE and CRCY to cells cultured in basal medium+vehicle (DMSO), but the low number of cells in the DMSO group and in the HSC/MPP cluster limited statistical power.

**Supplemental Figure 13**

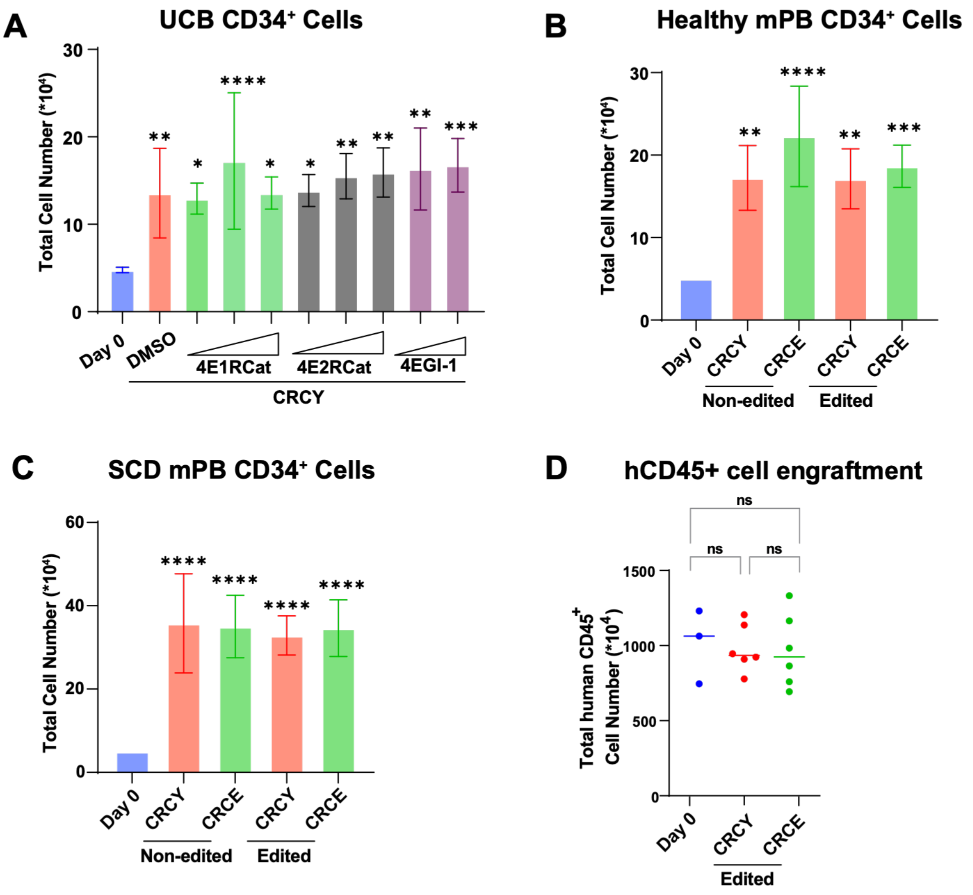


**Supplemental Figure 13: Total CD34$^+$ cell counts before (Day 0) and after (Day 7) culture in CRCY and CRCE.** A. From Figure 2, expansion in CRCY with vehicle (DMSO) or increasing doses of 4E1RCat, 4E2RCat, or 4EGI-1. B, C. From Figure 6, mobilized peripheral blood (mPB) CD34$^+$ cells at Day 0 or day 7 culture of non-edited vs edited cells cultured in CRCY or CRCE. D. From Figure 7, absolute number of engrafted human CD45+ cells. Mean for Day 0 = 1.01 x $10^7$ hCD45$^+$ cells/mouse; day 7 CRCY = 0.98 x $10^7$ hCD45$^+$ cells/mouse; day 7 CRCE = 0.97 x $10^7$ hCD45$^+$ cells/mouse. (* indicates $p < 0.05$; ** $p < 0.01$, *** $p < 0.005$, **** $p < 0.001$; “ns”: not significant; for all panels: one-way ANOVA).

**Supplemental Figure 14 Gating Strategies**

**A. Representative Phenotypic Hematopoietic Stem Cell (pHSC) Flow Gating**

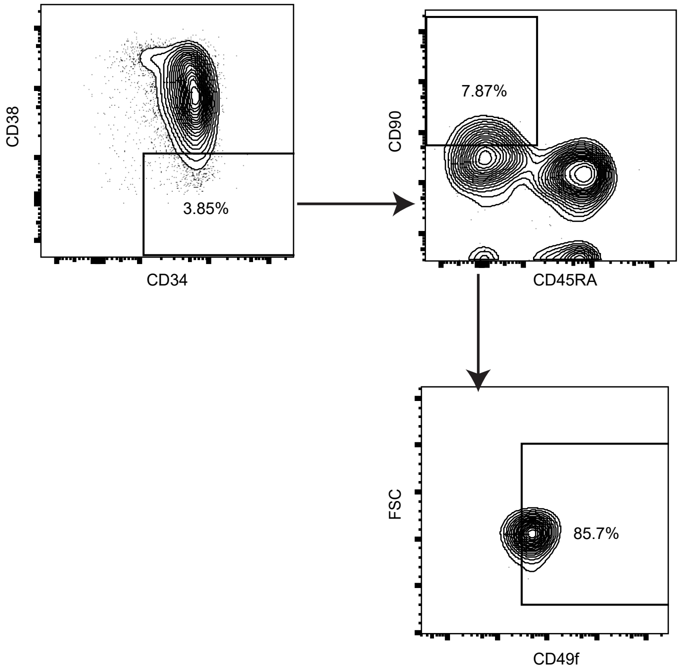


**B. Representative Human Engraftment Flow Gating**

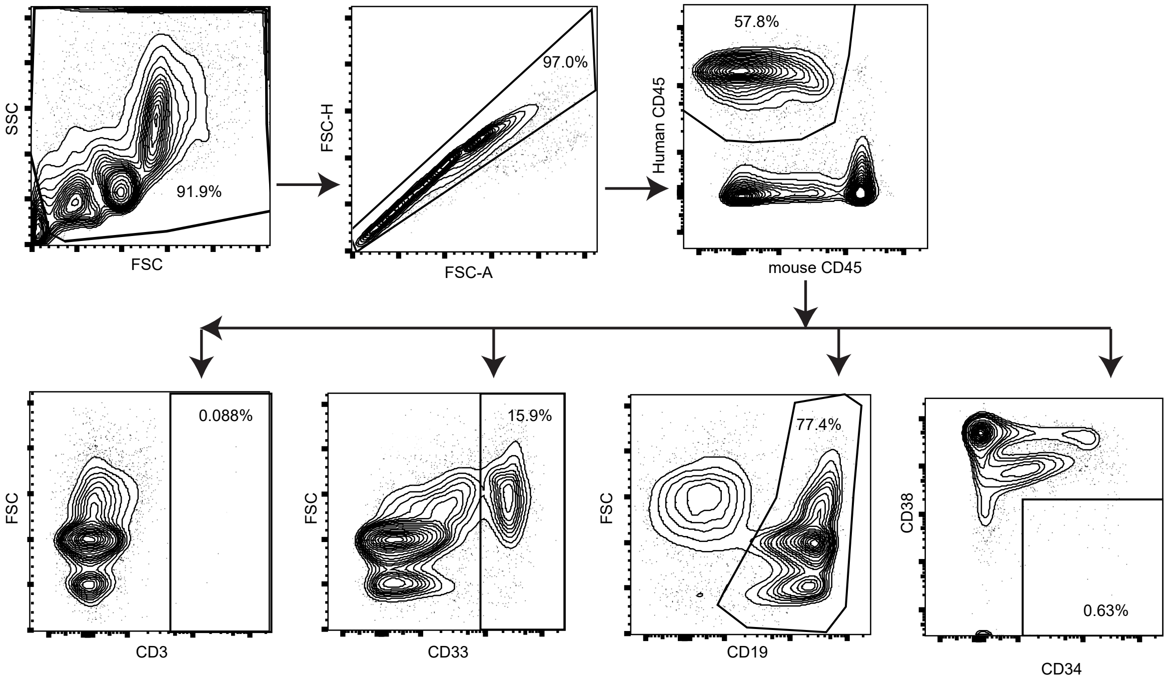

## Supplemental Figure 14    Gating Strategies

##### C. Representative HbF Flow Gating

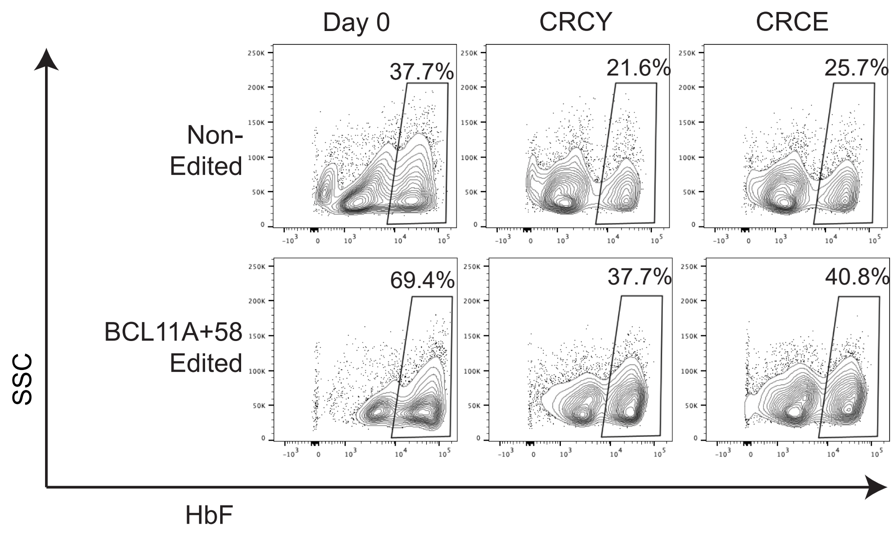


#### D. Representative CD34+ Erythroid Differentiation Flow Gating

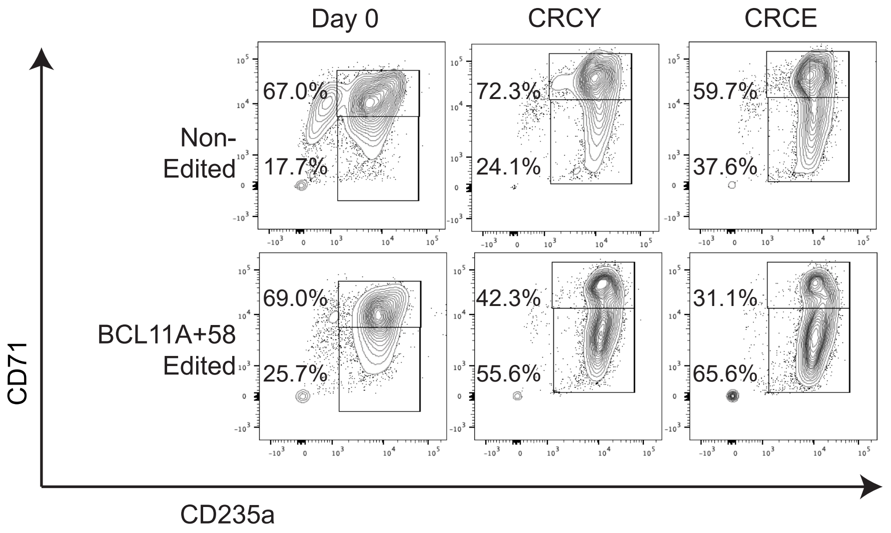

#### Supplemental Methods

##### Antibodies for flow cytometry

| Antibody | Company | Catalog # |
|---|---|---|
| PE anti-human CD45 | Biolegend | 368510 |
| APC/Cyanine7 anti-mouse CD45 | Biolegend | 103116 |
| APC anti-human CD3 | Biolegend | 317318 |
| PerCP/Cyanine5.5 anti-human CD19 | Biolegend | 302230 |
| BB700 mouse anti-Human CD19 | BD Biosciences | 566396 |
| FITC anti-human CD33 | Biolegend | 366620 |
| BB515 Mouse anti-Human CD33 | BD Biosciences | 564588 |
| Brilliant Violet 421™ anti-human CD41 | Biolegend | 303730 |
| BUV395 Mouse anti-Human CD235a | BD Biosciences | 563810 |
| Alexa Fluor 700 anti-human CD34 | Biolegend | 343622 |
| PE/Cyanine7 anti-human CD34 | Biolegend | 343616 |
| PE/Cyanine7 anti-human CD38 | Biolegend | 397114 |
| Brilliant Violet 711™ anti-human CD38 | Biolegend | 303528 |
| Brilliant Violet 421™ anti-human CD38 | Biolegend | 356618 |
| APC/Cyanine7 anti-human CD45RA | Biolegend | 304128 |
| BUV395 anti-human CD90 | BD Biosciences | 563804 |
| Alexa Fluor 488 anti-human CD49f | Biolegend | 313608 |
| Fetal Hemoglobin | Novus Biologicals | NB110-41084 |
| PE anti-human CD71 | Biolegend | 334106 |
| PE/Cyanine7 anti-human CD235ab (HbF analysis) | Biolegend | 306620 |
| FITC Mouse anti-human CD235a (engraftment) | Biolegend | 349104 |
| Alexa Fluor 488 anti-GRP78 mAb (C38) | Invitrogen | 53-9768-82 |
